# SpacePath enables pathway and molecular-family analysis beyond annotation ambiguity in spatial metabolomics and lipidomics

**DOI:** 10.64898/2026.10.16.759546

**Authors:** Johanna Galvis, Mélanie Moreau, Sébastian Lillo, Doriane Bomont, Léa Ledoux, Yanis Zirem, Michel Salzet, Isabelle Fournier, Nicolas Desbenoit, Thomas Daubon, Macha Nikolski

## Abstract

Mass spectrometry imaging (MSI) maps metabolites and lipids in tissue with spatial resolution, but molecular identity remains partly resolved: single ions match multiple candidate compounds, and most detected metabolites and lipids fall outside canonical biochemical pathways. We developed SpacePath, a computational framework that retains ion-to-compound ambiguity, scores canonical pathways across tissue locations, and groups assignments that were unmapped to pathways into *de novo* molecular families by chemical similarity. Pathway mappings covered on average only 1% of molecular assignments, whereas the combined SpacePath representation covered 24% of pooled assignments. Benchmarking on four public spatial lipidomics datasets (healthy mouse brain, cerebellum, a demyelination model, and human bladder cancer) recovered previously reported spatial patterns. In newly acquired glioblastoma xenograft datasets, SpacePath identified lipid programs specific to tumor and surrounding brain, and SpiderMass-MS/MS independently confirmed four tumor-associated features from a *de novo* family, refining the initial MS1 assignments. SpacePath extends the spatial interpretation of MSI data beyond canonical pathways while preserving annotation uncertainty.

## 1 Main

Mass spectrometry imaging (MSI) enables the spatial mapping of hundreds to thousands of metabolites and lipids directly in tissue sections, providing a molecular view of tissue organization that complements spatial transcriptomic and proteomic technologies [1, 2]. This is particularly relevant in heterogeneous tissues such as tumors due to high variability in metabolic states at both cell and tissue levels [3]. Glioblastoma (GB), for example, shows strong metabolic heterogeneity and adapts to the lipid-rich environment of myelinated brain regions [4–10], making spatial metabolomics and lipidomics a relevant technique for resolving tumor-associated metabolic niches.

However, functional interpretation of MSI data remains difficult. While MSI measurements provide spatially resolved mass-to-charge (*m/z*) values, compound identification from MS1 information alone is often ambiguous. Unlike liquid chromatography–mass spectrometry, MSI does not provide chromatographic separation, and molecular annotation is commonly based on MS1 information. Consequently, a single *m/z* feature can correspond to multiple candidate molecular identities [11]. Annotation tools such as Metaspace [12] score matches of ions to formula, but downstream analyses must still accommodate multiple plausible compound identities for the same ion. Thus, spatial structure can be measured at high resolution while molecular identity remains only partially resolved, complicating any attempt to move from individual ion maps to higher level metabolic interpretation.

Pathway level analysis could provide such higher level interpretation of MSI data by summarizing groups of molecular features into a pathway score for each spatial location. However, most spatial pathway scoring methods have been developed for transcriptomic data [13–16], where well-defined gene identities can be easily mapped to pathways. Existing MSI workflows mostly focus on image segmentation, feature level statistics or multimodal integration [17–22]. One exception is SPATA2, which integrates spatial transcriptomic and MALDI metabolomic data [23] and combines transcriptome-based scoring with pathway enrichment of spatial modules defined from metabolomic information. However, the framework was not designed to compute pathway scores directly from MSI molecular assignments. A dedicated MSI framework is needed to account for the uncertainty in molecular assignments when computing spot-wise scores and to retain spatially informative compounds that fall outside of canonical pathways. Another difficulty is due to the limited representation of MSI annotations in canonical metabolic pathways. Many metabolites, and particularly lipids detected by mass spectrometry, are absent from pathway databases or represented only at a generic level. Although resources such as HMDB [24] and LipidMaps [25] provide increasingly detailed molecular annotations, their correspondence with pathway resources remains incomplete. As a result, many spatially informative MSI features cannot be incorporated into conventional pathway level analyses. This limitation is well recognized in the literature on bulk metabolomics and lipidomics, where ontology- and structure-based strategies have been proposed to extend molecular coverage [26–28]. Its extent in spatial metabolomics and lipidomics has, however, not been systematically quantified. We refer to these assignments left unmapped to pathways as the “ghost metabolome”.

Here we introduce SpacePath, a computational framework for functional analysis of MSI-based spatial metabolomics and lipidomics. SpacePath addresses these two complementary sources of information loss (annotation ambiguity and limited canonical pathway coverage) within a single workflow. SpacePath propagates ion-to-compound ambiguity explicitly by distributing each ion’s signal across its compatible assignments. It then scores canonical pathways across tissue locations and groups assignments that were not mapped to pathways into *de novo* molecular families using chemical similarity. These families provide chemically interpretable groupings of molecules that were not mapped to pathways, extending the molecular organization of MSI data beyond canonical pathways.

We first quantify canonical pathway coverage across a diverse collection of public and novel MSI datasets and show that only a small fraction of annotated assignments can be linked to established metabolic pathways. We then evaluate SpacePath across public datasets with independent biological interpretation and assess its robustness through systematic computational perturbations. Finally, we apply the framework to newly acquired glioblastoma xenografts and use independent MS/MS measurements to confirm selected tumor-associated features from a *de novo* molecular family and refine their initial molecular assignments. Together, these analyses establish SpacePath as a framework for moving MSI interpretation beyond isolated ions toward spatially resolved canonical pathways and candidate chemical families, while retaining the uncertainty inherent to MS1-based molecular annotation.

## 2 Results

### 2.1 SpacePath enables functional interpretation of MSI-based metabolomics and lipidomics data

SpacePath takes as input an annotated MSI dataset comprising ion intensities across tissue locations, two-dimensional spot coordinates, and candidate molecular assignments for each detected ion (Fig. 1a). The framework adresses two major limitations that currently constrain functional interpretation of MSI data: the ambiguity of ion-to-molecule annotation and the poor representation of detected metabolites and lipids in canonical pathway databases.

**Fig. 1.**
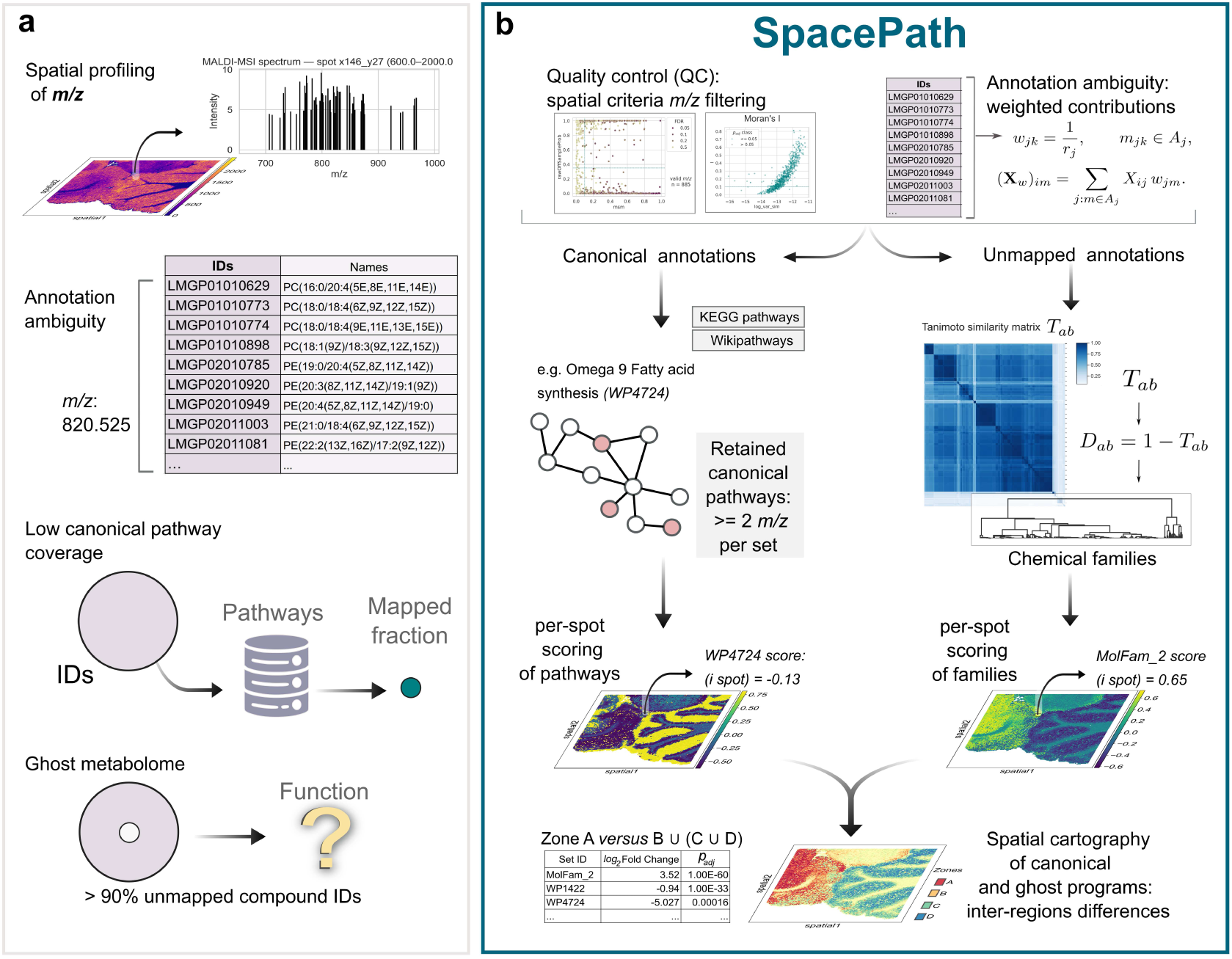
Conceptual overview of the analytical problem addressed by SpacePath and of the SpacePath framework. **a,** MSI-based spatial metabolomics and lipidomics generate spatially resolved ion-intensity profiles that are associated with one or more candidate molecular assignments. Two major sources of information loss limit downstream functional interpretation: ambiguity of molecular annotation and incomplete mapping of annotated compounds to canonical metabolic pathways. **b,** SpacePath processes annotated MSI profiles through quality and spatial filtering and, when required, spatial segmentation. Ambiguous ion signals are propagated across compatible molecular assignments. Assignments represented in reference pathway databases are used for spot-wise canonical pathway scoring, whereas pathway unmapped assignments are grouped according to chemical similarity into *de novo* molecular families and scored using the same framework. Pathway and family scores are subsequently combined for spatially resolved comparison between tissue regions.

SpacePath processes an annotated MSI dataset within a unified workflow (Fig. 1b). Ions are first filtered according to annotation quality and spatial structure, and tissue regions can be defined by unsupervised clustering or supplied externally. Candidate assignments incompatible with the biological context of the sample can be removed at this stage using curated exclusion lists (pharmaceuticals, plant-derived compounds, and bacterial or archaeal metabolites). All three lists were used in the analyses reported in this manuscript. SpacePath then distributes each ambiguous ion’s signal across its compatible molecular assignments, retaining annotation uncertainty in downstream analyses. Assignments mapped to reference pathway databases are integrated into spatially resolved pathway scores using kernel principal component analysis (k-PCA) [29], producing a pathway level representation for each tissue location.

A second core operation organises the annotated ions that fall outside canonical pathways. This unmapped to pathways “ghost metabolome” is grouped into *de novo* molecular families by chemical structural similarity, using ECFP4 fingerprints and Tanimoto distance (see Methods). These families are then scored across the tissue using the same k-PCA framework as the canonical pathways, giving canonical pathways and *de novo* families a common spatial representation. Region level comparisons can subsequently identify pathways and molecular families that distinguish spatial compartments.

The *de novo* families provide chemically coherent, hypothesis-generating groupings for spatially structured features that would otherwise remain outside higher level interpretation. They are not substitutes for experimentally validated metabolic pathways. This distinction is particularly important in MSI, where structural ambiguity is intrinsic to MS1-based annotation.

### 2.2 Canonical pathways capture only a small fraction of MSI annotations

A premise of SpacePath is that conventional pathway mapping leaves a large fraction of MSI molecular data without pathway level functional representation. Therefore, before evaluating the SpacePath framework, we quantified canonical pathway coverage across a broad collection of MSI spatial metabolomics and lipidomics datasets. The analysis included 27 public datasets retained after quality assessment, spanning different tissues, organisms and experimental conditions, and six newly acquired datasets corresponding to four mouse glioblastoma xenografts and two healthy-brain technical controls (Methods; Extended Data Fig. 2).

For each dataset, compound assignments provided in HMDB or LipidMaps nomenclature were mapped to KEGG [30] and WikiPathways [31, 32] (Fig. 2a–c). The mapping consistently lost most of the molecular information (Fig. 2d–g). Individual MSI datasets contained approximately 1,500–7,500 unique compound assignments, of which typically only 25–200 were associated with canonical functional sets. The loss was observed across both public datasets as well as those acquired in this study, irrespective of whether the starting annotations came from HMDB or LipidMaps or whether KEGG or WikiPathways was used as the pathway resource.

**Fig. 2.**
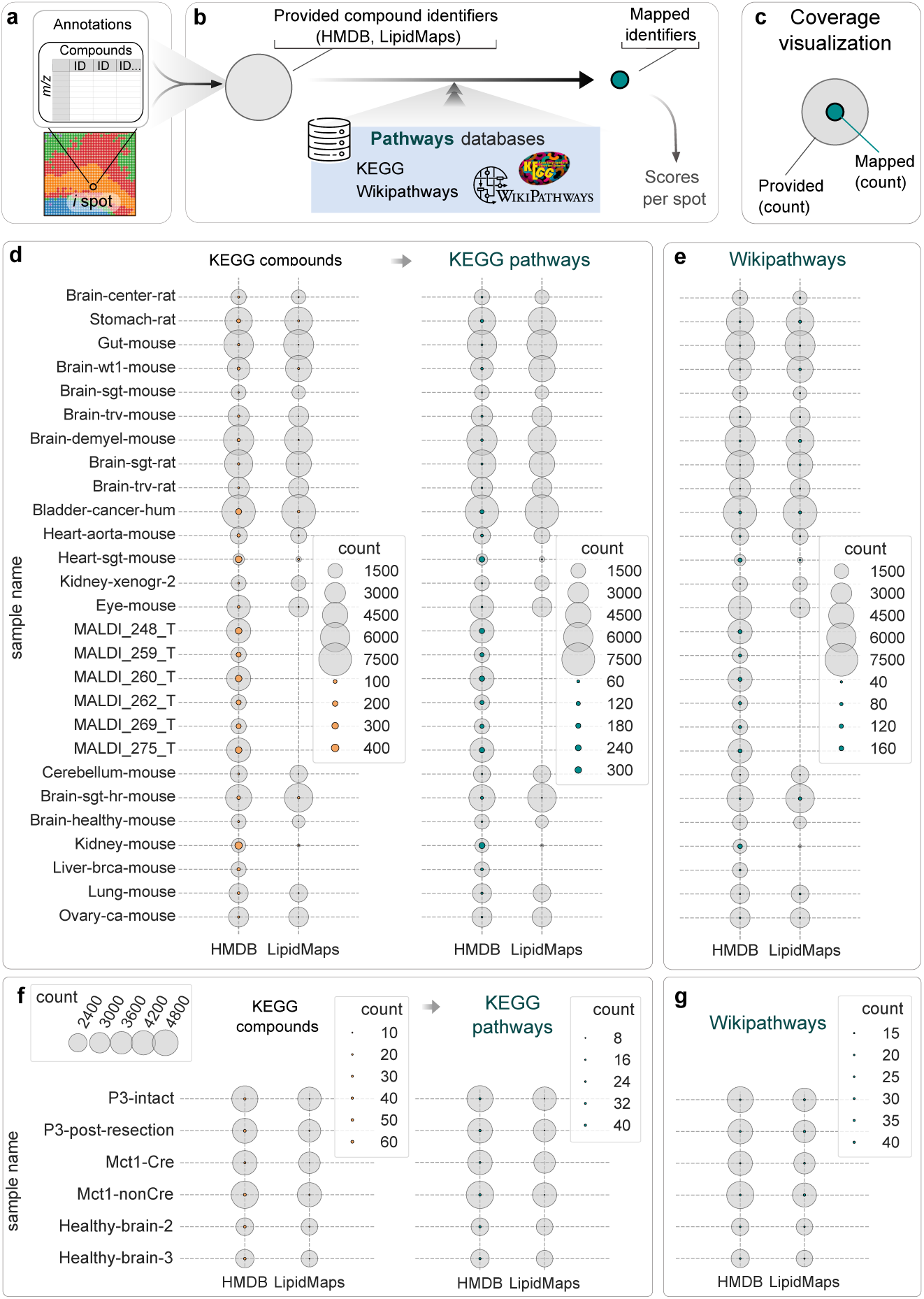
Canonical pathway coverage across MSI-based spatial metabolomics and lipidomics datasets. **a,** Compound annotations provided for each MSI dataset. **b,** Mapping of HMDB or LipidMaps identifiers to canonical functional sets using KEGG and WikiPathways. **c,** Coverage representation: outer circles indicate the number of provided compound identifiers and inner circles the subset successfully mapped to the indicated resource. **d,e,** Mapping coverage across the retained public datasets using KEGG (d) and WikiPathways (e). **f,g,** Corresponding analysis of the in-house datasets, including four glioblastoma xenografts, using KEGG (f) and WikiPathways (g). Circle diameter is proportional to the number of unique compound identifiers. IDs, compound identifiers.

Across all analyzed datasets, the mean dataset level canonical pathway coverage was 1.002% (1.03% using KEGG, 0.98% using WikiPathways). Approximately 99% of molecular assignments therefore remained outside the canonical pathway representation used here. The strongest bottleneck occurred during translation of detailed molecular identifiers to KEGG compound identifiers, reflecting the mismatch between the granularity of MSI annotations and the granularity represented in pathway databases (Fig. 2d,f). Limited pathway coverage has been recognised in bulk metabolomics and lipidomics [26, 27, 33]; our systematic survey shows that the limitation is particularly pronounced in MSI-based spatial profiling.

We next quantified the overall fraction of molecular assignments retained within the combined SpacePath representation. Across all analyzed datasets pooled together, 24.05% of the unique provided assignments were associated with either a canonical pathway or a *de novo* molecular family. This is a different quantity from the 1.002% above: 1.002% is the mean canonical-pathway coverage per dataset, whereas 24.05% is the pooled fraction of assignments covered by either canonical pathways or *de novo* families.

Importantly, most of these additional assignments are organised into chemically defined candidate families. We evaluated these *de novo* families by comparing their spatial distributions with lipid patterns reported in four published MSI studies.

### 2.3 SpacePath recovers established spatial lipid programs and extends their functional interpretation

We applied SpacePath to the 27 public datasets retained after quality assessment. For detailed benchmarking, we selected four spatial lipidomics datasets whose source publications provided explicit histological or molecular interpretations for direct comparison: healthy mouse brain [34], a murine model of demyelination [35], healthy mouse cerebellum [36], and human bladder cancer [37]. These datasets cover different tissues, biological conditions and MSI acquisition strategies. We asked whether SpacePath could recover the spatial molecular organization reported in the source studies and extend observations based on individual ions toward canonical pathways and *de novo* molecular families (Fig. 3).

**Fig. 3.**
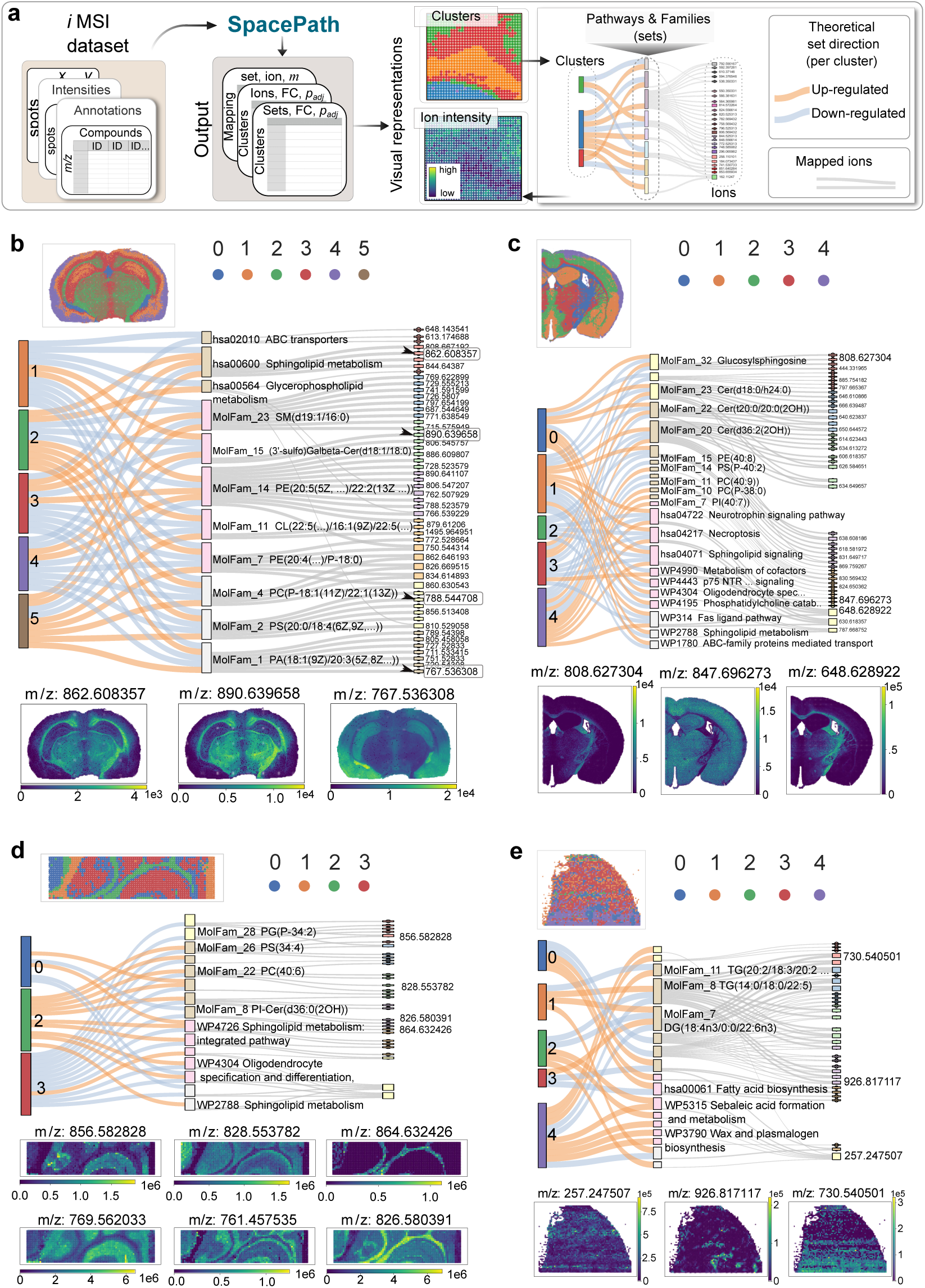
Validation of SpacePath across public spatial lipidomics datasets. **a,** Overview of the analysis and visualization of SpacePath outputs. Annotated MSI datasets are spatially segmented and analyzed to identify region-associated ions, canonical pathways and *de novo* molecular families. Alluvial diagrams connect spatial clusters to significant molecular sets and representative ions, while reconstructed ion-intensity maps provide direct spatial support for selected associations. **b,** Healthy mouse brain [34]. SpacePath recovers lipid programs associated with gray- and white-matter organization, including sphingolipid metabolism and molecular families of phosphatidylserines and galactosylceramides. **c,** Murine demyelination model [35]. SpacePath identifies spatially differentiated lipid programs associated with preserved white-matter tracts and the demyelinated callosal region. **d,** Healthy mouse cerebellum [36]. SpacePath segmentation recapitulates major cerebellar layers and associates them with distinct canonical pathways and molecular families. **e,** Human muscle-invasive bladder cancer [37]. SpacePath resolves tumor, intratumoral smooth-muscle and surrounding regions and identifies compartment-specific lipid pathways and molecu_7_lar families. Only representative significant associations are displayed for readability.

In the healthy mouse brain dataset of Clarke et al. [34], the original study established pronounced spatial differences in lipid composition between gray-and white-matter structures. SpacePath segmented the section into six molecularly distinct regions and recovered this organization at both canonical pathway and molecular family levels (Fig. 3b). ‘Sphingolipid metabolism’ was enriched in myelin-rich regions, consistent with the distribution of *m/z* 862.608357, annotated as 3-O-sulfogalactosylceramide. A phosphatidylserine family (Mol-Fam 2) was enriched in white matter. SpacePath also recovered *m/z* 890.639658, assigned to Gal*β*-Cer(d42:1), through a galactosylceramide family (MolFam 15) despite its absence from the canonical pathway representation. A phosphatidic acid family (MolFam 1) was enriched in gray-matter regions. These spatial patterns agreed with those reported in the original study despite differences in annotation methods, the original analysis used proprietary software with manual curation, whereas we used METASPACE.

The demyelination dataset of Konieczna-Wolska et al. [35] provided a benchmark of a different kind because the original analysis focused on cholesterol redistribution following myelin loss induced by cuprizone and did not examine global spatial lipid organization. SpacePath partitioned the brain section into five molecular regions (Fig. 3c). A top ranked *de novo* family of glucosylsphingosine-related species (MolFam 32), represented by *m/z* 808.627304, was enriched in the internal capsule and optic tract and showed reduced signal in the demyelinated callosal region. Canonical pathways associated with sphingolipid and neuronal signaling also displayed clear regional organization, including p75 NTR receptor-mediated signaling in white-matter structures. Additional region-specific associations are reported in the Supplementary Information.

In the healthy cerebellum dataset of Bond et al. [36], the original study related individual lipid distributions to the characteristic histological layers of the cerebellum. SpacePath resolved spatial clusters corresponding to the molecular layer, granular layer and white matter, in addition to a non-tissue background cluster (Fig. 3d). Among the top ranked programs, a phosphatidylserine family (MolFam 26) and ‘Sphingolipid metabolism’ were enriched in the molecular layer, and a phosphatidylcholine family (MolFam 22) was enriched in the granular layer. White matter was associated with ‘Sphingolipid metabolism’ and a PI-ceramide family (MolFam 8), with representative ions such as *m/z* 864.632426 reproducing the expected localization to myelin-rich regions. Individual phosphatidylcholine distributions identified using Metaspace were spatially concordant with those reported by Bond et al. using MassPix. These lipid associations reflected the layered organization of the cerebellum.

In the bladder cancer dataset of Tu et al. [37], the original study reported strong spatial variation in lipid composition between tumor, smooth muscle and surrounding tissue. SpacePath identified five compartments consistent with this histological organization, including the tumor core, intratumoral smooth muscle and peri-tumoral/non-tumor regions (Fig. 3e). Among the top ranked programs, ‘Fatty acid biosynthesis’ and a triacylglycerol family (MolFam 8) were enriched in the smooth muscle compartment, consistent with the original observation that higher-mass lipid species preferentially localized outside the tumor core. The non-tumoral regions also carried fatty acid associated programs, including the spatial distribution of *m/z* 257.247507, assigned to palmitic acid. A diglyceride family (MolFam 7), represented by *m/z* 730.540501, was enriched in the tumor core. Additional associations, including wax- and plasmalogen-related signals in non-tumor regions, are reported in the Supplementary Information.

Across the four datasets, SpacePath recovered major spatial lipid patterns reported in the original studies, including when different annotation methods were used. Molecular *de novo* families additionally captured regional associations involving lipids absent from the canonical pathway representation. Computational perturbation and sensitivity analyses are reported in Supplementary Section S2 and Supplementary Figs. 1–2.

### 2.4 SpacePath identifies spatial lipid programs in glioblastoma xenografts

We applied SpacePath to four original in-house spatial lipidomics datasets of glioblastoma xenografts acquired via MALDI-MSI (Methods). These datasets encompassed two experimental conditions: (i) human P3 cell tumors in immunodeficient mice, including intact and post-resection scenarios, and (ii) murine glioblastoma cell tumors in immunocompetent mice, with either normal or inhibited lactate transporter (Mct1) expression in macrophages. We first examined the intact xenograft generated from human P3 GB cells in an immunodeficient mouse (P3-intact). Leiden clustering identified six spatial compartments corresponding to the pia mater, outer and inner cortex, striatum, corpus callosum and tumor (Fig. 4a). Canonical pathways and *de novo* molecular families were scored across these compartments, giving a spatial molecular representation of both normal brain structures and the tumor-associated lipid landscape.

**Fig. 4.**
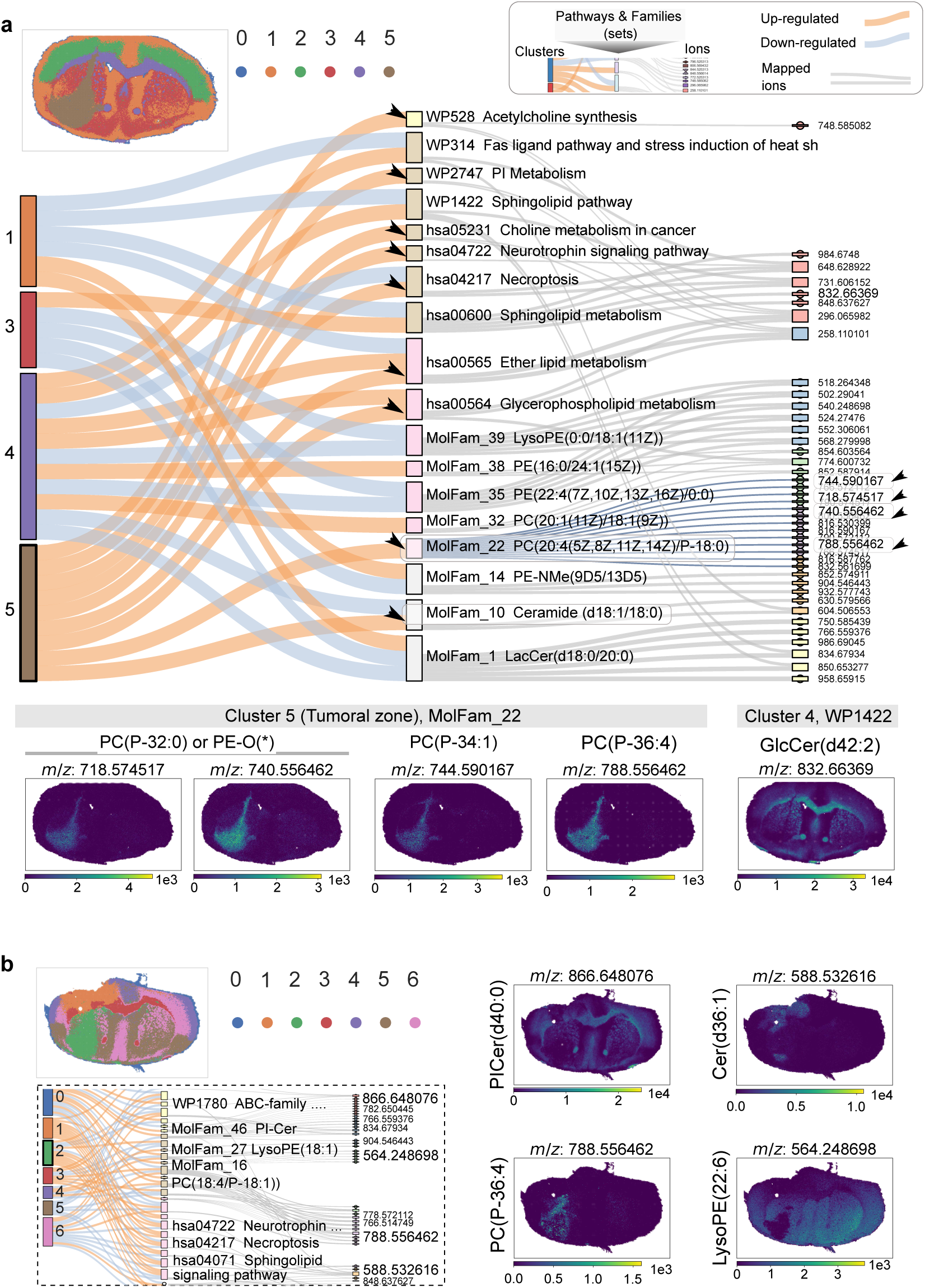
SpacePath analysis of glioblastoma xenografts. **a,** Intact P3 glioblastoma xenograft (P3-intact). Upper, Leiden clustering resolves six spatial regions corresponding to pia mater, outer cortex, inner cortex, striatum, corpus callosum and tumor. Center, representative associations between spatial clusters, canonical pathways, *de novo* molecular families and contributing ions. Bottom, spatial distributions of representative ions, including four tumor-associated ions grouped in MolFam 22 and a glucosylceramide associated with the corpus callosum. **b,** Post-resection P3 xenograft (P3-post-resection). Upper-left, Leiden clustering resolves seven regions including surgical scar, recurrent tumor and corpus callosum. Lower-left, representative canonical pathways and molecular families associated with these regions. Right, spatial distributions of selected ions associated with the corpus callosum, surgical scar and tumor. Only representative significant associations are displayed for readability.

In the corpus callosum, SpacePath identified lipid programs consistent with its myelin-rich composition, including ‘Sphingolipid metabolism’, ‘Ether lipid metabolism’ and ‘Glycerophospholipid metabolism’ (Fig. 4a). Representative signals included *m/z* 832.663690, assigned to GlcCer(d42:2), which localized to the callosal region. The tumor compartment was associated with glycerophospholipid, ether-lipid and choline-related pathways, but these collectively represented only a small number of detected ions. Molecular family analysis also identified tumor-associated lipid programs among assignments that were not mapped to pathways.

A top ranked *de novo* tumor-associated molecular family (MolFam 22) grouped candidate molecular assignments from seven ions and was enriched in the GB region (Fig. 4a). Four of these ions (*m/z* 718.574517, 740.556462, 744.590167 and 788.556462) showed sufficiently strong tumor-associated signals to permit independent investigation using SpiderMass coupled to tandem mass spectrometry (MS/MS).

SpiderMass-MS/MS independently detected all four selected features in the tumor region (Extended Data Fig. 5 and Extended Data Table 1). The initial MS1 annotations included phosphatidylcholine (PC) candidates for all four ions and ether-linked phosphatidylethanolamine alternatives for two. Fragmentation spectra supported phosphatidylethanolamine (PE) 34:1 for *m/z* 718.574517 and PE 36:4 for *m/z* 740.556462. The feature at *m/z* 744.590167 was compatible with either PE 36:2 or a HexCer species, whereas *m/z* 788.556462 was consistent with PC 36:1. These results demonstrate how a SpacePath family can guide the selection of tumor-associated ions for independent chemical investigation.

We also examined a post-resection P3 xenograft (P3-post-resection), in which seven spatial compartments included the surgical scar, tumor and surrounding brain structures (Fig. 4b). The corpus callosum retained a ceramide-rich signature, including a PI-Cer-associated family, while distinct lipid families were associated with the surgical scar and tumor. A phosphatidylcholine-associated family (MolFam 16, represented by *m/z* 788.556462) was enriched in the tumor region, whereas a LysoPE-associated family displayed a contrasting spatial distribution. Because this dataset represents a single post-resection specimen, these findings describe within-tissue spatial associations and do not establish systematic metabolic differences between primary and recurrent tumors.

The application of SpacePath to each of the four GB xenograft slices is summarized in Supplementary Figs. 9-11.

## 3 Discussion

SpacePath addresses a central limitation of MSI-based spatial metabolomics and lipidomics: the gap between rich spatial molecular measurements and their higher level functional interpretation. Current MSI analyses resolve ion distributions across tissues, but two sources of information loss remain present: uncertainty in assigning molecular identities to MS1 features, and the limited representation of annotated metabolites and lipids in canonical pathway databases. SpacePath treats both limitations explicitly. It distributes each ambiguous ion’s signal across its compatible molecular assignments, and it organizes assignments that are not mapped to pathways into *de novo* molecular families defined by chemical similarity. Canonical pathways and *de novo* families are scored spatially and compared across tissue regions, moving MSI analysis from individual ions toward spatially organized molecular programs.

Across the datasets examined here, canonical pathway mappings represented about 1% of candidate molecular assignments per dataset on average. The public benchmarks showed that lipids outside this representation could nevertheless reproduce anatomical distributions reported in the original studies.

Chemical structure and ontology have previously been used to address incomplete pathway mapping in metabolomics. Metabolomics2Network uses ontology relationships to connect detailed lipid annotations with the more generic entities represented in metabolic networks, whereas ChemRICH defines chemically related sets for enrichment analysis [26, 27]. SpacePath applies chemical grouping to spatial MSI while retaining alternative candidate identities for individual ions. Its *de novo* families allow assignments absent from reference pathways to be examined collectively across tissue regions. Their interpretation rests on the candidate structures and spatial associations they represent.

A molecular family that is associated with a given region provides a set of ions for targeted chemical investigation. Examining the contributing ion maps helps to identify features that support the regional association and are suitable for downstream MS/MS experiments. In the glioblastoma example, four ions selected from one tumor-associated family were independently detected in the tumor, while fragmentation spectra supported PE identities for two ions whose initial annotations included both PC and ether-linked PE candidates. These findings support the experimental relevance of the selected signals and show why the chemical interpretation of a family should be revisited when molecular assignments change. Because family construction depends on candidate structures, revised identities may also alter its membership if the analysis is repeated with updated annotations. The experiment therefore demonstrates how SpacePath can prioritize ions for subsequent molecular identification.

The approach remains dependent on the quality and completeness of molecular annotations. Equal allocation of an ion’s signal across compatible assignments provides a practical treatment of ambiguity, but does not estimate the relative probabilities of those identities or quantitatively separate co-isomeric species. Low mass MSI data present additional difficulties because matrix signals, spectral overlap and acquisition artifacts can compromise annotation reliability [38, 39]. The MS/MS investigation of selected members of one family provides limited evidence about families elsewhere in the datasets.

Further development should connect annotation evidence more directly to family construction and scoring. Candidate-specific confidence estimates could inform signal allocation, while MS/MS or ion mobility evidence could constrain the structures used to construct families. Repeating the analysis after such refinements would allow the stability of family membership and regional associations to be assessed. Comparisons across biological conditions will additionally require replicated sampling and statistical models that account for variation between specimens. These developments would help determine which spatial lipid programs recur across samples and which depend on unresolved molecular assignments.

SpacePath provides a framework for interrogating MSI data at a level between individual molecular features and established metabolic pathways. It retains annotation uncertainty and organizes spatially informative assignments that remain unmapped to pathways into chemically defined molecular families. Combining these families with canonical pathway scoring provides a basis for generating testable molecular hypotheses from spatial metabolomics and lipidomics data.

## 4 Methods

### 4.1 Study design and datasets

#### 4.1.1 Glioblastoma xenograft models and tissue processing

Four glioblastoma (GB) xenograft brains were analyzed in two experimental cohorts. In the first cohort, human P3 GB cells were implanted into the striatum of immunodeficient RAG*γ*2C^−/−^ mice (*n* = 2). One animal was monitored without further intervention (P3-intact), whereas the second underwent surgical tumor resection 40 days after implantation and was subsequently monitored for tumor regrowth (P3-post-resection). In the second cohort, murine GL261 GB cells were implanted into one *Mct1* ^fl/fl;Cre/−^ and one *Mct1* ^fl/fl;nonCre^ mouse (hereafter Mct1-Cre and Mct1-nonCre) in a macrophage-specific *Mct1* conditional model. The experimental designs are summarized in Extended Data Fig. 1.

Animals were euthanized at the predefined humane endpoint. Brains were collected, freshfrozen, sectioned coronally at 10 µm, thaw-mounted onto standard glass microscope slides and stored at −80 *^◦^*C until MSI acquisition. Animal procedures and the provenance of the P3 cells are described in the Ethics declarations.

Three additional partial brain sections from healthy mice were acquired under the same experimental protocol and served as technical healthy-brain controls. Seven novel spatial lipidomics datasets were acquired in total: four GB xenografts and three healthy-brain sections (Supplementary Table 3). Six met the inclusion criteria described in Public MSI datasets, inclusion criteria and benchmark selection (Healthy-brain-1 was excluded).

#### 4.1.2 Public MSI datasets, inclusion criteria and benchmark selection

A corpus of 29 public spatial metabolomics or lipidomics MSI datasets was assembled from METASPACE and the associated literature. The collection covered different tissues, organisms, biological conditions and acquisition protocols. Only MALDI- or IR-MALDESI-based acquisitions generated on high-resolution FT-ICR or Orbitrap instruments were included. TOF acquisitions were excluded because their mass accuracy did not meet the annotation requirements used in this manuscript. All retained acquisitions were MS1 datasets. Most were spatial lipidomics; six spatial metabolomics acquisitions came from from Ravi et al. [23]. Many datasets had been generated primarily for technical or instrumentation studies, but a smaller subset had explicit biological interpretation in the corresponding publication.

Dataset suitability for cross-dataset analyses was assessed using four quantities calculated from KEGG and WikiPathways mappings: (i) the assignment-to-ion ratio, defined as the number of unique compound assignments divided by the number of detected *m/z* features; (ii) the number of ions represented in retained pathways, with at least two distinct ions required per pathway; (iii) the number of retained pathways; and (iv) the mean number of mapped ions per pathway. Datasets were retained when the assignment-to-ion ratio was ≤ 2.2, at least 10 pathways met the mapping criterion for at least one mapping strategy, and the mean number of mapped ions per pathway was ≥ 1 (Extended Data Fig. 2). Twenty-seven of the 29 public datasets and six of the seven newly generated datasets met these criteria. The excluded public datasets were Kidney-xenogr-1 and Brain-lmrange-mouse.

Four of the retained public datasets provided sufficiently explicit histological or molecular interpretation in the source publication to support direct biological benchmarking: healthy mouse brain [34], a cuprizone-associated demyelination model [35], healthy mouse cerebellum [36], and human muscle-invasive bladder cancer [37]. These four were used for the detailed concordance analyses in Fig. 3.

Different analyses in the manuscript use different subsets of the full dataset corpus. Dataset provenance and quality outcomes for every dataset are listed in Supplementary Table 1; the composition of each analysis set is listed in Supplementary Table 2.

### 4.2 MSI acquisition and annotation

#### 4.2.1 MALDI-MSI acquisition of mouse spatial lipidomics data

Frozen sections were processed at the Bordeaux Proteome facility. Sections (12 µm-thick) were cut from the brain using an NX70 Star cryostat (Thermo Fisher Scientific) and thaw-mounted onto standard glass microscope slides.

For Matrix-Assisted Laser Desorption Ionization (MALDI)-MSI analysis, a uniform matrix layer was applied using a custom-built pneumatic sprayer. Specifically, 150 µm of a 2,5-dihydrobenzoic acid (DHB) matrix solution (20 mg mL^−1^ in methanol:water, 1:1 v/v) was sprayed onto tissue sections under the following optimized conditions: a matrix solution flow rate of 0.05 mL min^−1^ and a nitrogen gas flow of 0.9 bar.

Acquisitions were performed using an atmospheric pressure imaging ion source named AP-SMALDI 5 AF (TransMIT GmbH) connected to an orbital trapping mass spectrometer (QExactive Orbitrap, Thermo Fisher Scientific). The latter was operated in positive ion mode at a mass resolution of 140 000 @ *m/z* 200 over a mass range of *m/z* 200-1000. The ion source is equipped with a diode laser (Flare NX343, *λ* = 343 nm), operating at a repetition rate of 2 kHz, with a maximum pulse energy of 200 µJ at source attenuated to 20 nJ at the sample. Matrix clusters were used for online lock-mass calibration during acquisition to compensate for mass shifting. Images were acquired at 25 µm lateral resolution.

Preprocessed data were converted with Thermo Fisher Scientific software to imzML format [40] and exported as paired `.imzML` and `.ibd` files.

#### 4.2.2 Annotation of novel mouse datasets and reannotation of public datasets

All acquired MSI datasets were annotated with METASPACE [12]. Annotation parameters were: a resolving power of 140,000, a mass tolerance of 3 ppm, M+H and M+Na adducts, and the HMDB, CoreMetabolome, LipidMaps and SwissLipids databases. The METASPACE analysis type was set to `Original MSM`. Spectral alignment was performed with MSIWarp [39]. METASPACE annotation confidence and off-sample metrics were retained for downstream SpacePath filtering.

For the four GB xenografts, SpacePath filtering used FDR ≤ 0.20, off-sample probability ≤ 0.35, MSM ≥ 0.10, Moran’s *I* ≥ 0.10 and BH-adjusted Moran permutation *P* ≤ 0.05. The resulting feature sets contained 687 ions for P3-intact, 698 for P3-post-resection, 674 for Mct1-Cre and 735 for Mct1-nonCre.

Public datasets were reannotated or supplemented with METASPACE annotations when the original records did not contain the compound identifier types required by SpacePath. Where reannotation was necessary, mass tolerance, adduct definitions and related annotation parameters followed the corresponding publication or METASPACE registry record.

### 4.3 Data input, normalization and identifier handling

#### 4.3.1 Input representation and TIC normalization

For each MSI dataset, SpacePath requires (i) a spot-by-ion intensity matrix W ∈ ℝ*^n×m^*, where *n* is the number of spatial locations and *m*the number of detected *m/z* features; (ii) two-dimensional coordinates *S* = {*s*_1_*, …, s_n_*}; and (iii) an annotation table 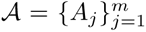, where *A_j_* is the set of candidate compound identifiers associated with ion *j*. Internally, SpacePath also uses compound-to-pathway mappings for canonical sets and chemical descriptors for those assignments that are unmapped to pathways. LipidMaps, HMDB, CoreMetabolome and KEGG identifiers can be supplied simultaneously.

Ion intensities were normalized by total ion count (TIC). For spot *i*,

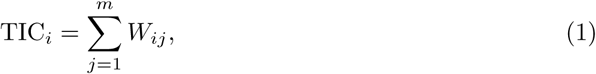

and the normalized intensity of ion *j* was

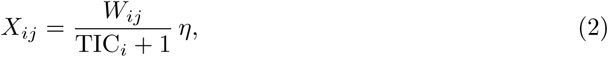

where *η* = median*_i_*(TIC*_i_*). The additive constant prevents division by zero. Unless explicitly stated otherwise, downstream spatial and differential analyses used the normalized matrix X ∈ ℝ*^n×m^*.

#### 4.3.2 Identifier harmonization and optional assignment filtering

Compound identifiers were harmonized before pathway mapping or molecular-family construction. Alias resolution allows assignments supplied in HMDB or LipidMaps nomenclature to be linked to the corresponding KEGG [30] and WikiPathways [31, 32] entities where valid cross-references are available. For database releases and retrieval dates see Data availability. Alias resolution precedence, deduplication and multiplet handling are detailed in the Supplementary Methods. Multiple candidate records associated with the same *m/z* feature are retained as alternative assignments rather than collapsed to a single molecular identity.

SpacePath additionally provides optional exclusion lists for identifiers covering three compound classes: pharmaceuticals, plant-derived compounds, and bacterial or archaeal metabolites. These lists can be applied before pathway mapping and molecular family construction to remove candidate assignments considered incompatible with the biological context. In this study all three exclusion lists were enabled, discarding candidate identifiers of non-animal origin across all datasets.

### 4.4 Ion quality and spatial filtering

SpacePath applies two successive filters. First, the MSM, FDR and off-sample scores reported by METASPACE are used to filter ion–formula annotation records. MSM summarises agreement between the experimental and theoretical isotopic pattern and spatial colocalization; FDR estimates the expected proportion of false ion–formula assignments; the off-sample score estimates the probability that a signal originates outside the tissue [41]. The thresholds are user-configurable and stored in the SpacePath parameter file for each dataset. When several formulas are assigned to one ion, the ion is retained if at least one formula passes the thresholds. These quality metrics attach to ion–formula records, whereas the subsequent spatial test operates on ion-intensity vectors; retained annotation records are therefore consolidated to their ion set before Moran testing.

Recommended defaults for SpacePath are MSM ≥ 0.3, FDR ≤ 0.20 and off-sample probability ≤ 0.20. For the analyses reported here, thresholds were tuned per dataset to retain a workable number of features (per dataset values are provided in Supplementary Table 2).

Second, spatial structure is assessed with Moran’s *I* on the TIC-normalised ion-intensity vector. For an ion profile x = (*x*_1_*, …, x_n_*),

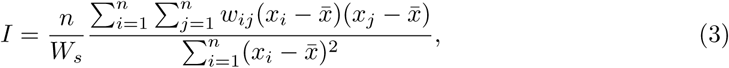

where *w_ij_* denotes the spatial weight between spots *i* and *j* and *W_s_* = Σ*_i,j_ w_ij_*. Spatial graphs were constructed from Euclidean spot coordinates using a *k*-nearest-neighbour graph (*k* = 6 in the analyses reported here); graph construction and autocorrelation were computed via `squidpy.gr.spatial neighbors` and `squidpy.gr.spatial autocorr`. Moran’s *I* was computed for each ion retained by the first filter. Significance was assessed with 999 per-mutations and corrected using Benjamini–Hochberg method. An ion was retained if its Moran’s *I* was positive and both the *I* and adjusted-*P* thresholds specified for that dataset were met.

### 4.5 Spatial region definition

SpacePath can use externally supplied region labels or derive regions directly from the matrix of normalized ion intensities. The current implementation supports *k*-means clustering [42] and Leiden community detection [43]. When regions were inferred by SpacePath, clustering used ion intensities only. Histological labels were not included as clustering features. Cluster labels were subsequently compared with tissue anatomy for interpretation.

For the public benchmark datasets, SpacePath used *k*-means with *k* = 6 (Clarke healthy brain), *k* = 5 (demyelination) and *k* = 4 (cerebellum), and Leiden clustering at resolution 0.27 (muscle-invasive bladder cancer). For the P3-intact and P3-post-resection xenografts, Leiden clustering at resolution 0.35 was applied. Cluster assignments were then propagated to the downstream ion- and molecular-set comparisons. Per-dataset clustering parameters (algorithm and parameter values) are listed in Supplementary Table 2.

### 4.6 Ion level spatial differential analysis

For each spatial region, ion intensities were compared with all remaining spots using the Wilcoxon rank-sum test. Tests were performed independently for each region versus rest contrast and *P* values were adjusted by the Benjamini–Hochberg procedure within each contrast. Ions with adjusted *P* ≤ 0.05 and | log_2_ FC| ≥ 0.8 were considered to be associated with the region.

For reporting and visualization, SpacePath retains up to 100 ranked ions per region, that include the 70 ions with the highest relative intensity and the 30 with the lowest relative intensity. This truncation limits exported/visualized ion lists and does not define the universe of ions that is statistically evaluated.

### 4.7 Ambiguity-aware compound representation

MS1 annotation commonly assigns several candidate molecular identities to the same *m/z* feature. SpacePath instead retains every compatible assignment. Let *A_j_* = {*m_j_*_1_*, …, m_jrj_*} denote the *r_j_* compatible compound assignments for ion *j*. In the current implementation, all compatible assignments receive equal allocation weight,

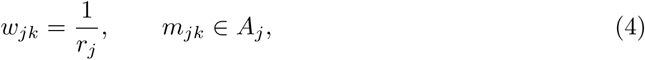

such that 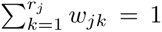. These coefficients define a signal-allocation rule that prevents the full intensity of one ion from being counted independently for every compatible molecular identity.

The full weighted matrix X*_w_* is constructed by summing weighted contributions from all ions compatible with compound *m*.

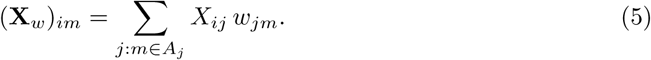

For example, an ion of intensity *X_ij_* with two candidate compounds contributes *X_ij_/*2 to each. A compound supported by several ions receives the sum of their weighted signals, the total contribution of an individual ion is conserved across its candidate assignments. This ambiguity-aware compound representation is used as input to canonical-pathway and molecular-family scoring.

### 4.8 Canonical pathway mapping

Harmonized compound assignments were mapped to canonical metabolic sets from KEGG [30] and WikiPathways [31, 32]. A canonical pathway was retained for scoring only when it contained at least two compound assignments originating from at least two distinct *m/z* features in the analyzed dataset. This requirement limits sets supported by a single ambiguous ion. Compound assignments can participate in more than one canonical pathway when the source databases contain overlapping sets.

### 4.9 Construction of *de novo* molecular families

Candidate assignments from the significant differential ions (adjusted *P* ≤ 0.05) within the top *n* per region lists are grouped according to chemical similarity to rescue a spatially relevant fraction of assignments that could not be linked to a retained canonical pathway. In this study, *n* was set to 25 for stringency.

For each eligible assignment, an ECFP4 fingerprint was generated with RDKit [44]. Candidate fingerprints were pooled within each dataset and pairwise structural similarity between compounds *a* and *b* was quantified with the Tanimoto coefficient *T_ab_*. Similarities were converted to dissimilarities: *D_ab_* = 1 − *T_ab_*.

The distance matrix was partitioned by agglomerative clustering to define molecular families per dataset. The default linkage was Ward’s method, computed on the condensed distances matrix. Families were formed by cutting the dendrogram at a distance threshold.

Family membership was defined at the compound level using correspondence tables and propagated back to ion signals. Alternative assignments of a single ambiguous ion can therefore contribute to different families. A molecular family was retained only when supported by at least two compound assignments originating from at least two distinct *m/z* features. Per-dataset family counts are reported in Supplementary Table 2.

### 4.10 Per spot scoring of pathways and molecular families

Let *G* denote the canonical pathways and *de novo* molecular families retained for a dataset. Each set *g* ∈ *G* is represented by a subset *M_g_* of columns of the compound matrix X*_w_*. The corresponding submatrix for set *g* is X*_g_* = X*_w_*[:*, M_g_*].

SpacePath transforms X*_g_* to a single score per spatial spot using kernel principal component analysis (kPCA) with a radial basis function (RBF) kernel [29]. For spots *i* and *j*,

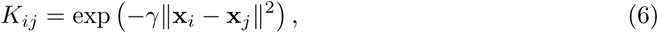

where x*_i_* and x*_j_* are the weighted compound profiles at the two spots. The bandwidth was 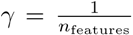, where *n*_features_ is the number of features in X*_g_*. The kernel matrix is centered before decomposition, and the first kernel principal component is retained as the spatial score vector s*_g_* ∈ ℝ*^n^*. Scoring uses the kPCA procedure exposed by the `sspa` package [45]. A set is scored only when it contains at least two compound assignments originating from at least two distinct *m/z* features.

The score vectors of all retained canonical pathways and molecular families are concatenated into a common score matrix 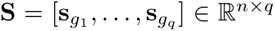, where *q* is the total number of scored sets.

### 4.11 Region level differential analysis of molecular sets

For each spatial region, canonical pathway and molecular family scores were compared with scores in all remaining spots. Because set scores can be negative or positive, a shift transformation S ← S+ | min(S)| was applied before the comparison to avoid division by zero. Effect sizes were computed as log-transformed ratios of geometric means, and statistical significance was assessed with the Wilcoxon rank-sum test for each region-versus-rest contrast. The *p*-values were adjusted for multiple testing using the Benjamini-Hochberg procedure, applied jointly across canonical pathways and molecular families within each region-versus-rest comparison (i.e., separately per region contrast).

In this study, molecular sets meeting BH-adjusted *P* ≤ 0.05, |log_2_ FC| ≥ 0.8 and Moran’s *I* ≥ 0.2 were considered spatially differentiated. The 30 with the highest |log_2_ FC| were retained and are referred to as top-ranked molecular sets or families throughout the Results. These thresholds can be modified by the user.

### 4.12 Canonical and combined molecular set coverage

Canonical pathway coverage was calculated from the unique compound assignments supplied for each retained dataset. For dataset *d* and pathway resource *b*, let *U_d_* denote the set of unique provided compound assignments and 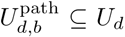 the assignments mapped to at least one retained canonical pathway in resource *b*. Dataset level coverage was was defined as

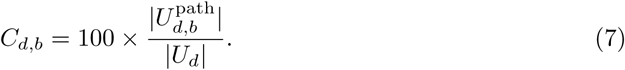

Coverage was calculated independently for KEGG and WikiPathways and averaged across datasets. The reported 1.03% and 0.98% values correspond to the mean dataset level coverage for KEGG and WikiPathways, respectively; 1.002% is the mean of these two database-specific averages.

To quantify representation after molecular family construction, assignments from all ana-lyzed datasets were pooled and the fraction associated with either a retained canonical pathway or a SpacePath molecular family was calculated relative to the pooled set of provided assignments. This analysis yielded the 24.05% overall representation reported in the Results.

### 4.13 Systematic computational perturbation and runtime analyses

The stability of SpacePath was evaluated by comparing the baseline workflow with two perturbations. In the no-filter configuration, the MSM, FDR, off-sample and Moran-based ion filters were disabled. In the random-ID configuration, one candidate compound identifier was selected at random for each ambiguously annotated ion instead of the equal allocation rule in Eq. 5. The random ID configuration is a destructive hard assignment stress test.

The configurations were compared using four summaries calculated from the resulting molecular set scores: (i) region versus rest effect sizes; (ii) Moran’s *I* of the set score maps; (iii) an ambiguity propagation score for *de novo* families; and (iv) the number of molecular sets passing adjusted *P* ≤ 0.05. These quantities were evaluated separately by dataset and set type. For a molecular family *F* and a given *m/z* feature represented in that family, let *S*_in_ denote the set of contracted (sum composition-level notation) molecular species assigned to *F* at that *m/z*, and let *S*_out_ denote the species assigned to other families at the same *m/z*. The ambiguity contribution of the feature was

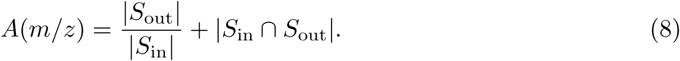

The ambiguity propagation score of family *F* was the mean contribution across the *m/z* features represented in that family,

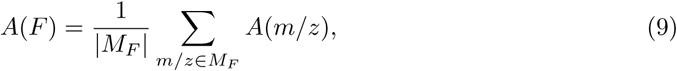

where *M_F_* is the set of contributing *m/z* features. Lower values indicate less propagation of candidate level ambiguity across molecular families.

A separate computational comparison measured kPCA and GSVA runtime (in seconds) for canonical pathway scoring across 30 selected MSI datasets: 27 of the public cohort (after excluding Brain-lmrange-mouse and Brain-sgt-hr-mouse datasets) and 3 in-house datasets (specifically comprising Healthy-brain-1, Healthy-brain-2, and Healthy-brain-3). Identical input matrices and KEGG pathway definitions were used for both methods. The number of spatial spots varied across datasets. Detailed results are reported in the Supplementary Information.

### 4.14 Software implementation and reproducibility

SpacePath was implemented in Python 3.10 and is distributed as a command line workflow. A run is defined by a dataset metadata/parameter file, an input directory and an output directory. The command line layer orchestrates analytical functions implemented in the nested library. The same core functions and data structures can be imported programmatically for notebook or pipeline use.

SpacePath uses `sspa` [45] for kPCA-based single-sample molecular-set scoring; `AnnData, scanpy` and `squidpy` for matrix, statistical and spatial operations; `pyimzML` for imzML input; RDKit and `fastcluster` for chemical representation and molecular-family construction; and `pydantic` for data validation. All computations were performed on a server equipped with two AMD EPYC 7413 24 core processors (48 physical cores, 96 logical CPUs), running Rocky Linux 9.4; jobs were allocated up to 120 GB of memory, with a peak usage of 15.99 GB. The workflow was also tested on Ubuntu 22.04 and macOS Sequoia 15.5. Source code, example datasets and dataset specific parameter files are provided under Code availability and Data availability.

### 4.15 Tandem MS characterization by SpiderMass

Four tumor-associated *m/z* features selected from MolFam 22 in the P3-intact xenograft were investigated independently by SpiderMass coupled to tandem mass spectrometry (MS/MS). Sampling positions were chosen within the tumor region using the MALDI-MSI images with the localization of each ion of interest.

The SpiderMass platform was configured as described by [46–48] with remote laser micro-sampling coupled through a transfer line to a REIMS interface and mass spectrometry. The laser system comprised a pulsed Nd:YAG source (6-ns pulse duration, 20 Hz) pumping a tunable optical parametric oscillator (Opolette, OPOTEK). Sampling used a 1-m biocompatible optical fiber (450-µm inner diameter) with a handpiece fitted with a 4-cm focusing lens. Laser energy was 4 mJ per pulse for a 10-s irradiation period, corresponding to an estimated fluence of 3 J cm^−2^. Ablated material was transferred through 2 m of Tygon ND 100-65 tubing (2.4-mm inner diameter, 4-mm outer diameter) to the REIMS interface. Before each acquisition, isopropanol was infused at 200 mL min^−1^ with leucine enkephalin (200 mg mL^−1^) as lock mass. Spectra were acquired in positive ion sensitivity mode with a 1-s scan time over *m/z* 50-1000. Tandem MS was performed on a Waters Xevo G3 mass spectrometer.

Selected precursor ions were isolated and fragmented by collision-induced dissociation in the transfer cell, with collision energies of 30-40 eV adjusted per precursor. Lipid identities were assigned by manual interpretation of diagnostic fragmentation patterns and comparison with LipidMaps [25], Alex123 [49], MetFrag [50] and published spectra. MS/MS provided independent chemical characterization for the four ions selected by SpacePath. The remaining members of the molecular family were not tested.

## Extended data

**Extended Data Fig. 1.**
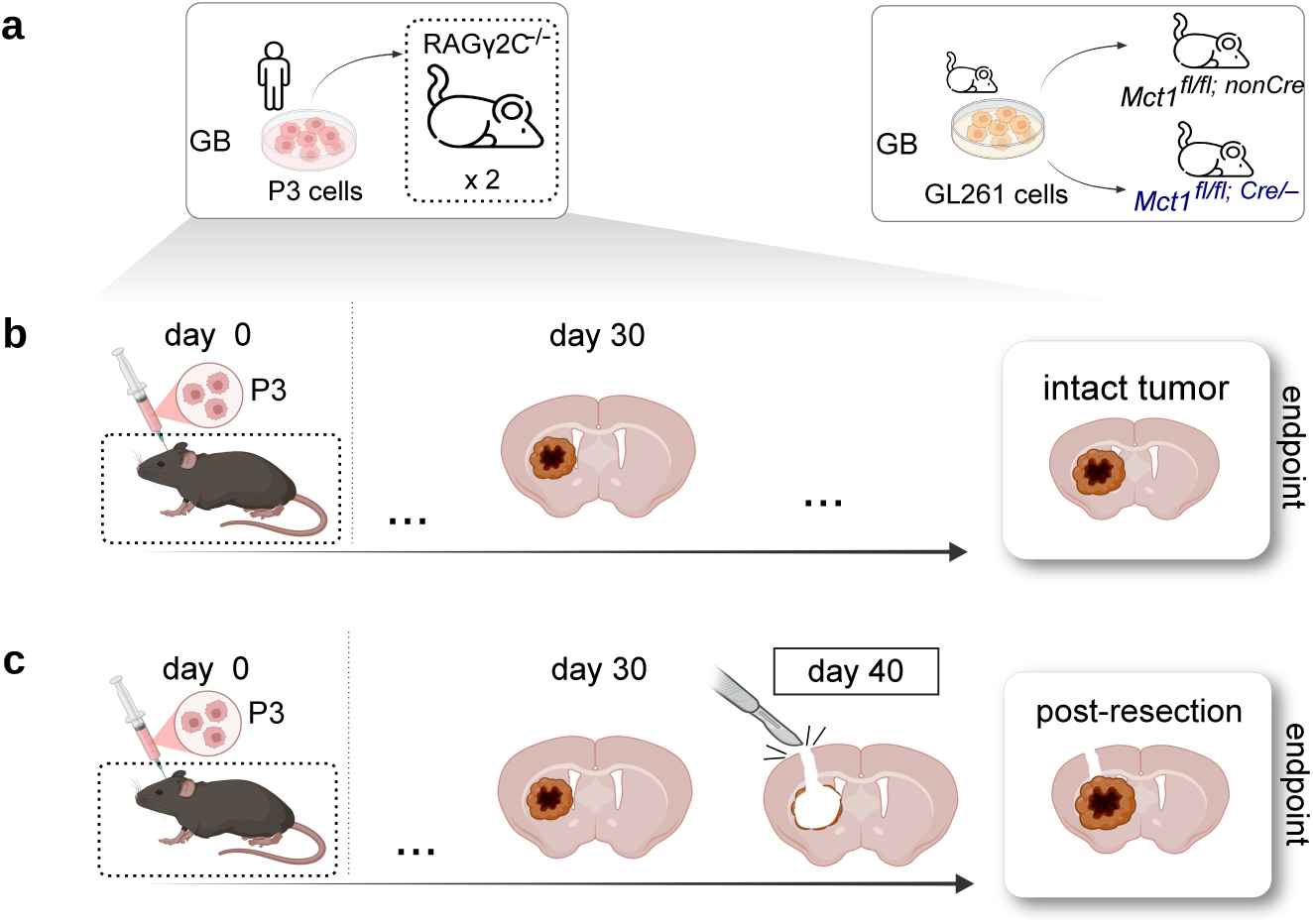
Glioblastoma Xenografts experiment schema. Experiments of the GB xenograft model for spatial lipidomics data acquisitions. **a,** specific cell lines and mouse strains configured two different cohorts: a-left, immunodeficient mice, injected with human P3 glioblastoma cells. a-right, Black6 mice (*Mct1* ^fl/fl;nonCre^ and *Mct1* ^fl/fl;Cre/−^), injected with murine GB cells. **b** and **c** depict the design followed for the mice of the cohort defined in a-left. See Methods for a detailed description of the experiment. The figures were elaborated with Biorender^®^ and Inkscape software.

**Extended Data Fig. 2.**
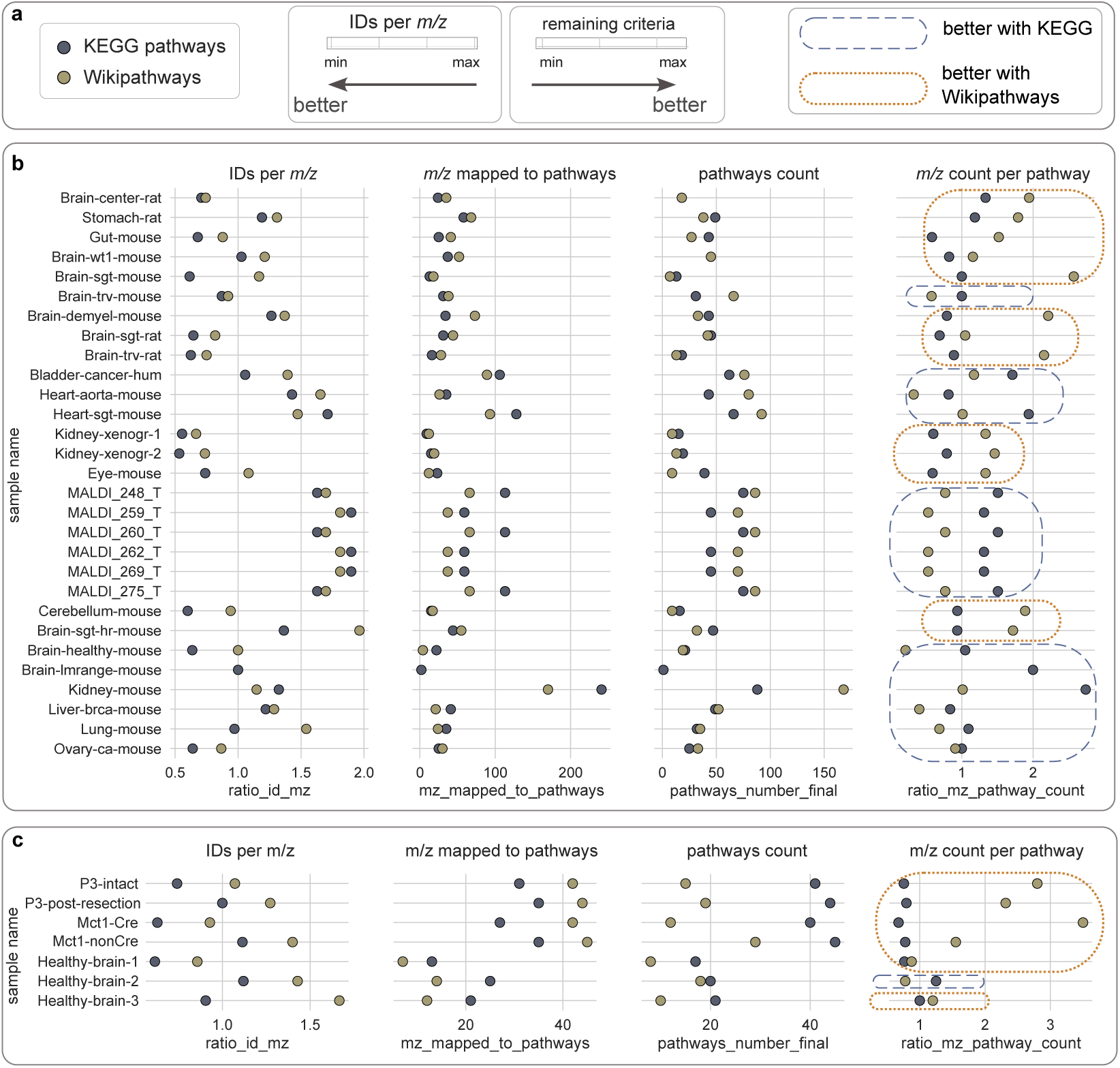
Mapping metrics across MSI lipidomics and metabolomics datasets. **a** legend of the presented evaluation. a-left, the types of pathways that were included. a-center, ‘IDs per *m/z*’ is the quotient of the total count of unique compound identifiers (IDs) by the total count of *m/z*s: values *>* 1 indicate annotation ambiguity (multiple assignments to one single ion) which negatively affects mapping efficiency. Values *<* 1 indicate overlap of assignments between pairs of ions, which is tolerated in this study. The remaining criteria are ‘*m/z* mapped to pathways’, ‘pathways count’ and the ratio of ‘*m/z* count per pathway’ (Methods): for these three criteria the bigger the value, the better the mapping efficiency. a-right, line patterns to encircle the type of pathway for which the efficiency was better, per pathway. Also thresholds for considering the dataset for further analyses were: IDs per *m/z* ≤ 2.2, at least 10 enriched pathways, and ion-per-pathway ratio ≥ 1 (Methods). **b** public datasets evaluated for mapping metrics. Thresholds were unmet by the samples Kidney-xenogr-1 and Brain-lmrange-mouse so they were excluded from the subsequent analyses. c, novel datasets including four glioblastoma xenografts and three additional partial tissue sections. Thresholds were unmet by one sample (Healthy-brain-1) so it was excluded from the subsequent analyses.

**Extended Data Fig. 3.**
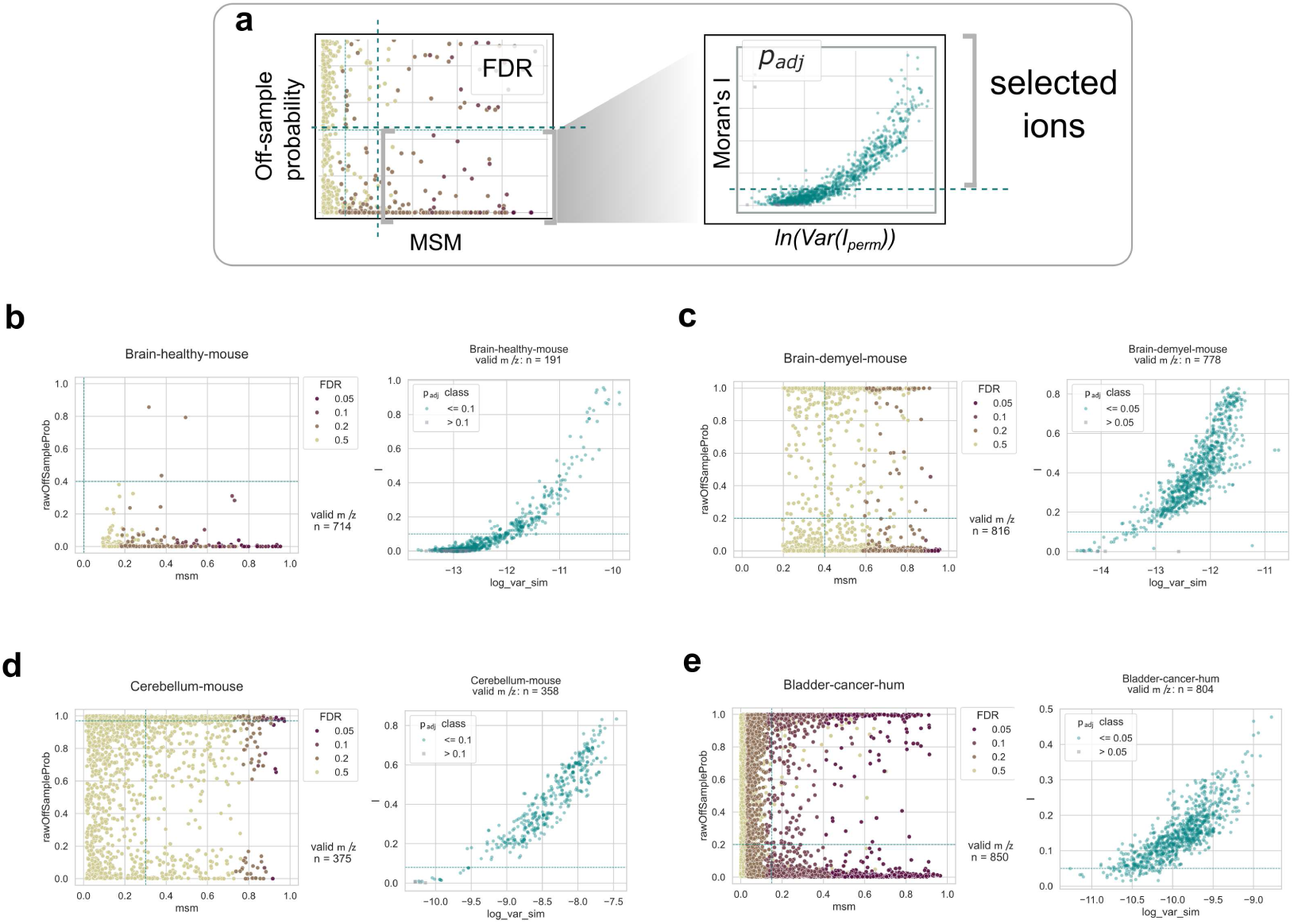
Features across public datasets: number of ions retained after application of quality thresholds. The four MSI datasets from public sources used for SpacePath benchmarking are shown. The scatter-plots represent the ions and their associated quality metrics. Each ion (*m/z*) is represented by a dot in each scatter plot area. **a,** legend explaining the logic of the SpacePath ions filtering: a-left, MSM values (x-axis), off-sample probabilities (y-axis) and FDR (color scale); the thresholds (dashed lines) enable the first phase of filtering by SpacePath. a-right, the ions passing the first phase undergo Moran’s I computation: the x-axis shows the log-transformed permutations variance (*ln*(*V ar*(*I_perm_*))) and the y-axis the I values. Thresholds for I values and *p*_adj_ values enable SpacePath to perform a second (final) phase of filtering. The thresholds were fine-tuned to retain a reasonable number of good quality ions for subsequent analysis. **b,** healthy mouse brain dataset. **c,** demyelinated brain dataset from a murine model. **d,** mouse cerebellum dataset. **e,** human muscle invasive bladder cancer. Ion and *m/z* are exchangeable terms in this work. The visualizations were automatically produced by the SpacePath workflow.

**Extended Data Fig. 4.**
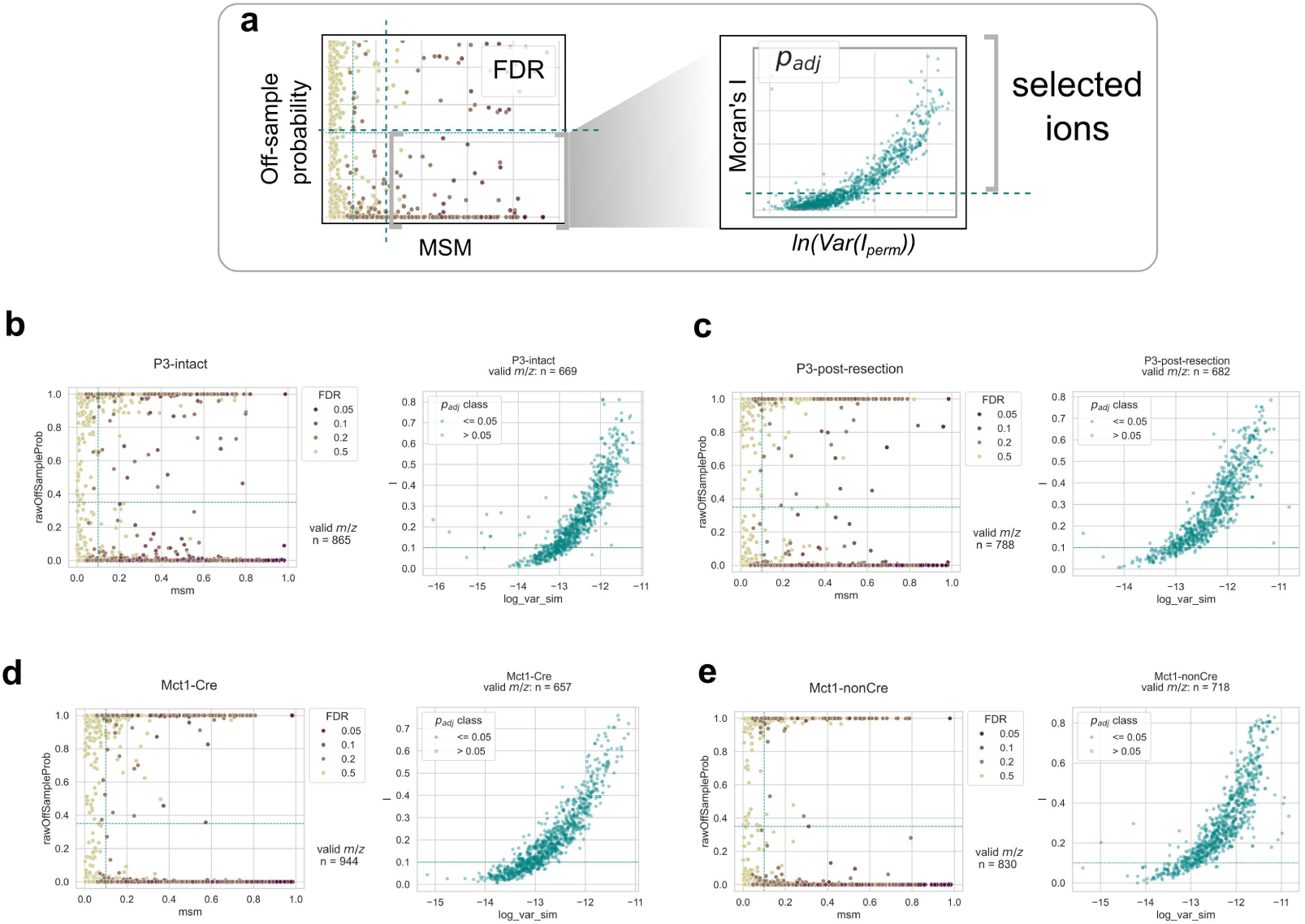
Features across in-house tissues of the glioblastoma xenograft models. The spatial lipidomics datasets of our in-house glioblastoma xenografts are shown. The scatter-plots represent the ions and their associated quality metrics. Each ion (*m/z*) is represented by a dot in each scatter plot area. **a,** legend explaining the logic of the SpacePath ions filtering: a-left, MSM values (x-axis), off-sample probabilities (y-axis) and FDR (color scale); the thresholds (dashed lines) enable the first phase of filtering by SpacePath. a-right, the ions passing the first phase undergo Moran’s I computation: the x-axis shows the log-transformed permutations variance (*ln*(*V ar*(*I_perm_*))) and the y-axis the I values. Thresholds for I values and *p*_adj_ values enable SpacePath to perform a second (final) phase of filtering. The thresholds were fine-tuned to retain a reasonable number of good quality ions for subsequent analysis. **b,** non-resected P3 tumor in immunocompromised host. **c,** post-resection P3 tumor in immunocompromised host. **d, e,** tissues of murine glioblastoma (GL261) in Black6 mice with a conditional knock-down of a lactate transporter in macrophages. Ion and *m/z* are exchangeable terms in this work. The visualizations were produced by the SpacePath workflow automatically.

**Extended Data Table 1.**
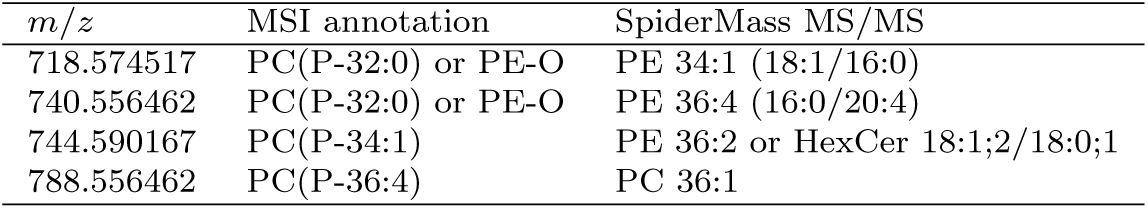
Independent MS/MS characterization of SpacePath-selected tumor ions. The MSI column reports candidate molecular assignments obtained from the initial MS1-based annotation, whereas the SpiderMass MS/MS column reports assignments supported by fragmentation spectra.

| *m*/*z* | MSI annotation | SpiderMass MS/MS |
|---|---|---|
| 718.574517 | PC(P-32:0) or PE-O | PE 34:1 (18:1/16:0) |
| 740.556462 | PC(P-32:0) or PE-O | PE 36:4 (16:0/20:4) |
| 744.590167 | PC(P-34:1) | PE 36:2 or HexCer 18:1;2/18:0;1 |
| 788.556462 | PC(P-36:4) | PC 36:1 |

**Extended Data Fig. 5.**
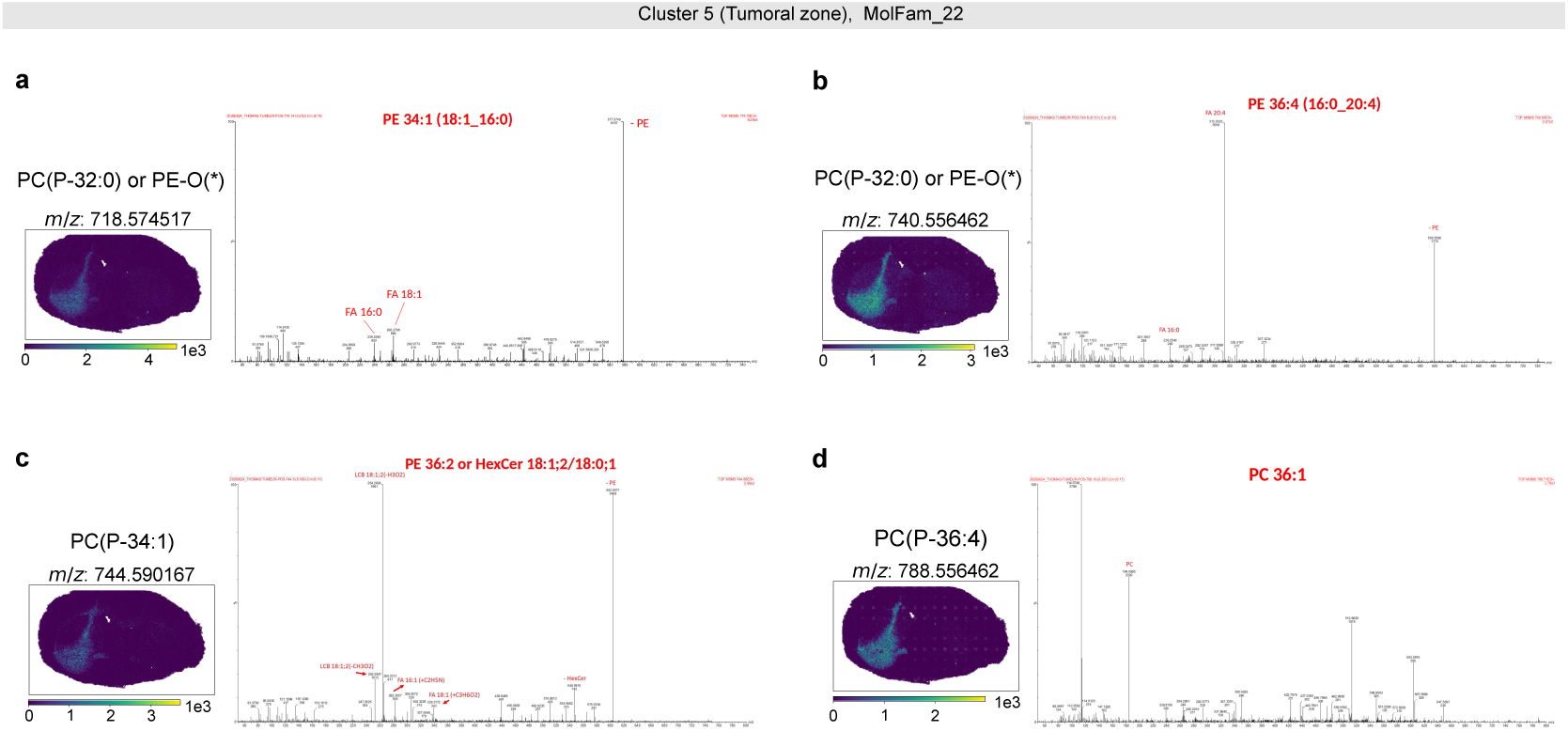
Independent MS/MS characterization of ions selected from the tumor-associated MolFam 22. Each panel (**a–d**) shows the spatial distribution of one SpacePath-selected *m/z* feature in the intact xenograft (left) together with its SpiderMass tandem mass spectrum acquired from the tumor region (right). All four features were independently detected by SpiderMass-MS/MS. Fragmentation spectra refined the initial MS1 assignments: *m/z* 718.574517 to PE 34:1 (**a**), *m/z* 740.556462 to PE 36:4 (**b**), *m/z* 744.590167 to PE 36:2 or HexCer (**c**), and *m/z* 788.556462 to PC 36:1 (**d**).

## 5 Data availability

Example datasets of small size, that allow for a fast test of the SpacePath pipeline, are available at https://zenodo.org/records/19113236.

The complete collection of preprocessed datasets used in this research is available at the Zenodo record https://zenodo.org/records/23085023 This collection includes novel glioblastoma xenografts and accompanying–healthy brain–calibration datasets. The METAS-PACE registries of the corresponding raw datasets are specified in the .tsv files within https://github.com/cbib/SpacePath/datasets_parameters.

The Metaspace links to the original xenograft datasets are: https://metaspace2020.org/dataset/2025-05-26_11h43m25s (P3-intact), https://metaspace2020.org/dataset/2025-07-30_16h54m35s (P3-post-resection), https://metaspace2020.org/dataset/2025-05-29_14h37m38s (Mct1-Cre), https://metaspace2020.org/dataset/2025-05-28_16h28m09s (Mct1-NonCre). The in-house healthy brain sections are available at https://metaspace2020.org/dataset/2025-07-28_15h11m01s (Healthy-Brain-1), https://metaspace2020.org/dataset/2025-07-28_15h09m59s (Healthy-Brain-2) and https://metaspace2020.org/dataset/2025-07-28_15h08m37s (Healthy-Brain-3).

The following databases and versions were used in this study: HMDB: Version 5.0 (released 2021-11-17), downloaded on 2024-05-07. LipidMaps: Release version 2024-10-15, downloaded on 2024-10-18. KEGG: Release 110.0 (2021-04-01), downloaded on 2024-04-16. WikiPathways: Version 20250410 (released 2025-04-10), downloaded on 2025-04-15.

## 6 Code availability

SpacePath, including its command-line workflow and importable core analytical library, is avail-able at https://github.com/cbib/tree/main/SpacePath. The release used for this manuscript, its software environment, dataset-specific configurations and reproduction instructions are archived at DOI https://doi.org/10.5281/zenodo.23390206. Identifier-mapping tables are distributed with the software. Scripts used to construct these reference tables are archived at https://zenodo.org/records/23079160.

The code used for both the systematic computational perturbation, and runtime analyses, is available at https://github.com/cbib/comp-evaluations-spacepath.

## Supplementary Information

The Supplementary Material is provided in a separate .pdf file.

## Acknowledgements

Thanks to CBiB members for IT infrastructure support. To Dr Theodore Alexandrov and his team, for scientific support regarding the usage of the METASPACE platform.

## Author information

All authors equally contributed.

## Ethics declarations

P3 cells were derived from patient donor samples; the regional ethical committee approved the collection of biopsy tissue at Haukeland University Hospital, Bergen, Norway (REK 013.09). All mice were housed under specifically pathogen-free conditions at the Animalerie Mutualisee de Talence (University of Bordeaux, France) on a 12-h dark/light cycle with unlimited access to food and water. All experiments were performed with 8- to 11-week-old male mice and were randomly allocated to the treatment groups. Male RAG*γ*2C^−/−^ mice were born, housed, and treated in the same animal facility. All animal procedures have been done according to the institutional guidelines and approved by the local ethics committee (agreement number: A23802).

## Supplementary material

### S1 SpacePath workflow overview

The SpacePath workflow is described in detail in the Methods section of the main manuscript and summarized schematically in Supplementary Fig. 1. Briefly, SpacePath integrates spatial ion-intensity profiles with candidate molecular assignments, applies ion-quality and spatial filtering, and defines spatial regions for downstream comparison. An ambiguity-aware compound representation is then used to score canonical pathways and chemically defined *de novo* molecular families at each spatial location. These scores are combined into a common molecular-set matrix for region-level comparison.

**Supplementary Fig. 1.**
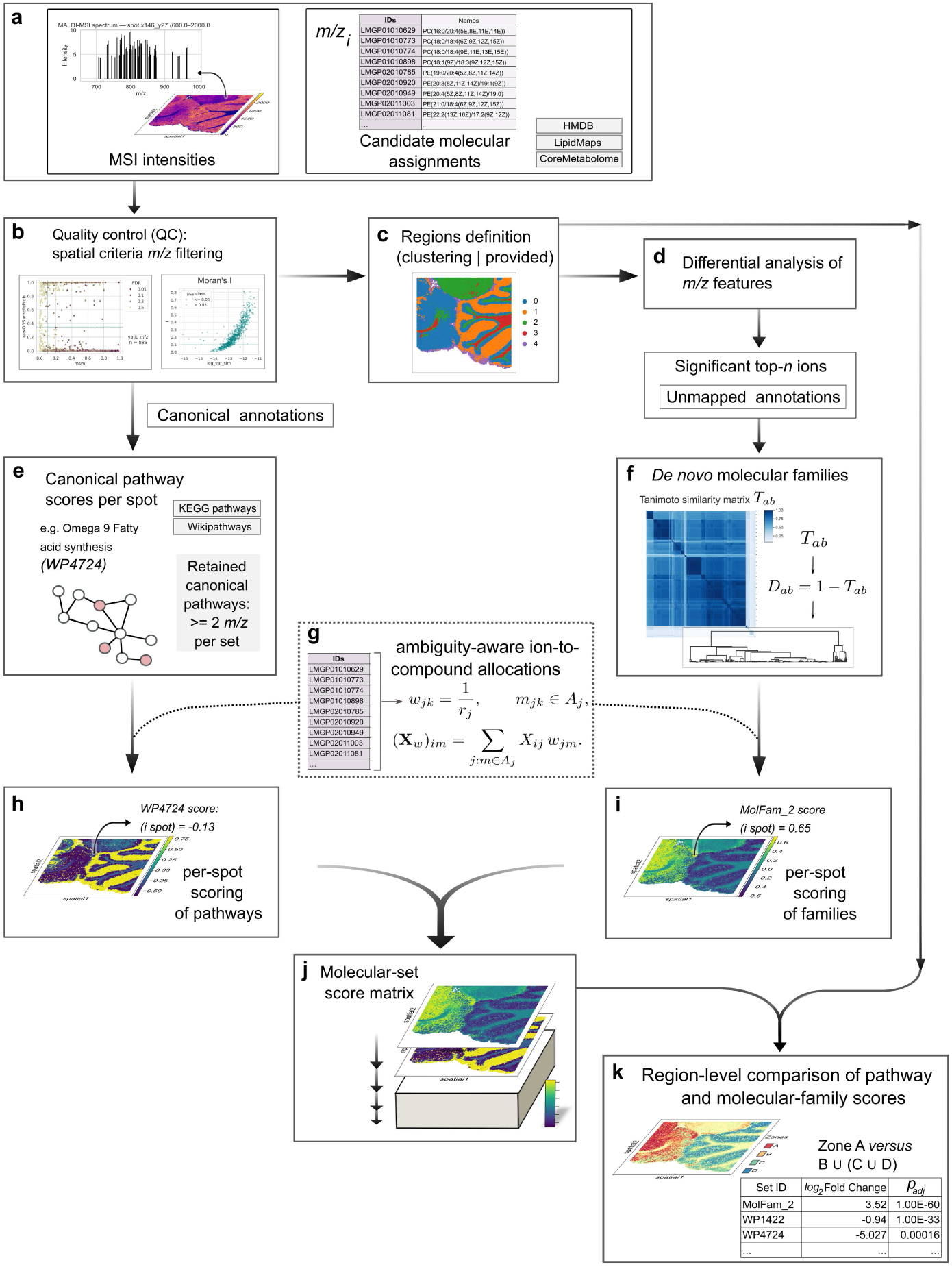
Overview of the SpacePath workflow. SpacePath integrates spatial ion-intensity profiles with candidate molecular assignments, applies ion-quality and spatial filtering, defines spatial regions, constructs an ambiguity-aware compound representation, and derives spatial scores for canonical pathways and *de novo* molecular families. These scores are combined into a common molecular-set matrix for region-level comparison.

**Supplementary Table 1.**
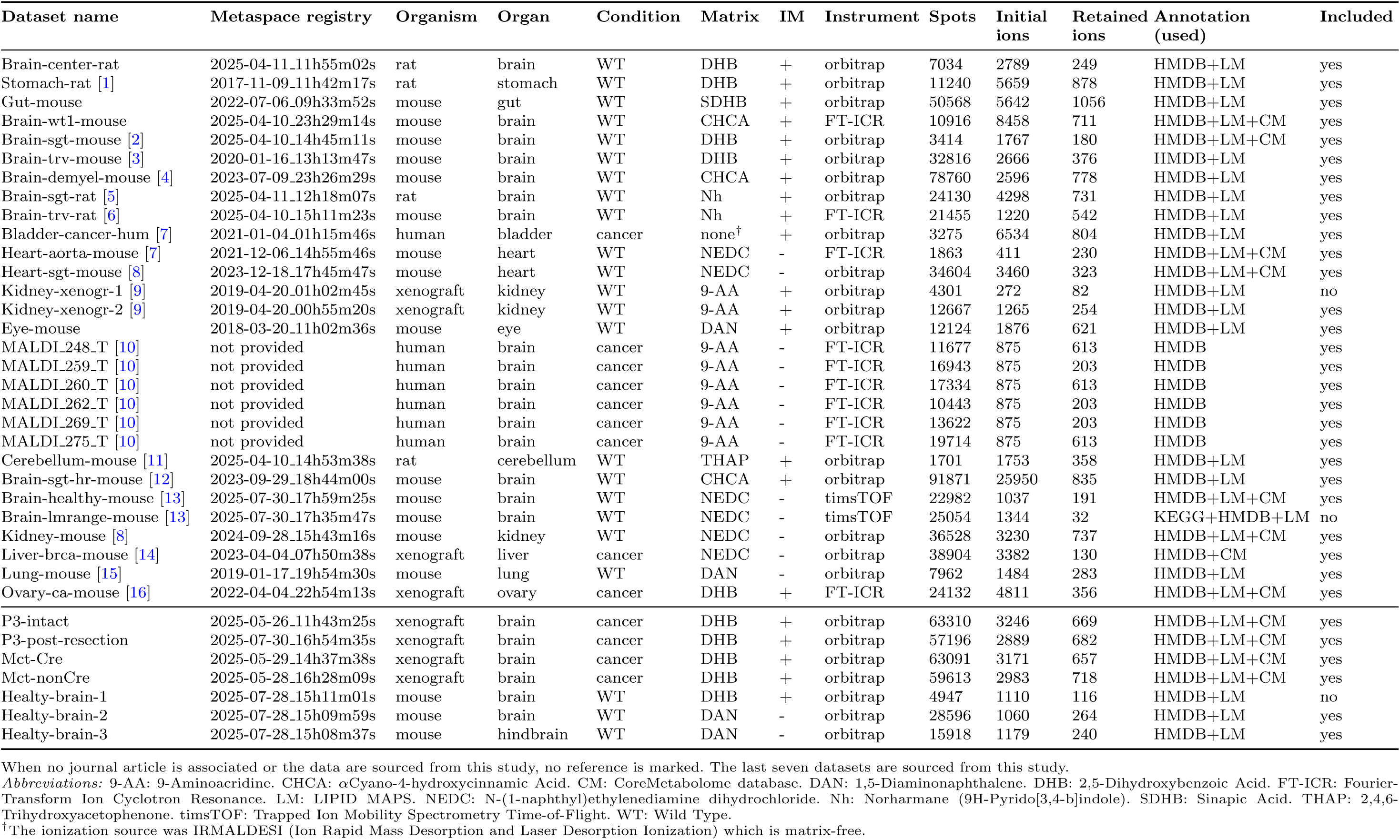
Dataset provenance and selection role.

Throughout the Supplementary Information, the term *molecular set* refers collectively to canonical pathways and *de novo* molecular families.

#### S1.1 Dataset provenance and analysis inclusion

The provenance and analytical role of all public and study-generated datasets are summarized in Supplementary Table 1. Public datasets were assembled from METASPACE, the associated literature, and other specified sources (e.g. MetaboLights), whereas the study-generated datasets comprised four glioblastoma xenografts and three healthy-brain sections. Dataset inclusion criteria and the distinction between the cross-dataset, benchmarking, glioblastoma, perturbation, and MS/MS analysis sets are described in the main Methods.

#### S1.2 Dataset-specific SpacePath parameters

Dataset-specific SpacePath settings are summarized in Supplementary Table 2. Parameters that were constant across analyses are reported in the main Methods and are not repeated here. The table therefore focuses on settings that varied between datasets, including ion-filtering thresholds, spatial region definition, clustering parameters and molecular-family construction settings used for each analysis.

**Supplementary Table 2.**
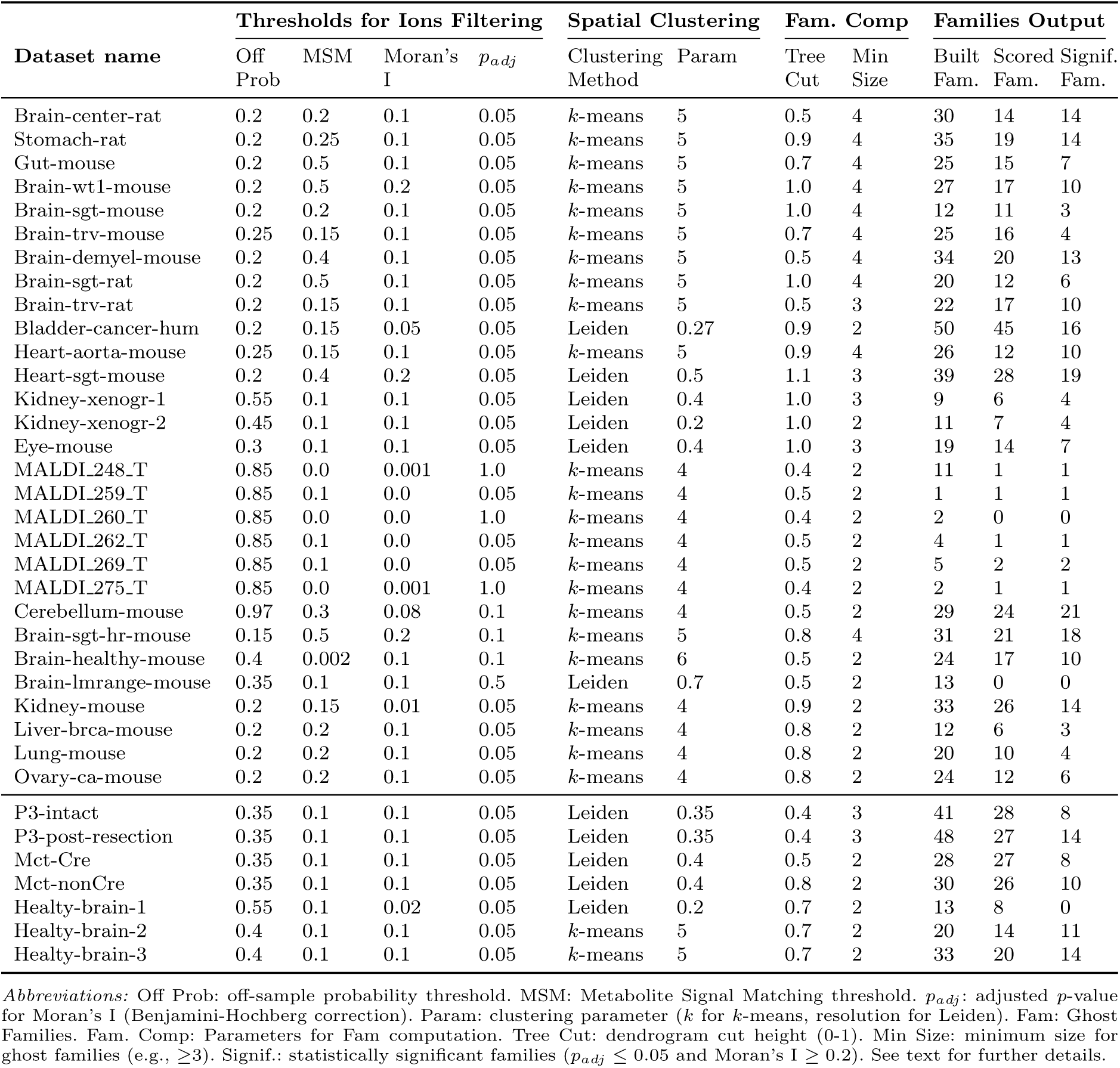
Dataset-specific SpacePath parameters. Only parameters that varied between datasets are shown; invariant settings are described in the main Methods.

As shown in the Supplementary Table 2, the ‘off-sample probability’ (Off Prob) and Metabolite Signal Matching (MSM) thresholds were dataset-specific to retain sufficient ions (lower Off Prob = higher stringency; MSM closer to 1 = higher stringency). The *p_adj_* thresholds for Moran’s I were relaxed for specific datasets (e.g., 0.5 for Brain-lmrange-mouse) to preserve features. Regarding the False Discovery Rate (FDR) reported by Metaspace (linked to the MSM), the threshold was set to 0.2 for Brain-sgt-hr-mouse, 0.5 for all the rest (as only varied for one dataset it was omitted in the table). Canonical pathways satisfied ≥2 compounds from ≥2 *m/z* as general rule, except for Heartsgt-mouse (≥3 from ≥3 *m/z*). The number of families in the Supplementary Table 2 was registered at three stages: i) before Min Size condition evaluation, i.e. after grouping of assignments by chemical composition (Built); ii) after satisfying Min Size and undergoing per-spot scoring (Scored); and iii) after differential analysis if satisfying *p_adj_*≤ 0.05 and Moran’s I ≥ 0.2 (Signif.).

The paths of the datasets configuration files are: SpacePath/datasets parameters/public-datasets-parameters.tsv (for the public datasets) SpacePath/datasets parameters/inhouse-datasets-parameters.tsv (for the in-house datasets)

### S2 Systematic computational evaluation of SpacePath

The perturbation analysis was designed to examine two complementary components of the SpacePath workflow: ion filtering, which limits the contribution of noisy or spatially unstructured features, and ambiguity-aware allocation, which avoids forcing ambiguous MS1 signals to a single candidate identity. These analyses (available at https://github.com/cbib/comp-evaluations-spacepath) are intended as sensitivity tests of the SpacePath representation rather than as biological validation.

#### S2.1 Ablation design

The baseline SpacePath workflow was compared with two perturbations:

1. no filter: annotation-quality and spatial ion filters were disabled;
2. random ID: one candidate compound identity was selected at random for each ambiguously annotated ion, replacing the baseline ambiguity-aware allocation strategy.

The analysis included 22 MSI datasets. Results were assessed using the four metrics defined in the Methods section of the main manuscript: region-versus-rest effect size, Moran’s *I*, ambiguity propagation, and number of significant molecular sets. Canonical pathways and *de novo* molecular families were analyzed separately (Supplementary Figs. 2 and 3).

#### S2.2 Results of the perturbation analysis

Across datasets, removal of ion filtering generally reduced the contrast of region-associated molecular-set scores and often decreased spatial autocorrelation, with the strongest effects observed for *de novo* molecular families in several datasets (Supplementary Figs. 2 and 3). In parallel, the number of nominally significant canonical pathways frequently increased under the no filter perturbation despite weaker effect sizes or spatial structure. This pattern is consistent with the inclusion of poorly supported or spatially unstructured ions increasing the number of detectable associations without a corresponding improvement in spatial coherence.

The random ID perturbation produced a different failure mode. Hard assignment of one candidate identity per ambiguous ion generally reduced the number of significant canonical pathways and molecular families and altered region-level effect-size distributions, with the magnitude of the effect varying substantially across datasets. As expected from the construction of the perturbation, ambiguity-propagation scores for molecular families collapsed toward zero because annotation ambiguity had been removed artificially rather than resolved by additional molecular evidence.

The response to both perturbations was dataset dependent. Some datasets retained substantial spatial structure under all configurations, whereas others showed marked loss of region-level contrast or Moran’s *I*. The small-molecule MALDI datasets were among the least stable, consistent with the lower number of lipid-like assignments available for molecular-family construction.

**Supplementary Fig. 2.**
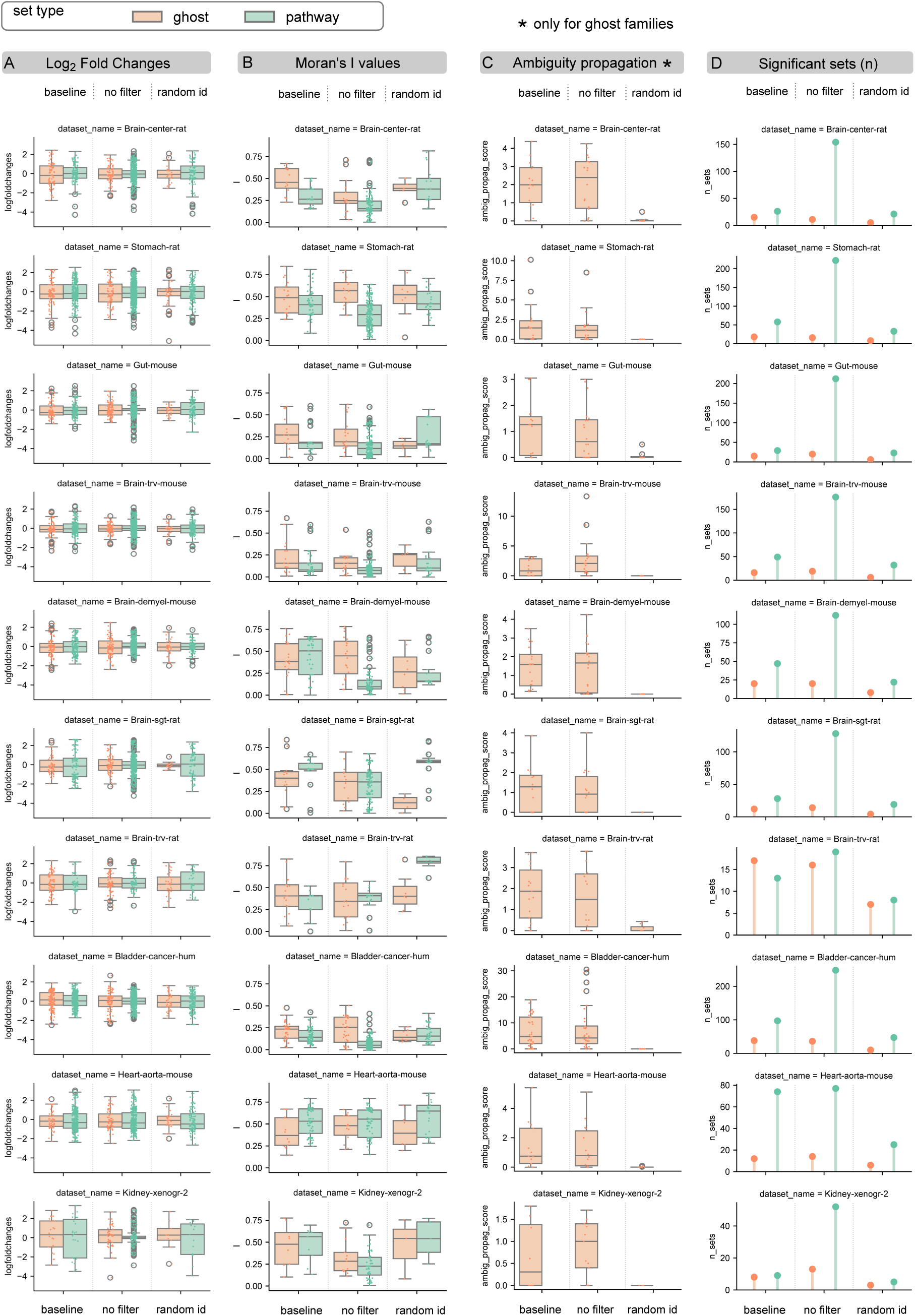
Systematic perturbation analysis of SpacePath. Results for the first 10 datasets are shown for the baseline workflow and the no filter and random ID perturbations. Columns show: **A,** region-versus-rest effect sizes for molecular-set/region comparisons; **B,** Moran’s *I* of spot-wise molecular-set score maps; **C,** ambiguity-propagation scores for *de novo* molecular families; and **D,** the number of molecular sets passing the adjusted significance threshold (*P*_adj_ ≤ 0.05). Canonical pathways and *de novo* molecular families are displayed separately. Box plots show the median and interquartile range; points in A represent molecular-set/region comparisons, whereas points in B and C represent individual molecular sets.

**Supplementary Fig. 3.**
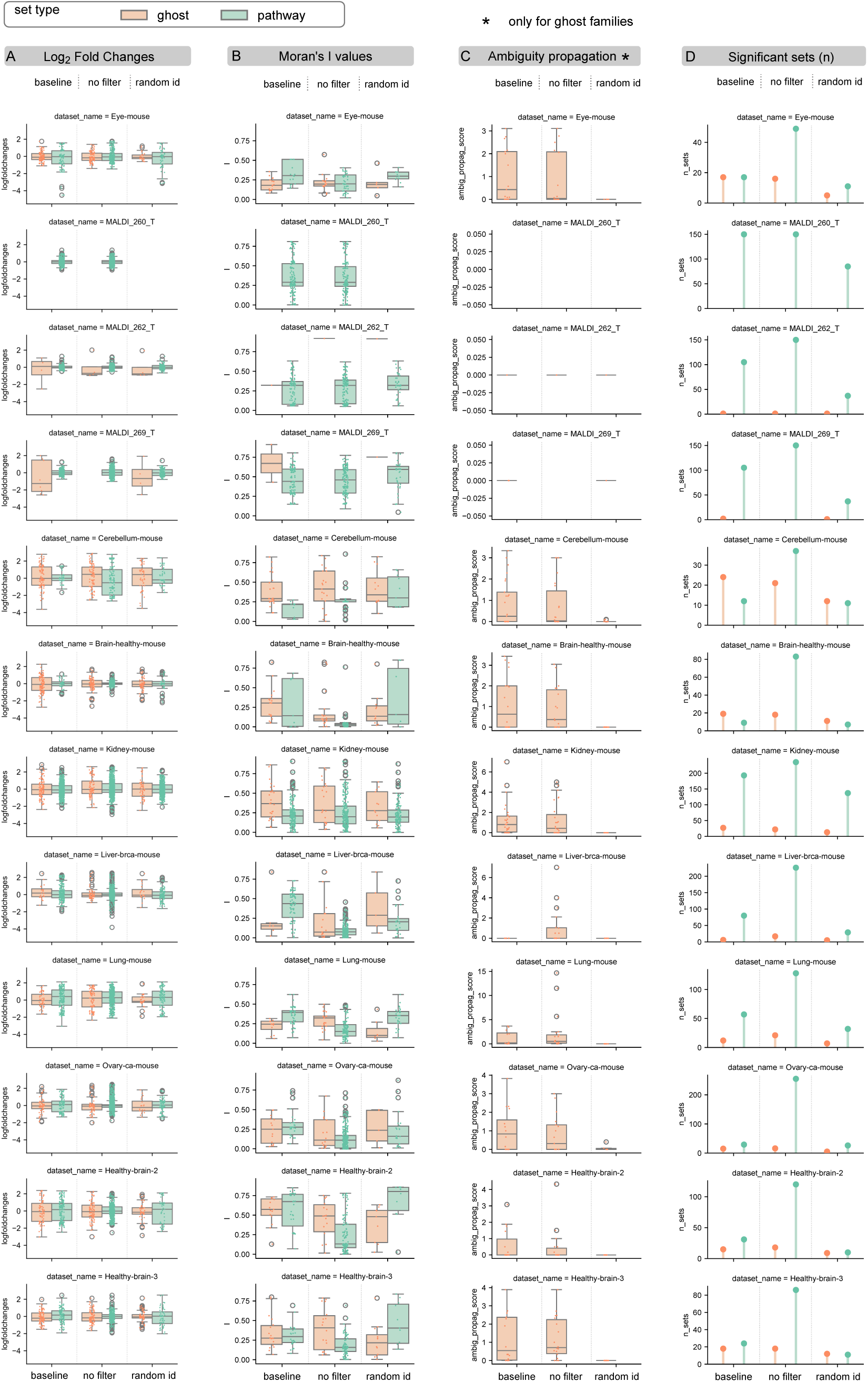
Systematic perturbation analysis of SpacePath, continued. Results for the remaining 12 datasets are displayed using the same metrics and conventions as in Supplementary Fig. 2. The three MALDI 260 T, MALDI 262 T and MALDI 269 T datasets correspond to small-molecule acquisitions and showed weaker performance across one or more configurations.

### S3 Supplementary details of public-dataset benchmarking

The main manuscript reports the principal concordance results for four public MSI datasets with independently described biological structure: healthy mouse brain, cuprizone-associated demyelination, healthy mouse cerebellum, and muscle-invasive bladder cancer. This section provides additional ion-level and molecular-set results for the healthy-brain, cerebellum, and bladder-cancer benchmarks. The demyelination benchmark is described in full in the main Results and is not repeated here.

#### S3.1 Healthy mouse brain

The healthy-brain lipidomics acquisition from Clarke et al. [13] was re-annotated with METASPACE and analyzed with SpacePath as described in the main Methods. Supplementary Fig. 4 provides direct ion-level comparisons with selected lipid distributions reported by Clarke et al., whereas Supplementary Fig. 5 shows the additional decomposition of the *Glycerophospholipid metabolism* pathway into its contributing ions.

Several ion-level correspondences provide additional detail beyond the main Results:

- Phosphatidylserines: high-mass phosphatidylserine species showed preferential localization to white matter, consistent with the PS distributions highlighted by Clarke et al. (Supplementary Fig. 4C-i).
- *m/z* 890.639: this SpacePath feature, assigned to a galactosylceramide species and represented in a *de novo* molecular family, localized to the same white-matter region as the closely matched unannotated *m/z* 890.663 feature reported by Clarke et al. (Supplementary Fig. 4C-ii).
- *m/z* 862.608: the feature assigned to a sulfogalactosylceramide and represented in *Sphingolipid metabolism* reproduced the localization of the unannotated *m/z* 862.606 feature reported in the source study (Supplementary Fig. 4C-iii).
- *m/z* 834.529: SpacePath recovered this ion, reported in the supplementary material of Clarke et al., with a candidate PS(40:6) assignment. The signal was concentrated in cortical and hippocampal regions and contributed to several retained canonical sets, including *Glycerophospholipid metabolism* (Supplementary Fig. 4C-iv).

Beyond the ion-level matches summarized above, *Glycerophospholipid metabolism* showed spatial differentiation across the healthy-brain section, with higher pathway scores in cortical regions (clusters 1 and 3) and the amygdala-associated region (cluster 4). Supplementary Fig. 5 shows the six displayed ions contributing to this molecular-set signal and their individual spatial distributions.

#### S3.2 Healthy mouse cerebellum

The cerebellum dataset from Bond et al. [11] provides an ion-level benchmark because the source study reported phosphatidylcholine distributions for the principal cerebellar layers. SpacePath resolved the molecular, granular, and white-matter compartments, with one additional non-tissue cluster treated as background, as described in the main Results.

The three phosphatidylcholine markers emphasized by Bond et al. were recovered in the corresponding anatomical compartments:

- PC(36:1) in white matter (cluster 2; Supplementary Fig. 6C);
- PC(38:6) in the granular layer (cluster 0; Supplementary Fig. 6D);
- PC(40:6) in the molecular layer (cluster 3; Supplementary Fig. 6E).

These assignments were obtained independently through METASPACE rather than the massPix annotation used in the source study. PC(38:6) remained informative at the individual-ion level but did not satisfy the support requirements for inclusion in a retained canonical pathway or *de novo* molecular family. This illustrates that SpacePath preserves region-associated ions even when they do not contribute to a higher-level molecular set.

The expanded molecular-set representation in Supplementary Fig. 6F additionally shows white-matter associations with *Sphingolipid metabolism* and an oligodendrocyte-differentiation pathway, together with chemically related long-chain phospholipid families. These findings complement the molecular-layer, granular-layer, and white-matter programs summarized in the main Results.

**Supplementary Fig. 4.**
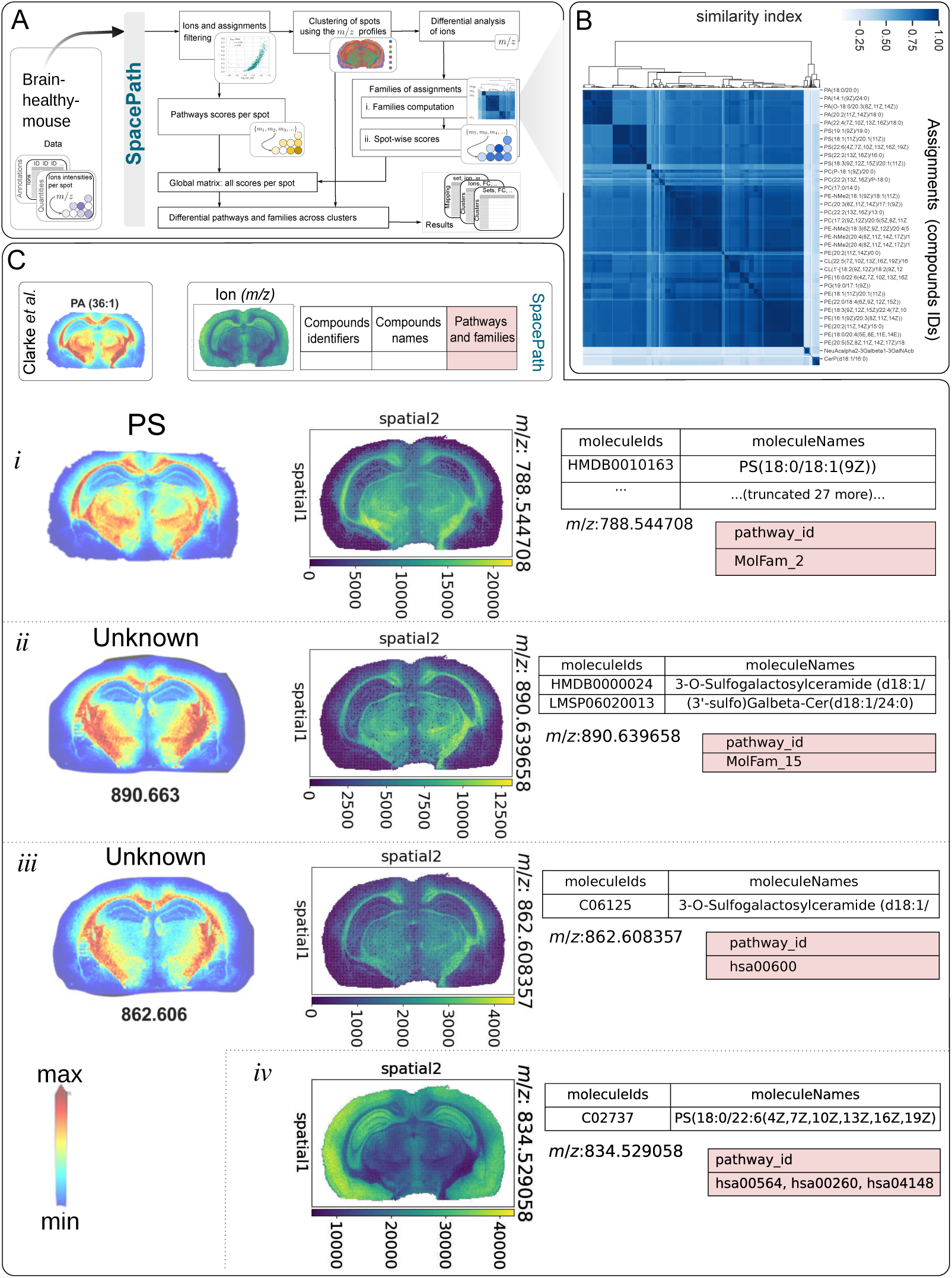
A) SpacePath analysis of the healthy-mouse-brain lipidomics dataset from Clarke et al. B) Hierarchical organization of candidate molecular assignments as a result of the computation of *de novo* molecular families: for illustration purposes, a dendrogram is shown in the upper side of the Tanimoto similarities matrix, but such dendrogram is computed on the distances matrix. C) Direct comparison between selected lipid distributions reported by Clarke et al. and corresponding SpacePath results.

#### S3.3 Muscle-invasive bladder cancer

The muscle-invasive bladder cancer dataset from Tu et al. [7] was used to compare SpacePath with molecular distributions reported in the source study. Whereas Tu et al. primarily compared pooled tumor and non-tumor tissue, SpacePath analyzed spatial heterogeneity within a single annotated cancer section. The resulting regions and representative ion-level correspondences are shown in Supplementary Fig. 7.

**Supplementary Fig. 5.**
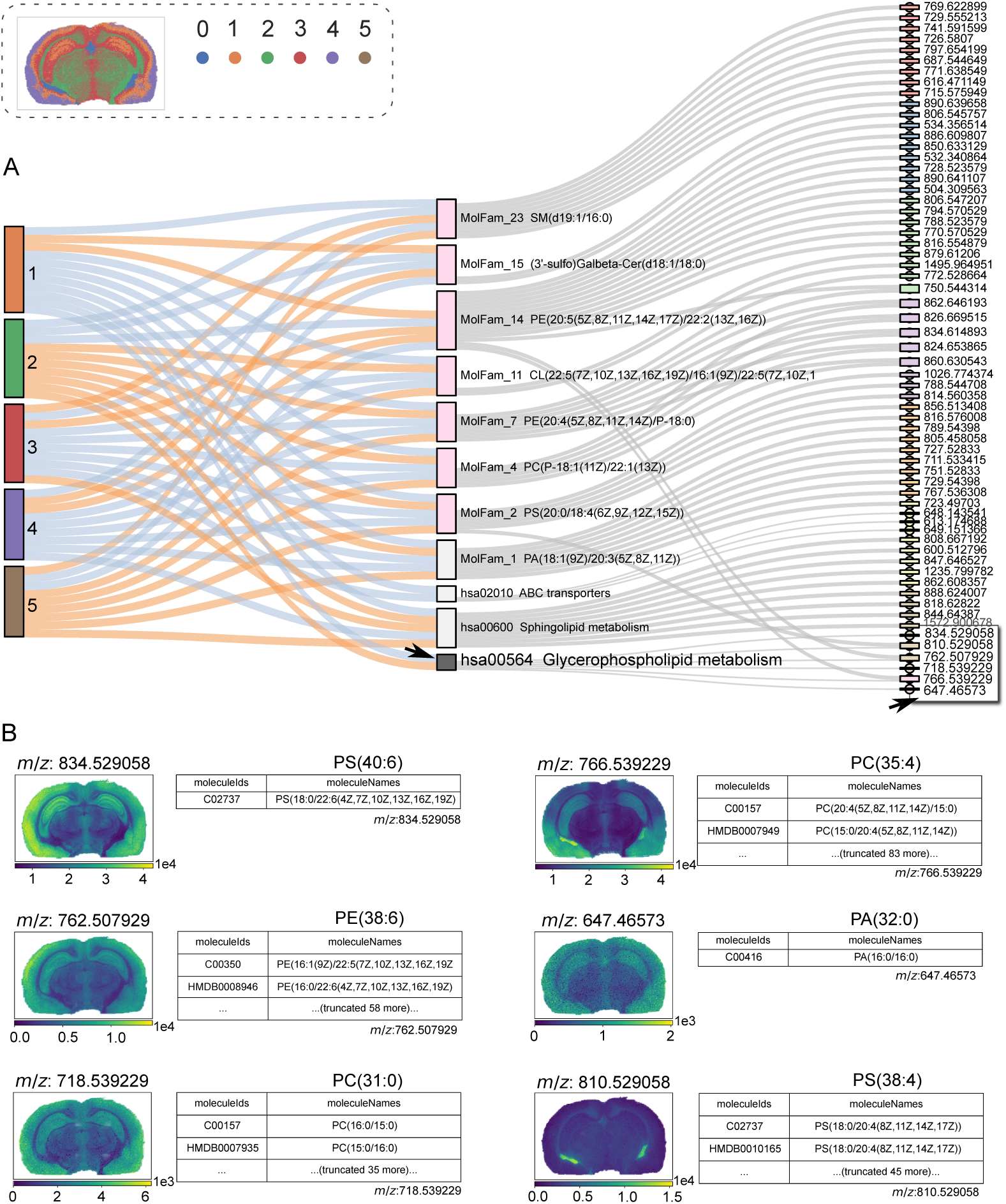
Additional detail for the *Glycerophospholipid metabolism* pathway in the healthy mouse brain. A) Alluvial representation linking spatial regions, molecular sets, and selected contributing ions; the pathway and its contributing ions are indicated by arrows. B) Spatial distributions and candidate assignments of the six displayed ions contributing to this pathway.

Three ion-level observations provide additional detail beyond the main Results:

- PE(32:1) showed preferential signal in the tumor-associated region (Supplementary Fig. 7C-i). Although this candidate compound maps to KEGG pathways, the corresponding molecular sets did not satisfy the SpacePath support requirements in this section.
- PC(P-34:1) likewise localized preferentially to tumor-associated tissue, consistent with the source study (Supplementary Fig. 7C-ii).
- TG(54:3) was concentrated in the smooth-muscle-associated region within the section (Supplementary Fig. 7C-iii).

Thus, individual ions can remain informative even when they do not contribute to a retained molecular set because fewer than two distinct ion-supported assignments are available.

The expanded analysis in Supplementary Fig. 8 provides several additional region-associated signals. A diacylglycerol-rich molecular family was associated with the tumor core, consistent with the increased DG/MG abundance reported by Tu et al. The smooth-muscle region showed higher *Fatty acid biosynthesis* scores, while a high-carbon-number triacylglycerol family (MolFam 11) containing TG(58:7) was associated with peri-tumoral or non-tumor regions. These observations complement the representative fatty-acid and diglyceride signals reported in the main Results.

**Supplementary Fig. 6.**
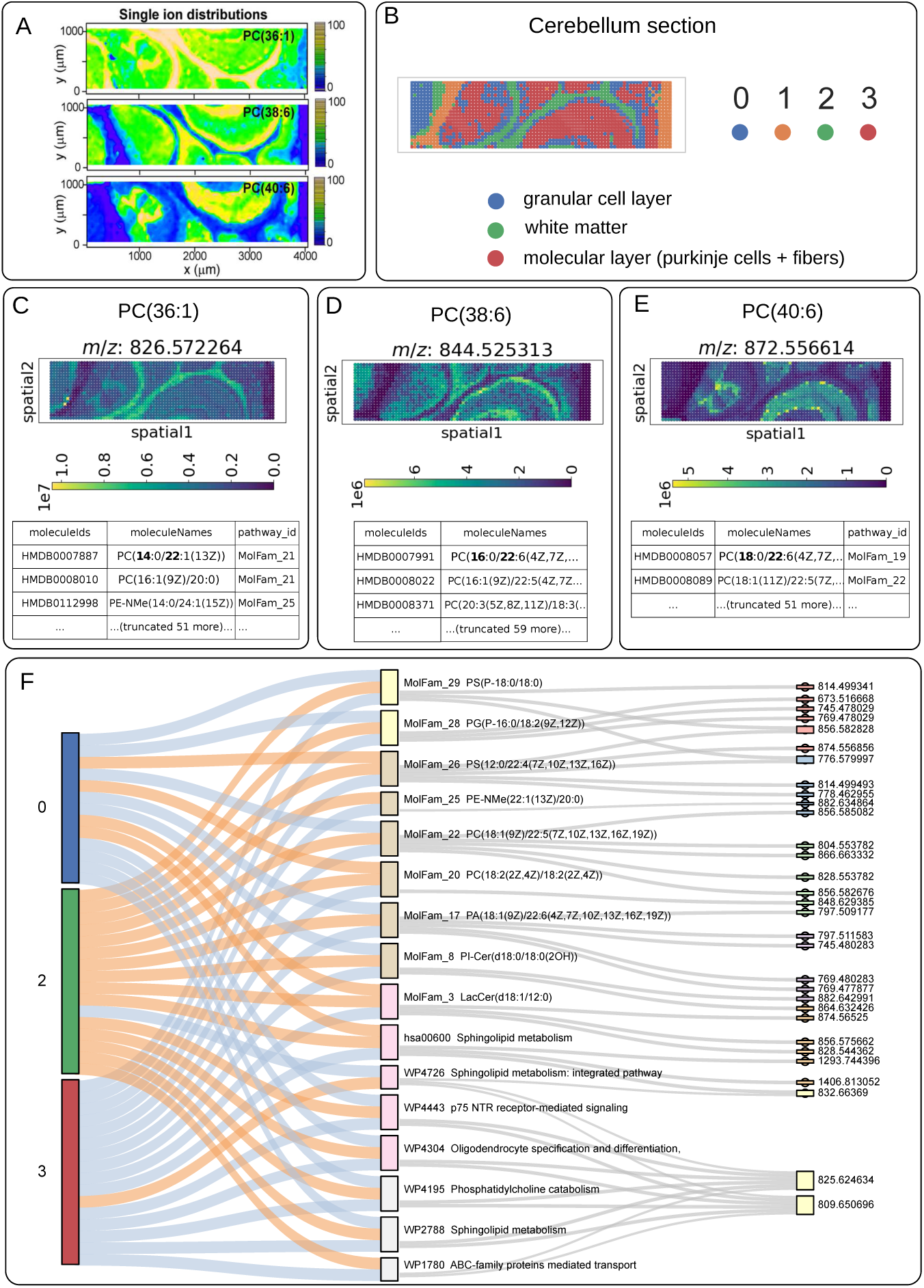
Benchmarking on healthy mouse cerebellum. A) Original ion maps from Bond et al. B) Tissue segmentation by *k*-means (*k* = 4), including one non-tissue background cluster. C–E) Spatial distributions of phosphatidylcholine features recovered by SpacePath. F) Expanded alluvial representation linking spatial regions, significant canonical pathways and *de novo* molecular families, and contributing ions.

Additional canonical-set associations included wax/plasmalogen-related signals and the *Alphalinolenic acid metabolism* and *Sebaleic acid metabolism* sets (Supplementary Fig. 8C). These associations were not reported in the source study and are therefore treated as dataset-specific candidate observations rather than as independently validated bladder-cancer programs.

### S4 Supplementary analysis of study-generated glioblastoma xenografts

The main Results report the principal SpacePath findings for the P3-intact and P3-post-resection xenografts. This section provides additional ion- and molecular-set associations for these tissues and presents the analysis of the two xenografts from the conditional *Mct1* cohort.

**Supplementary Fig. 7.**
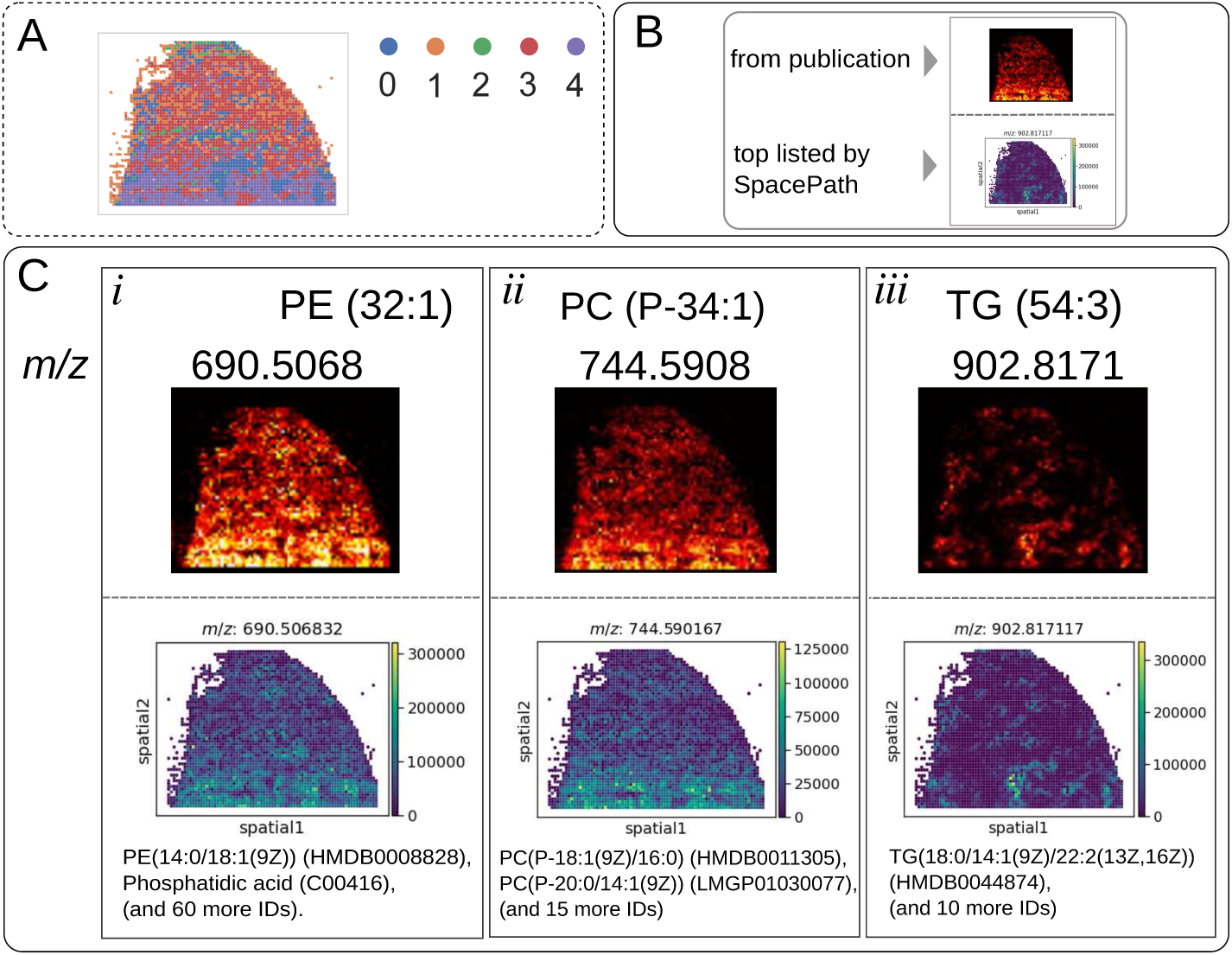
Spatial distribution of selected ions in the bladder-cancer benchmark. A) Spatial regions identified in the analyzed tissue section. B) Legend for panel C. C) Ion distributions reported by Tu et al. (top) and corresponding features recovered by SpacePath (bottom).

**Supplementary Table 3.**
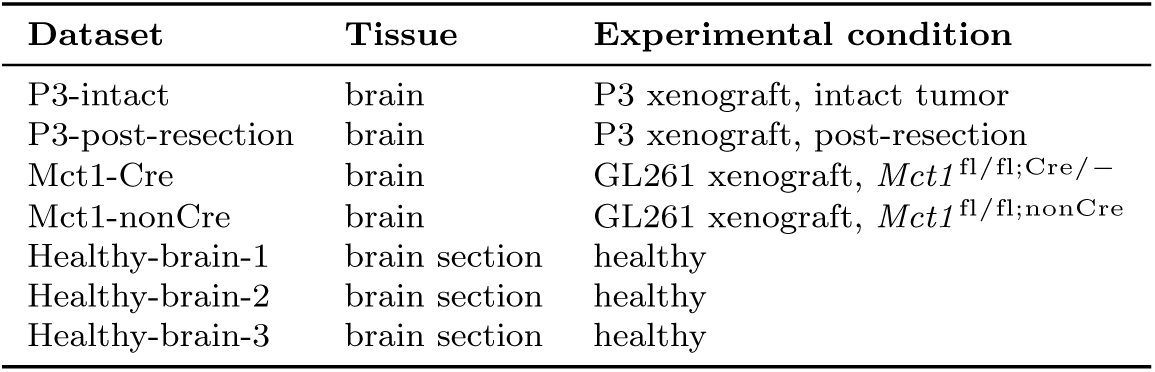
Spatial lipidomics datasets generated in this study. Four glioblastoma xenograft tissues and three healthy mouse brain sections were acquired. Healthy-brain-1 was excluded from the cross-dataset analyses after application of the dataset-level inclusion criteria described in the main Methods. The four xenograft datasets are analyzed in this section.

| **Dataset** | **Tissue** | **Experimental condition** |
|---|---|---|
| P3-intact | brain | P3 xenograft, intact tumor |
| P3-post-resection | brain | P3 xenograft, post-resection |
| Mct1-Cre | brain | GL261 xenograft, *Mct1*<sup>fl/fl;Cre/−</sup> |
| Mct1-nonCre | brain | GL261 xenograft, *Mct1*<sup>fl/fl;nonCre</sup> |
| Healthy-brain-1 | brain section | healthy |
| Healthy-brain-2 | brain section | healthy |
| Healthy-brain-3 | brain section | healthy |

#### S4.1 P3 xenografts

##### S4.1.1 P3-intact xenograft

Supplementary Fig. 9 extends the P3-intact analysis shown in the main manuscript by displaying associations for all six spatial regions and additional representative ions.

Several additional ion-level patterns were not included in the main figure. The feature at *m/z* 870.540963, assigned to PC(40:7) (HMDB0008090) and represented in MolFam 27, showed lower signal in the tumor and preferential localization to cortical and striatal regions (Supplementary Fig. 9B). LysoPE(20:4), represented in MolFam 36, was associated with the pia-mater region and also detected in the striatal compartment, whereas a glucosylceramide-associated signal localized preferentially to white matter.

The feature at *m/z* 725.556796, assigned to SM(d34:1), showed higher signal in the tumor and mapped to the WikiPathways set *Sphingolipid metabolism integrated pathway* (WP4726). This association is not displayed in the alluvial summary because only a subset of statistically ranked overlapping sets is shown. The example illustrates that one candidate compound can contribute to several overlapping canonical sets and that the alluvial representation is a visualization subset rather than an exhaustive pathway inventory.

**Supplementary Fig. 8.**
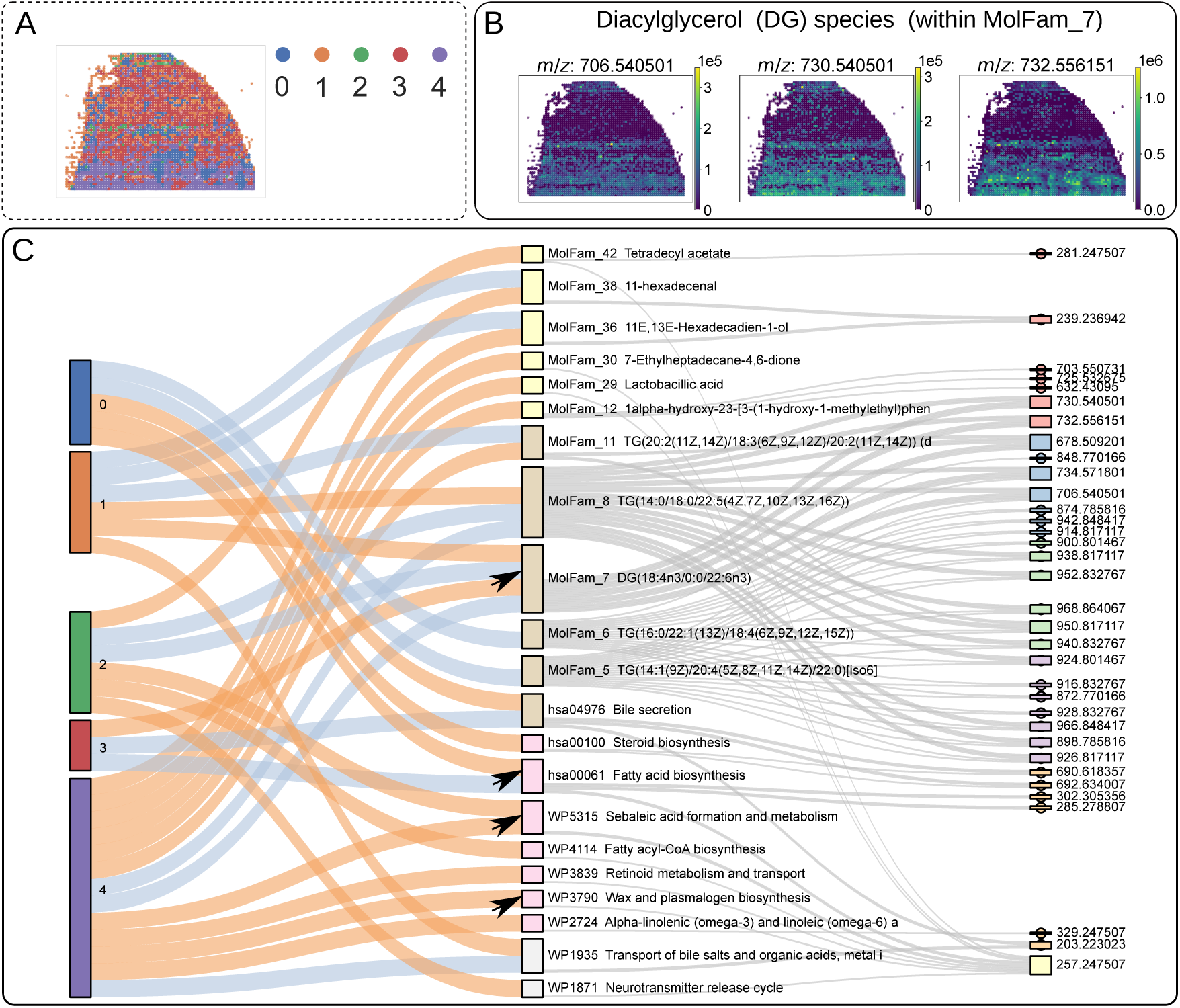
Additional molecular-set results for the bladder-cancer benchmark. A) Spatial regions. B) Representative diacylglycerol distributions in tumor-associated tissue. C) Alluvial summary linking regions, canonical pathways and *de novo* molecular families, and contributing ions. Directionality labels in the original visualization represent relative within-section score differences and should not be interpreted as direct measurements of pathway activity. Arrows indicate associations discussed in the text.

SpacePath also identified tumor-associated ions whose corresponding molecular families did not reach the molecular-set significance threshold. In particular, *m/z* 669.446577, assigned to PA(32:1) and represented in MolFam 33, was significant in the ion-level region-versus-rest analysis despite MolFam 33 itself not passing the set-level threshold (Supplementary Fig. 9C). This provides an example in which a spatially informative ion is retained even when its higher-level molecular set is not significantly differentiated.

A second tumor-localized feature, *m/z* 854.590561, was assigned by METASPACE to PS(41:4) (LMGP03010478) and represented in MolFam 16 (Supplementary Fig. 9C). The odd total carbon number of this candidate assignment warrants cautious interpretation, and no alternative even-carbon candidate was available in the annotation used here. The spatial association of the ion is therefore distinguishable from certainty about its molecular identity, which would require orthogonal structural evidence.

##### S4.1.2 P3-post-resection xenograft

The main manuscript reports the principal spatial programs in the P3-post-resection section. Supplementary Fig. 10 provides the complete alluvial representation and additional ion-level associations across all seven spatial regions.

A phosphatidylcholine-rich molecular family (MolFam 23) was preferentially associated with gray-matter regions (clusters 4–6; Supplementary Fig. 10B). Additional sphingomyelin-associated signals were observed in the recurrent-tumor region. For example, SM(d42:2) contributed to the canonical *Sphingolipid metabolism integrated pathway* (WP4726).

**Supplementary Fig. 9.**
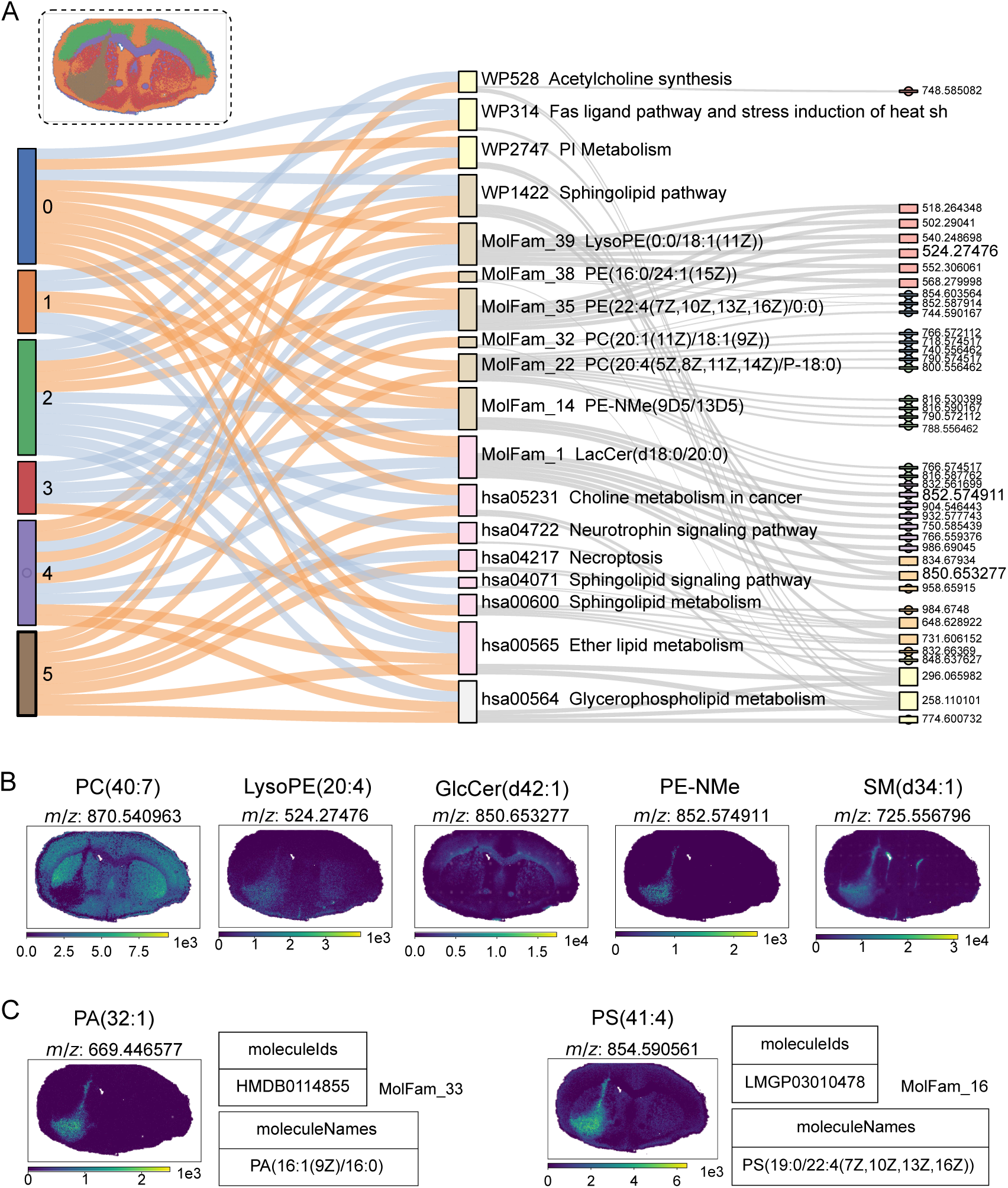
Additional SpacePath results for the P3-intact glioblastoma xenograft. A) Spatial regions and alluvial representation linking regions, canonical pathways and *de novo* molecular families, and contributing ions. All six spatial regions are included. B) Spatial distributions of additional representative ions. C) Examples of ion-level output combining candidate METASPACE assignments with canonical-pathway or molecular-family membership.

The tumor-associated SM(d34:1) assignment provides an additional example of overlapping canonical-set membership: the same candidate compound was represented in *Phosphatidylcholine catabolism* (WP4195), *Oligodendrocyte specification and differentiation* (WP4304), *ABC-family proteins mediated transport* (WP1780), and *Sphingolipid metabolism integrated pathway* (WP4726) (Supplementary Fig. 10B). These multiple memberships reflect overlap among pathway definitions and should not be interpreted as four independent biological signals.

The feature at *m/z* 854.590561 was again associated with the tumor region and carried the PS(41:4) candidate assignment discussed above (Supplementary Fig. 10C). Its recurrence across the two P3 sections supports the spatial reproducibility of the ion-level observation but does not resolve the molecular identity of the feature.

**Supplementary Fig. 10.**
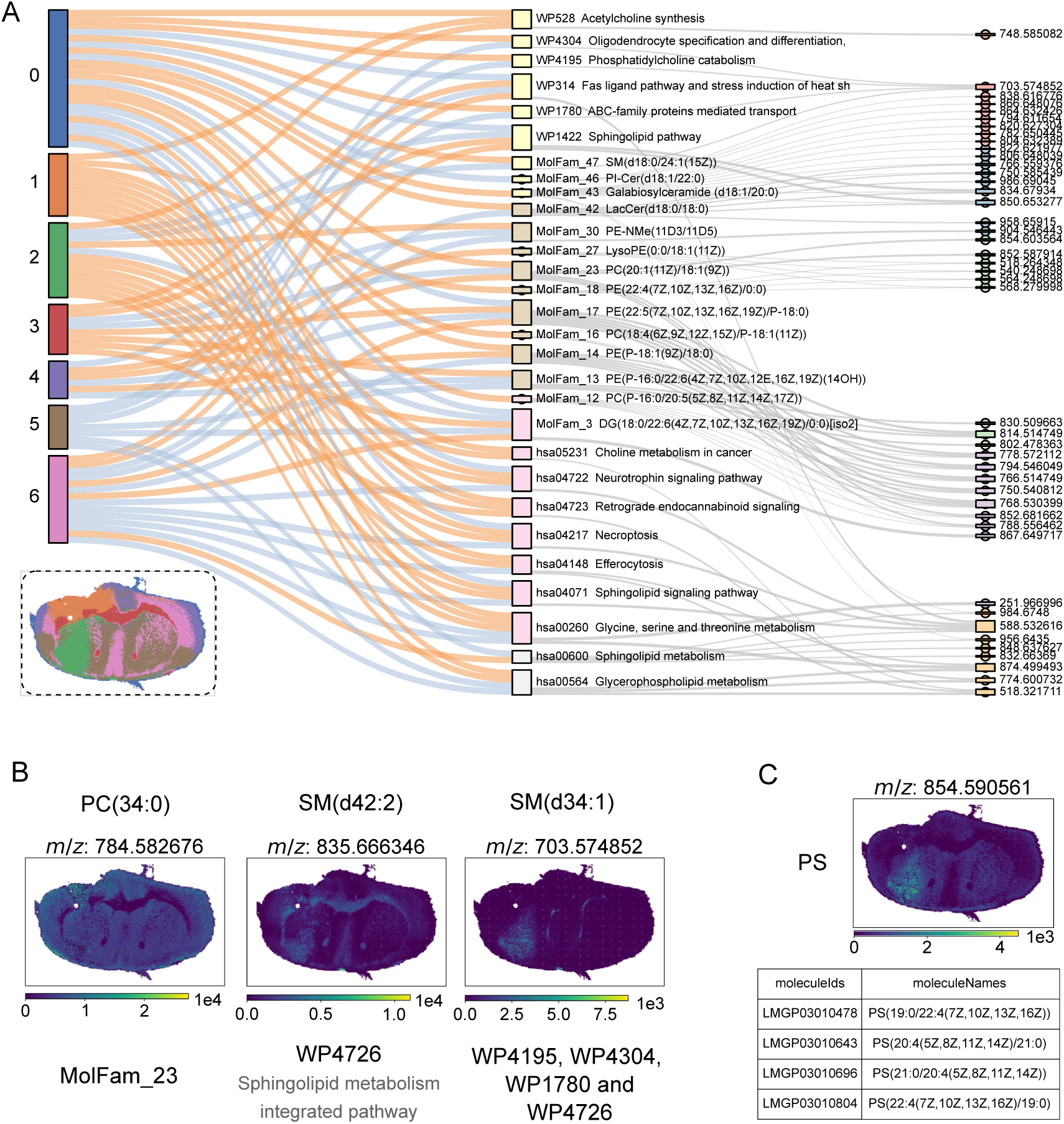
Additional SpacePath results for the P3-post-resection glioblastoma xenograft. A) Complete alluvial representation linking spatial regions, canonical pathways and *de novo* molecular families, and contributing ions. All seven spatial regions are included; the region-color key is shown below. B) Spatial distributions of additional representative ions. C) Example of a candidate molecular assignment requiring cautious interpretation.

#### S4.2 Second cohort: GL261 xenografts in Mct1-Cre and Mct1-nonCre mice

The second cohort comprised two GL261 xenografts, one generated in an *Mct1* ^fl/fl;Cre/−^ mouse and one in an *Mct1* ^fl/fl;nonCre^ mouse (Supplementary Table 3). SpacePath was applied independently to both sections.

In both sections, the tumor-associated region was differentiated from surrounding brain by several molecular-set scores, including *Sphingolipid metabolism*, *Efferocytosis*, and the *Fas ligand pathway* (Supplementary Fig. 11). These observations are within-section spatial associations.

The MSI acquisition used a lateral resolution of 25 µm, which does not isolate macrophage-specific signals. The two sections are retained as additional examples of SpacePath analysis across distinct xenograft backgrounds, but no macrophage-specific mechanistic conclusion is drawn from them.

**Supplementary Fig. 11.**
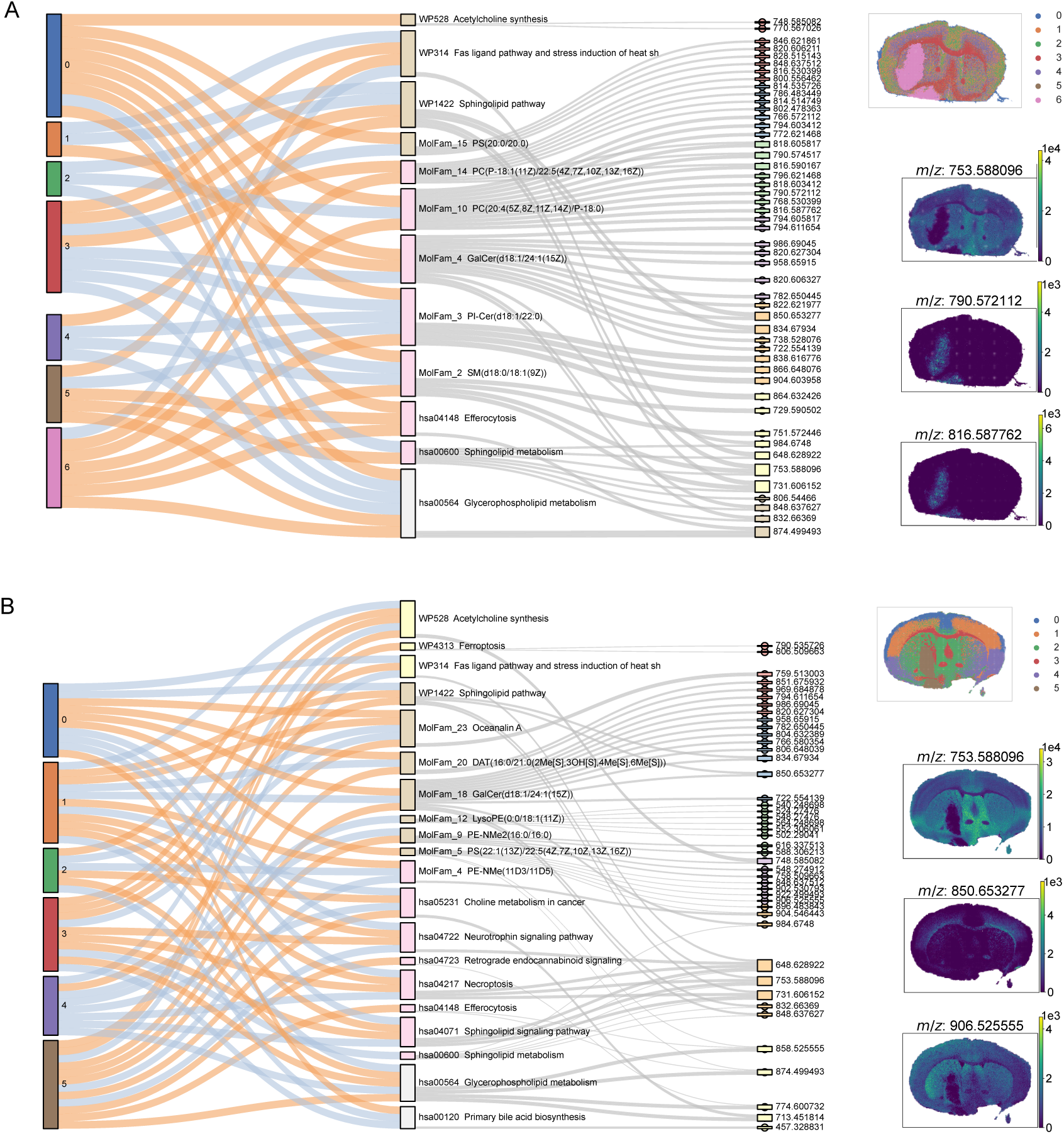
SpacePath analysis of the second glioblastoma xenograft cohort. Global SpacePath outputs are shown for the two GL261 xenograft sections, with all spatial regions included. A) Mct1-Cre. B) Mct1-nonCre.

### S5 Supplementary note: relation to existing chemical-set approaches

The limited representation of metabolites and lipids in canonical pathway databases has motivated several approaches that organize molecular measurements independently of conventional pathway definitions. Metabolomics2Network, for example, uses lipid ontologies to improve the correspondence between experimentally detected lipid species and genome-scale metabolic networks [17]. ChemRICH instead performs enrichment analysis using chemical similarity and ontology-based compound groups, thereby reducing dependence on incomplete biochemical pathway annotations [18]. These approaches established the utility of chemical structure and lipid ontologies for interpreting metabolomics data when canonical pathway mapping is sparse.

SpacePath addresses a related problem in the specific context of spatial MSI, where limited pathway coverage is compounded by uncertainty in MS1-based molecular assignment. Rather than collapsing each ion to a single identity or mapping lipid species directly to broad classes, SpacePath retains compatible candidate assignments and uses their chemical structures to construct dataset-specific *de novo* molecular families for pathway-unmapped compounds. These families are subsequently scored at each spatial location using the same framework as canonical pathways. They therefore provide a chemically defined level of organization for spatially structured signals that cannot be represented by the reference pathway resources used here.

Preserving molecular granularity is particularly relevant for lipidomics. Lipids belonging to the same broad class can differ substantially in acyl-chain length, unsaturation and other structural features that influence membrane organization and molecular interactions. Collapsing species such as PC(36:2) and PC(42:6) into a single phosphatidylcholine class can therefore remove distinctions that remain detectable in MSI. The molecular families used by SpacePath provide a complementary representation in which structural similarity can be captured at a finer scale without requiring each group to correspond to an established biochemical pathway.

These *de novo* families should consequently not be interpreted as inferred pathways or as replacements for curated metabolic resources. Their role is to retain chemically coherent, spatially organized information from the large fraction of MSI assignments that remains outside canonical pathway mappings, and to provide hypothesis-generating molecular sets for subsequent experimental investigation.

### S6 Supplementary Methods

#### S6.1 Compound Identifier Handling and Disambiguation

The processes related to compound identifiers handling, disambiguation and filtering are carried out by the core library of SpacePath. Each process is defined by specific functions that are explained in the Wiki page of the library. Here, the rationale of each step, in the order as they are performed in SpacePath, is provided.

### Compound Identifier Filtering

The compound identifiers filtering is the first step that is applied on the user-provided annotations. It is based on specific criteria to ensure biological relevance: Small odd-chain lipids (with fewer than 17 carbons) are systematically excluded unless explicitly disabled by the user. Additionally, SpacePath is equipped with internal curated lists that are used to filter out identifiers associated with plants, microorganisms, or medications, when these are not relevant the biological context of interest (the user can chose the type(s) to exclude). This filtering step ensures that only high-confidence and contextually appropriate identifiers are retained for downstream analysis. This is performed automatically by the .load() function, with user parameters.

### Multiple m/z to One ID

Multiple *m/z* features can eventually be annotated to the same compound identifier, by the external software. This type of annotation error is infrequent, nevertheless, our library employs a multi-criteria selection process to disambiguate and retain the most reliable signals. First, it checks for the presence of multiple *m/z* values per ID using a grouping operation. Then, it ranks the *m/z*features based on a combination of MSM (Metabolite Signal Matching) scores, off-sample values, FDR (False Discovery Rate), prioritizing features with the highest MSM, lowest offsample, and lowest FDR. After sorting, the top *m/z* features are selected, and only the best two per compound ID are retained for further analysis. This ensures that ambiguous cases are resolved objectively, preserving the most biologically relevant signals.

### Multiple ID to One m/z

When a single *m/z* feature matches multiple compound identifiers, our library retains all plausible annotations as-is. No arbitrary resolution is applied; instead, the ambiguity is preserved, allowing users to manually review or apply additional biological or analytical criteria to refine the annotations. This approach ensures transparency and flexibility in handling inherent uncertainties in mass spectrometry data.

### Alias Resolution Precedence and Deduplication

On the the filtered annotations, the compound identifiers aliases are found and resolved. For instance, HMDB0000032, C01164, and LMST01010069 are three aliases for ‘7-Dehydrocholesterol’, provided in HMDB, KEGG, and LipidMaps nomenclature, respectively. SpacePath is internally equipped with a mapping table of aliases. The library uses this internal information to select, on the dataset annotations, one compound identifier among the aliases, in this strict order: LipidMaps take the highest precedence, followed by HMDB, and finally, KEGG identifiers. Additionnally, CoreMetabolome identifiers can be present in the user-provided and filtered annotations, which correspond to manually curated entities authored by METASPACE developers, conserving the HMDB nomenclature. Therefore, our library maps CoreMetabolome identifiers to the other types of identifiers using the HMDB mappings.

The comprehensive aliases table of HMDB, LipidMaps, and KEGG identifiers was generated with custom scripts available at the public Zenodo registry https://zenodo.org/records/23079160. This constructed aliases table is shipped within SpacePath > SpacePath > resources/ in the package GitHub repository.

