## Supplementary Material for "SpacePath enables pathway and molecular-family analysis beyond annotation ambiguity in spatial metabolomics and lipidomics"

### SpacePath: A computational framework for the functional annotation of spatial metabolomics and lipidomics data
### Supplementary material

J. Galvis, M. Moreau, S. Lillo, D. Bomont, L. Ledoux, Y. Zirem, M. Salzet, I. Fournier, N. Desbenoit, T. Daubon and M. Nikolski

### S1 SpacePath workflow overview

The SpacePath workflow is described in detail in the Methods section of the main manuscript and summarized schematically in Supplementary Fig. 1. Briefly, SpacePath integrates spatial ion-intensity profiles with candidate molecular assignments, applies ion-quality and spatial filtering, and defines spatial regions for downstream comparison. An ambiguity-aware compound representation is then used to score canonical pathways and chemically defined *de novo* molecular families at each spatial location. These scores are combined into a common molecular-set matrix for region-level comparison.

Throughout the Supplementary Information, the term *molecular set* refers collectively to canonical pathways and *de novo* molecular families.

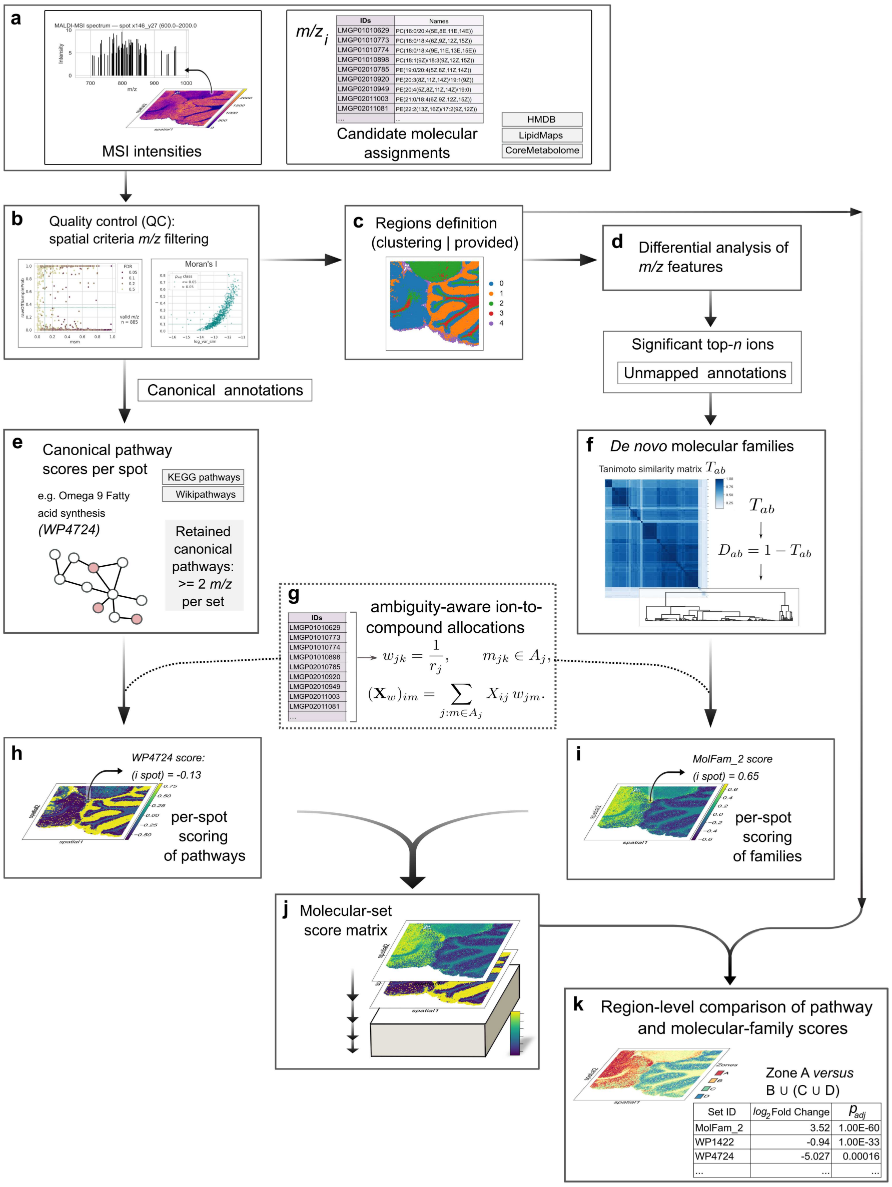

**Supplementary Fig. 1 Overview of the SpacePath workflow.** SpacePath integrates spatial ion-intensity profiles with candidate molecular assignments, applies ion-quality and spatial filtering, defines spatial regions, constructs an ambiguity-aware compound representation, and derives spatial scores for canonical pathways and *de novo* molecular families. These scores are combined into a common molecular-set matrix for region-level comparison.

**Supplementary Table 1** Dataset provenance and selection role

| Dataset name | Metaspace registry | Organism | Organ | Condition | Matrix | IM | Instrument | Spots | Initial ions | Retained ions | Annotation (used) | Included |
|---|---|---|---|---|---|---|---|---|---|---|---|---|
| Brain-center-rat | 2025-04-11_11h55m02s | rat | brain | WT | DHB | + | orbitrap | 7034 | 2789 | 249 | HMDB+LM | yes |
| Stomach-rat [1] | 2017-11-09_11h42m17s | rat | stomach | WT | DHB | + | orbitrap | 11240 | 5659 | 878 | HMDB+LM | yes |
| Gut-mouse | 2022-07-06_09h33m52s | mouse | gut | WT | SDHB | + | orbitrap | 50568 | 5642 | 1056 | HMDB+LM | yes |
| Brain-wt1-mouse | 2025-04-10_23h29m14s | mouse | brain | WT | CHCA | + | FT-ICR | 10916 | 8458 | 711 | HMDB+LM+CM | yes |
| Brain-sgt-mouse [2] | 2025-04-10_14h45m11s | mouse | brain | WT | DHB | + | orbitrap | 3414 | 1767 | 180 | HMDB+LM+CM | yes |
| Brain-trv-mouse [3] | 2020-01-16_13h13m47s | mouse | brain | WT | DHB | + | orbitrap | 32816 | 2666 | 376 | HMDB+LM | yes |
| Brain-demyel-mouse [4] | 2023-07-09_23h26m29s | mouse | brain | WT | CHCA | + | orbitrap | 78760 | 2596 | 778 | HMDB+LM | yes |
| Brain-sgt-rat [5] | 2025-04-11_12h18m07s | rat | brain | WT | Nh | + | orbitrap | 24130 | 4298 | 731 | HMDB+LM | yes |
| Brain-trv-rat [6] | 2025-04-10_15h11m23s | mouse | brain | WT | Nh | + | FT-ICR | 21455 | 1220 | 542 | HMDB+LM | yes |
| Bladder-cancer-hum [7] | 2021-01-04_01h15m46s | human | bladder | cancer | none† | + | orbitrap | 3275 | 6534 | 804 | HMDB+LM | yes |
| Heart-aorta-mouse [7] | 2021-12-06_14h55m46s | mouse | heart | WT | NEDC | - | FT-ICR | 1863 | 411 | 230 | HMDB+LM+CM | yes |
| Heart-sgt-mouse [8] | 2023-12-18_17h45m47s | mouse | heart | WT | NEDC | - | orbitrap | 34604 | 3460 | 323 | HMDB+LM+CM | yes |
| Kidney-xenogr-1 [9] | 2019-04-20_01h02m45s | xenograft | kidney | WT | 9-AA | + | orbitrap | 4301 | 272 | 82 | HMDB+LM | no |
| Kidney-xenogr-2 [9] | 2019-04-20_00h55m20s | xenograft | kidney | WT | 9-AA | + | orbitrap | 12667 | 1265 | 254 | HMDB+LM | yes |
| Eye-mouse | 2018-03-20_11h02m36s | mouse | eye | WT | DAN | + | orbitrap | 12124 | 1876 | 621 | HMDB+LM | yes |
| MALDI_248_T [10] | not provided | human | brain | cancer | 9-AA | - | FT-ICR | 11677 | 875 | 613 | HMDB | yes |
| MALDI_259_T [10] | not provided | human | brain | cancer | 9-AA | - | FT-ICR | 16943 | 875 | 203 | HMDB | yes |
| MALDI_260_T [10] | not provided | human | brain | cancer | 9-AA | - | FT-ICR | 17334 | 875 | 613 | HMDB | yes |
| MALDI_262_T [10] | not provided | human | brain | cancer | 9-AA | - | FT-ICR | 10443 | 875 | 203 | HMDB | yes |
| MALDI_269_T [10] | not provided | human | brain | cancer | 9-AA | - | FT-ICR | 13622 | 875 | 203 | HMDB | yes |
| MALDI_275_T [10] | not provided | human | brain | cancer | 9-AA | - | FT-ICR | 19714 | 875 | 613 | HMDB | yes |
| Cerebellum-mouse [11] | 2025-04-10_14h53m38s | rat | cerebellum | WT | THAP | + | orbitrap | 1701 | 1753 | 358 | HMDB+LM | yes |
| Brain-sgt-hr-mouse [12] | 2023-09-29_18h44m00s | mouse | brain | WT | CHCA | + | orbitrap | 91871 | 25950 | 835 | HMDB+LM | yes |
| Brain-healthy-mouse [13] | 2025-07-30_17h59m25s | mouse | brain | WT | NEDC | - | timsTOF | 22982 | 1037 | 191 | HMDB+LM+CM | yes |
| Brain-lmrange-mouse [13] | 2025-07-30_17h35m47s | mouse | brain | WT | NEDC | - | timsTOF | 25054 | 1344 | 32 | KEGG+HMDB+LM | no |
| Kidney-mouse [8] | 2024-09-28_15h43m16s | mouse | kidney | WT | NEDC | - | orbitrap | 36528 | 3230 | 737 | HMDB+LM+CM | yes |
| Liver-brca-mouse [14] | 2023-04-04_07h50m38s | xenograft | liver | cancer | NEDC | - | orbitrap | 38904 | 3382 | 130 | HMDB+CM | yes |
| Lung-mouse [15] | 2019-01-17_19h54m30s | mouse | lung | WT | DAN | - | orbitrap | 7962 | 1484 | 283 | HMDB+LM | yes |
| Ovary-ca-mouse [16] | 2022-04-04_22h54m13s | xenograft | ovary | cancer | DHB | + | FT-ICR | 24132 | 4811 | 356 | HMDB+LM+CM | yes |
| P3-intact | 2025-05-26_11h43m25s | xenograft | brain | cancer | DHB | + | orbitrap | 63310 | 3246 | 669 | HMDB+LM+CM | yes |
| P3-post-resection | 2025-07-30_16h54m35s | xenograft | brain | cancer | DHB | + | orbitrap | 57196 | 2889 | 682 | HMDB+LM+CM | yes |
| Mct-Cre | 2025-05-29_14h37m38s | xenograft | brain | cancer | DHB | + | orbitrap | 63091 | 3171 | 657 | HMDB+LM+CM | yes |
| Mct-nonCre | 2025-05-28_16h28m09s | xenograft | brain | cancer | DHB | + | orbitrap | 59613 | 2983 | 718 | HMDB+LM+CM | yes |
| Healty-brain-1 | 2025-07-28_15h11m01s | mouse | brain | WT | DHB | + | orbitrap | 4947 | 1110 | 116 | HMDB+LM | no |
| Healty-brain-2 | 2025-07-28_15h09m59s | mouse | brain | WT | DAN | - | orbitrap | 28596 | 1060 | 264 | HMDB+LM | yes |
| Healty-brain-3 | 2025-07-28_15h08m37s | mouse | hindbrain | WT | DAN | - | orbitrap | 15918 | 1179 | 240 | HMDB+LM | yes |

When no journal article is associated or the data are sourced from this study, no reference is marked. The last seven datasets are sourced from this study.
*Abbreviations:* 9-AA: 9-Aminoacridine. CHCA: $\alpha$Cyano-4-hydroxycinnamic Acid. CM: CoreMetabolome database. DAN: 1,5-Diaminonaphthalene. DHB: 2,5-Dihydroxybenzoic Acid. FT-ICR: Fourier-Transform Ion Cyclotron Resonance. LM: LIPID MAPS. NEDC: N-(1-naphthyl)ethylenediamine dihydrochloride. Nh: Norharmane (9H-Pyrido[3,4-b]indole). SDHB: Sinapic Acid. THAP: 2,4,6-Trihydroxyacetophenone. timsTOF: Trapped Ion Mobility Spectrometry Time-of-Flight. WT: Wild Type.
†The ionization source was IRMALDESI (Ion Rapid Mass Desorption and Laser Desorption Ionization) which is matrix-free.

#### S1.1 Dataset provenance and analysis inclusion

The provenance and analytical role of all public and study-generated datasets are summarized in Supplementary Table 1. Public datasets were assembled from METASPACE, the associated literature, and other specified sources (e.g. MetaboLights), whereas the study-generated datasets comprised four glioblastoma xenografts and three healthy-brain sections. Dataset inclusion criteria and the distinction between the cross-dataset, benchmarking, glioblastoma, perturbation, and MS/MS analysis sets are described in the main Methods.

#### S1.2 Dataset-specific SpacePath parameters

Dataset-specific SpacePath settings are summarized in Supplementary Table 2. Parameters that were constant across analyses are reported in the main Methods and are not repeated here. The table therefore focuses on settings that varied between datasets, including ion-filtering thresholds, spatial region definition, clustering parameters and molecular-family construction settings used for each analysis.

**Supplementary Table 2 Dataset-specific SpacePath parameters.** Only parameters that varied between datasets are shown; invariant settings are described in the main Methods.

| | Thresholds for Ions Filtering | | | | Spatial Clustering | | Fam. Comp | | Families Output | | |
|---|---|---|---|---|---|---|---|---|---|---|---|
| **Dataset name** | Off Prob | MSM | Moran's I | $p_{adj}$ | Clustering Method | Param | Tree Cut | Min Size | Built Fam. | Scored Fam. | Signif. Fam. |
| Brain-center-rat | 0.2 | 0.2 | 0.1 | 0.05 | $k$-means | 5 | 0.5 | 4 | 30 | 14 | 14 |
| Stomach-rat | 0.2 | 0.25 | 0.1 | 0.05 | $k$-means | 5 | 0.9 | 4 | 35 | 19 | 14 |
| Gut-mouse | 0.2 | 0.5 | 0.1 | 0.05 | $k$-means | 5 | 0.7 | 4 | 25 | 15 | 7 |
| Brain-wt1-mouse | 0.2 | 0.5 | 0.2 | 0.05 | $k$-means | 5 | 1.0 | 4 | 27 | 17 | 10 |
| Brain-sgt-mouse | 0.2 | 0.2 | 0.1 | 0.05 | $k$-means | 5 | 1.0 | 4 | 12 | 11 | 3 |
| Brain-trv-mouse | 0.25 | 0.15 | 0.1 | 0.05 | $k$-means | 5 | 0.7 | 4 | 25 | 16 | 4 |
| Brain-demyel-mouse | 0.2 | 0.4 | 0.1 | 0.05 | $k$-means | 5 | 0.5 | 4 | 34 | 20 | 13 |
| Brain-sgt-rat | 0.2 | 0.5 | 0.1 | 0.05 | $k$-means | 5 | 1.0 | 4 | 20 | 12 | 6 |
| Brain-trv-rat | 0.2 | 0.15 | 0.1 | 0.05 | $k$-means | 5 | 0.5 | 3 | 22 | 17 | 10 |
| Bladder-cancer-hum | 0.2 | 0.15 | 0.05 | 0.05 | Leiden | 0.27 | 0.9 | 2 | 50 | 45 | 16 |
| Heart-aorta-mouse | 0.25 | 0.15 | 0.1 | 0.05 | $k$-means | 5 | 0.9 | 4 | 26 | 12 | 10 |
| Heart-sgt-mouse | 0.2 | 0.4 | 0.2 | 0.05 | Leiden | 0.5 | 1.1 | 3 | 39 | 28 | 19 |
| Kidney-xenogr-1 | 0.55 | 0.1 | 0.1 | 0.05 | Leiden | 0.4 | 1.0 | 3 | 9 | 6 | 4 |
| Kidney-xenogr-2 | 0.45 | 0.1 | 0.1 | 0.05 | Leiden | 0.2 | 1.0 | 2 | 11 | 7 | 4 |
| Eye-mouse | 0.3 | 0.1 | 0.1 | 0.05 | Leiden | 0.4 | 1.0 | 3 | 19 | 14 | 7 |
| MALDI_248_T | 0.85 | 0.0 | 0.001 | 1.0 | $k$-means | 4 | 0.4 | 2 | 11 | 1 | 1 |
| MALDI_259_T | 0.85 | 0.1 | 0.0 | 0.05 | $k$-means | 4 | 0.5 | 2 | 1 | 1 | 1 |
| MALDI_260_T | 0.85 | 0.0 | 0.0 | 1.0 | $k$-means | 4 | 0.4 | 2 | 2 | 0 | 0 |
| MALDI_262_T | 0.85 | 0.1 | 0.0 | 0.05 | $k$-means | 4 | 0.5 | 2 | 4 | 1 | 1 |
| MALDI_269_T | 0.85 | 0.1 | 0.0 | 0.05 | $k$-means | 4 | 0.5 | 2 | 5 | 2 | 2 |
| MALDI_275_T | 0.85 | 0.0 | 0.001 | 1.0 | $k$-means | 4 | 0.4 | 2 | 2 | 1 | 1 |
| Cerebellum-mouse | 0.97 | 0.3 | 0.08 | 0.1 | $k$-means | 4 | 0.5 | 2 | 29 | 24 | 21 |
| Brain-sgt-hr-mouse | 0.15 | 0.5 | 0.2 | 0.1 | $k$-means | 5 | 0.8 | 4 | 31 | 21 | 18 |
| Brain-healthy-mouse | 0.4 | 0.002 | 0.1 | 0.1 | $k$-means | 6 | 0.5 | 2 | 24 | 17 | 10 |
| Brain-lmrange-mouse | 0.35 | 0.1 | 0.1 | 0.5 | Leiden | 0.7 | 0.5 | 2 | 13 | 0 | 0 |
| Kidney-mouse | 0.2 | 0.15 | 0.01 | 0.05 | $k$-means | 4 | 0.9 | 2 | 33 | 26 | 14 |
| Liver-brca-mouse | 0.2 | 0.2 | 0.1 | 0.05 | $k$-means | 4 | 0.8 | 2 | 12 | 6 | 3 |
| Lung-mouse | 0.2 | 0.2 | 0.1 | 0.05 | $k$-means | 4 | 0.8 | 2 | 20 | 10 | 4 |
| Ovary-ca-mouse | 0.2 | 0.2 | 0.1 | 0.05 | $k$-means | 4 | 0.8 | 2 | 24 | 12 | 6 |
| P3-intact | 0.35 | 0.1 | 0.1 | 0.05 | Leiden | 0.35 | 0.4 | 3 | 41 | 28 | 8 |
| P3-post-resection | 0.35 | 0.1 | 0.1 | 0.05 | Leiden | 0.35 | 0.4 | 3 | 48 | 27 | 14 |
| Mct-Cre | 0.35 | 0.1 | 0.1 | 0.05 | Leiden | 0.4 | 0.5 | 2 | 28 | 27 | 8 |
| Mct-nonCre | 0.35 | 0.1 | 0.1 | 0.05 | Leiden | 0.4 | 0.8 | 2 | 30 | 26 | 10 |
| Healty-brain-1 | 0.55 | 0.1 | 0.02 | 0.05 | Leiden | 0.2 | 0.7 | 2 | 13 | 8 | 0 |
| Healty-brain-2 | 0.4 | 0.1 | 0.1 | 0.05 | $k$-means | 5 | 0.7 | 2 | 20 | 14 | 11 |
| Healty-brain-3 | 0.4 | 0.1 | 0.1 | 0.05 | $k$-means | 5 | 0.7 | 2 | 33 | 20 | 14 |

*Abbreviations:* Off Prob: off-sample probability threshold. MSM: Metabolite Signal Matching threshold. $p_{adj}$: adjusted $p$-value for Moran's I (Benjamini-Hochberg correction). Param: clustering parameter ($k$ for $k$-means, resolution for Leiden). Fam: Ghost Families. Fam. Comp: Parameters for Fam computation. Tree Cut: dendrogram cut height (0-1). Min Size: minimum size for ghost families (e.g., $\geq 3$). Signif.: statistically significant families ($p_{adj} \leq 0.05$ and Moran's I $\geq 0.2$). See text for further details.

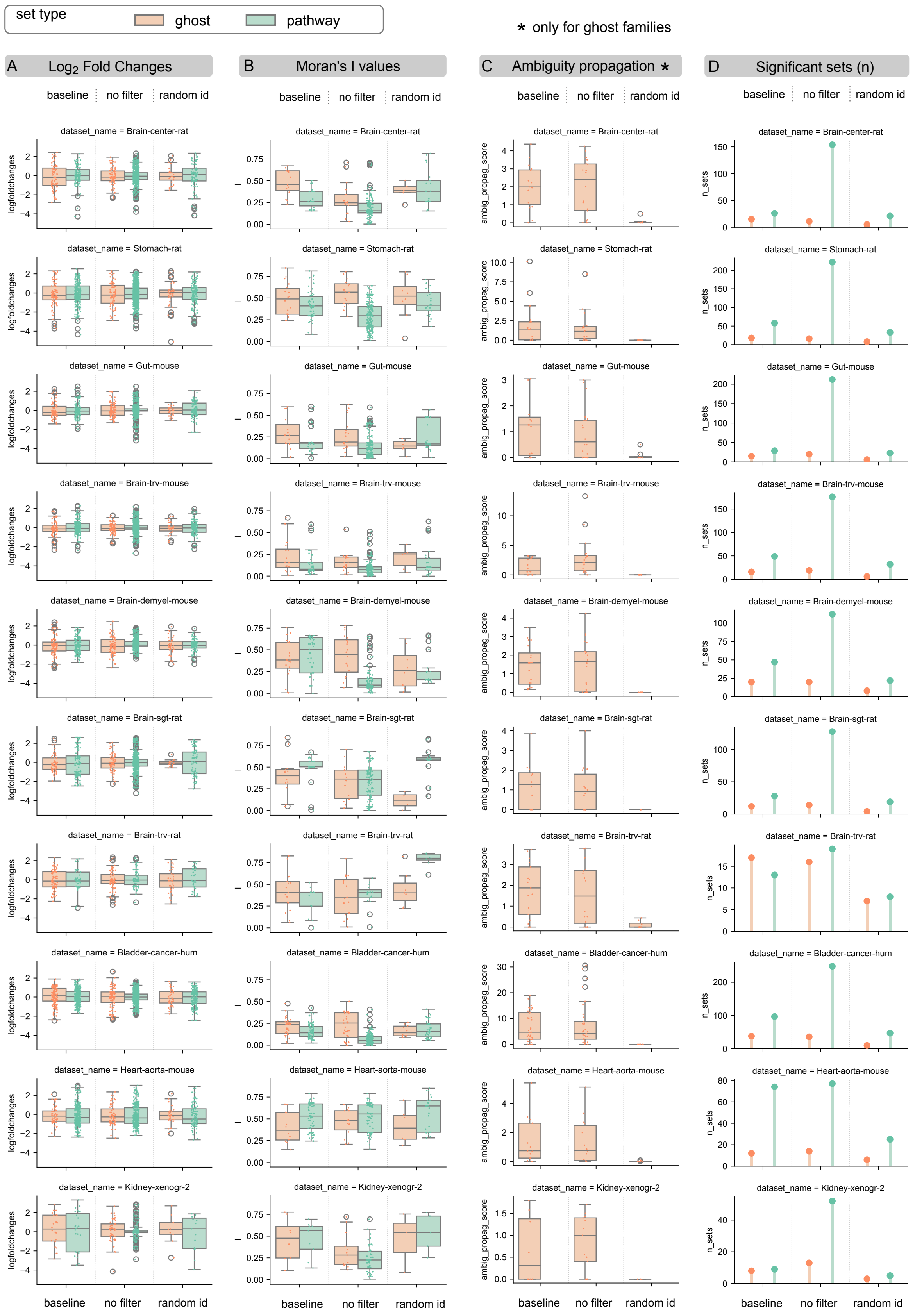

**Supplementary Fig. 2** **Systematic perturbation analysis of SpacePath.** Results for the first 10 datasets are shown for the baseline workflow and the no filter and random ID perturbations. Columns show: **A**, region-versus-rest effect sizes for molecular-set/region comparisons; **B**, Moran's $I$ of spot-wise molecular-set score maps; **C**, ambiguity-propagation scores for *de novo* molecular families; and **D**, the number of molecular sets passing the adjusted significance threshold ($P_{\mathrm{adj}} \leq 0.05$). Canonical pathways and *de novo* molecular families are displayed separately. Box plots show the median and interquartile range; points in A represent molecular-set/region comparisons, whereas points in B and C represent individual molecular sets.

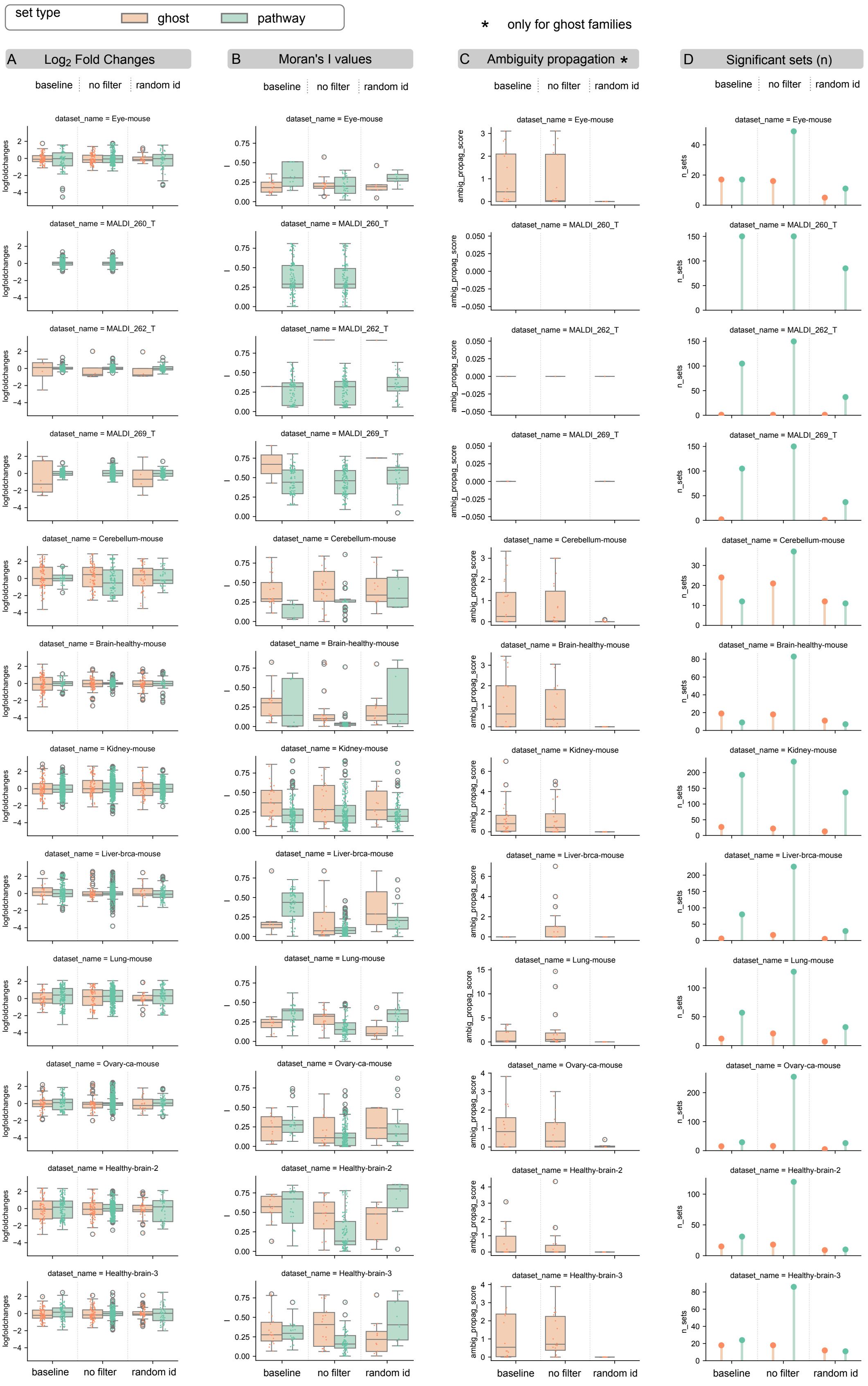

**Supplementary Fig. 3 Systematic perturbation analysis of SpacePath, continued.** Results for the remaining 12 datasets are displayed using the same metrics and conventions as in Supplementary Fig. 2. The three MALDI_260_T, MALDI_262_T and MALDI_269_T datasets correspond to small-molecule acquisitions and showed weaker performance across one or more configurations.

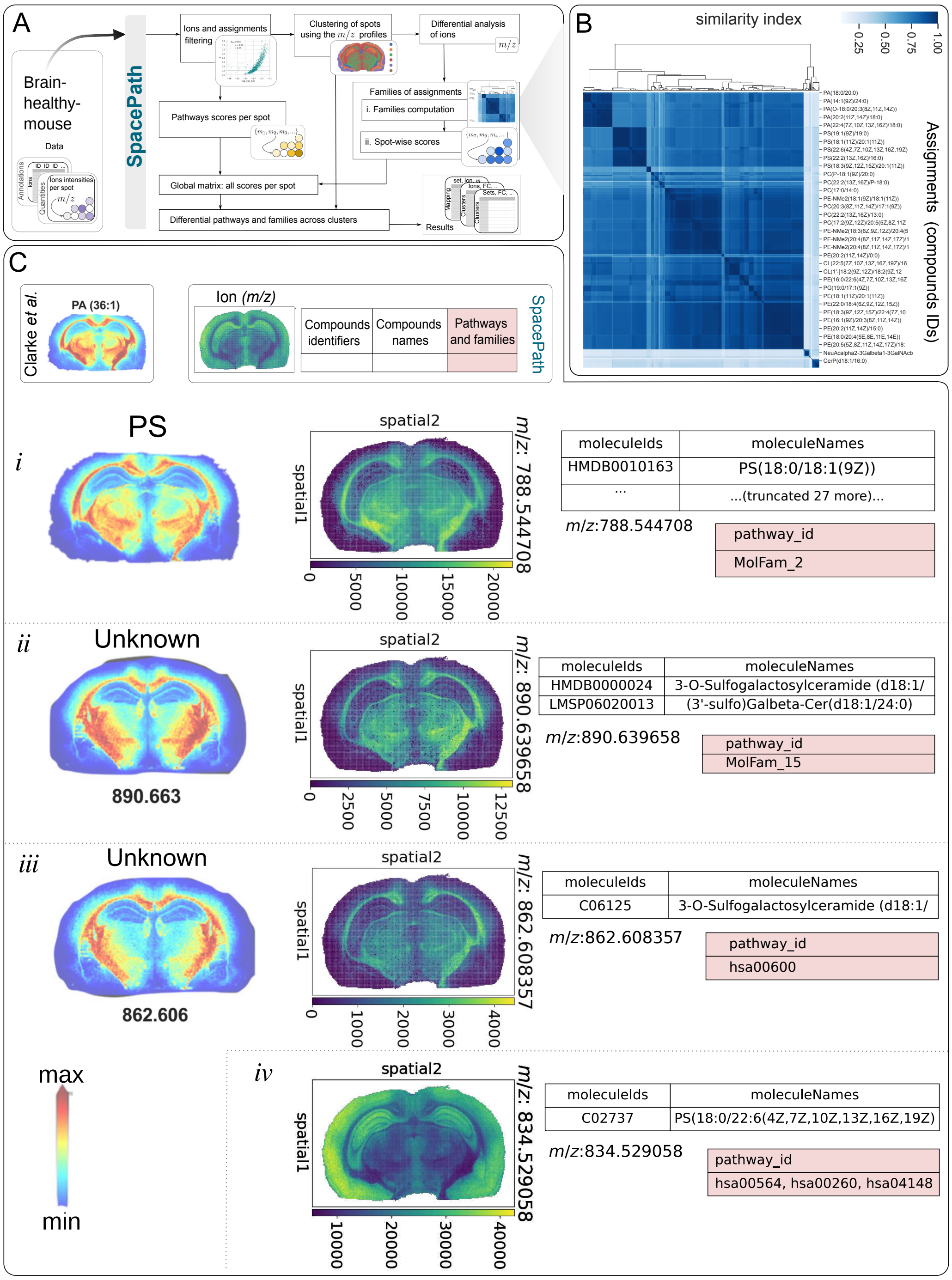

**Supplementary Fig. 4** A) SpacePath analysis of the healthy-mouse-brain lipidomics dataset from Clarke et al. B) Hierarchical organization of candidate molecular assignments as a result of the computation of *de novo* molecular families: for illustration purposes, a dendrogram is shown in the upper side of the Tanimoto similarities matrix, but such dendrogram is computed on the distances matrix. C) Direct comparison between selected lipid distributions reported by Clarke et al. and corresponding SpacePath results.

#### S3.3 Muscle-invasive bladder cancer

The muscle-invasive bladder cancer dataset from Tu et al. [7] was used to compare SpacePath with molecular distributions reported in the source study. Whereas Tu et al. primarily compared pooled tumor and non-tumor tissue, SpacePath analyzed spatial heterogeneity within a single annotated cancer section. The resulting regions and representative ion-level correspondences are shown in Supplementary Fig. 7.

Three ion-level observations provide additional detail beyond the main Results:

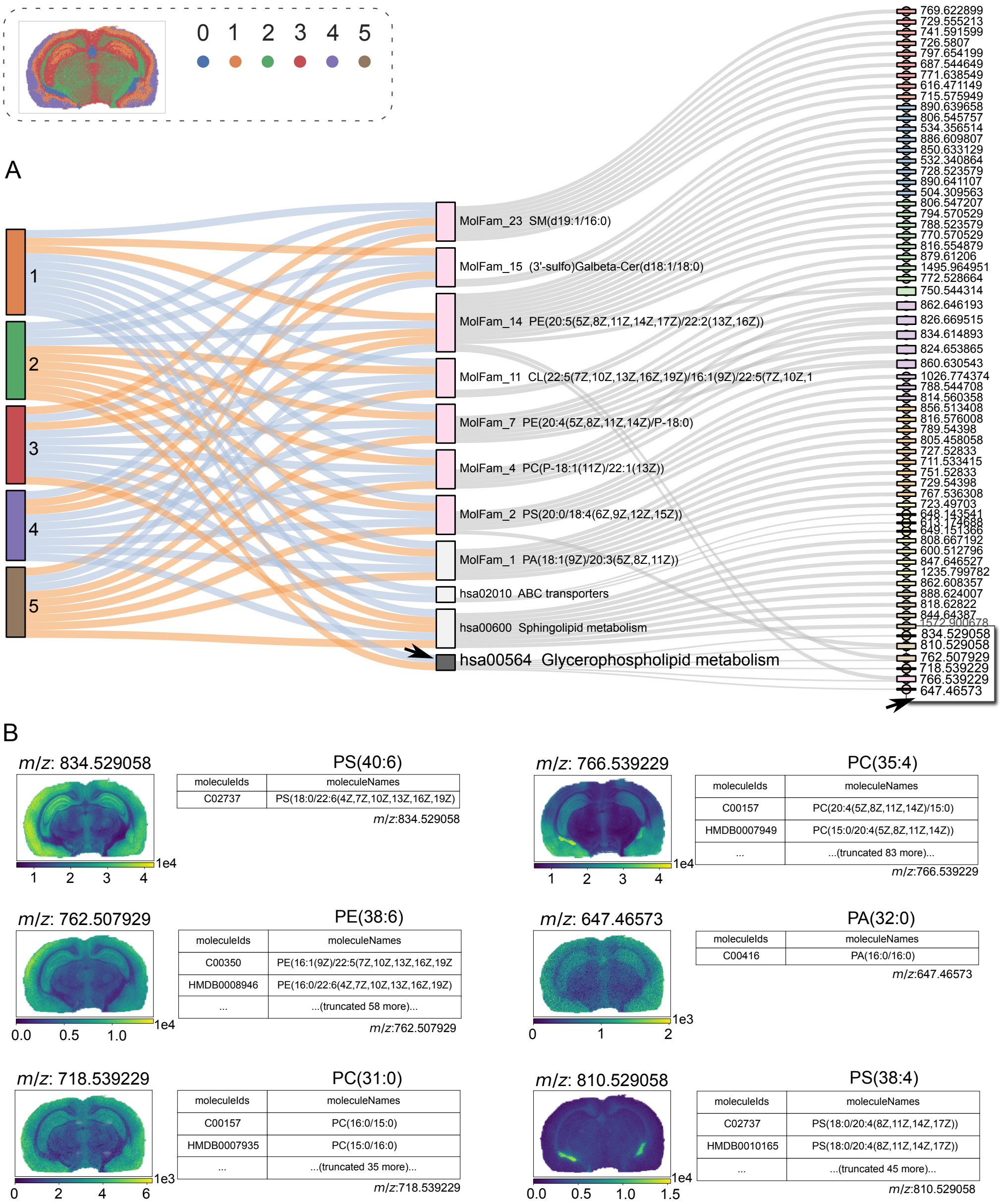

**Supplementary Fig. 5** Additional detail for the *Glycerophospholipid metabolism* pathway in the healthy mouse brain. A) Alluvial representation linking spatial regions, molecular sets, and selected contributing ions; the pathway and its contributing ions are indicated by arrows. B) Spatial distributions and candidate assignments of the six displayed ions contributing to this pathway.

- **PE(32:1)** showed preferential signal in the tumor-associated region (Supplementary Fig. 7C-i). Although this candidate compound maps to KEGG pathways, the corresponding molecular sets did not satisfy the SpacePath support requirements in this section.
- **PC(P-34:1)** likewise localized preferentially to tumor-associated tissue, consistent with the source study (Supplementary Fig. 7C-ii).
- **TG(54:3)** was concentrated in the smooth-muscle-associated region within the section (Supplementary Fig. 7C-iii).

Thus, individual ions can remain informative even when they do not contribute to a retained molecular set because fewer than two distinct ion-supported assignments are available.

The expanded analysis in Supplementary Fig. 8 provides several additional region-associated signals. A diacylglycerol-rich molecular family was associated with the tumor core, consistent with the increased DG/MG abundance reported by Tu et al. The smooth-muscle region showed higher *Fatty acid biosynthesis* scores, while a high-carbon-number triacylglycerol family (MolFam_11) containing

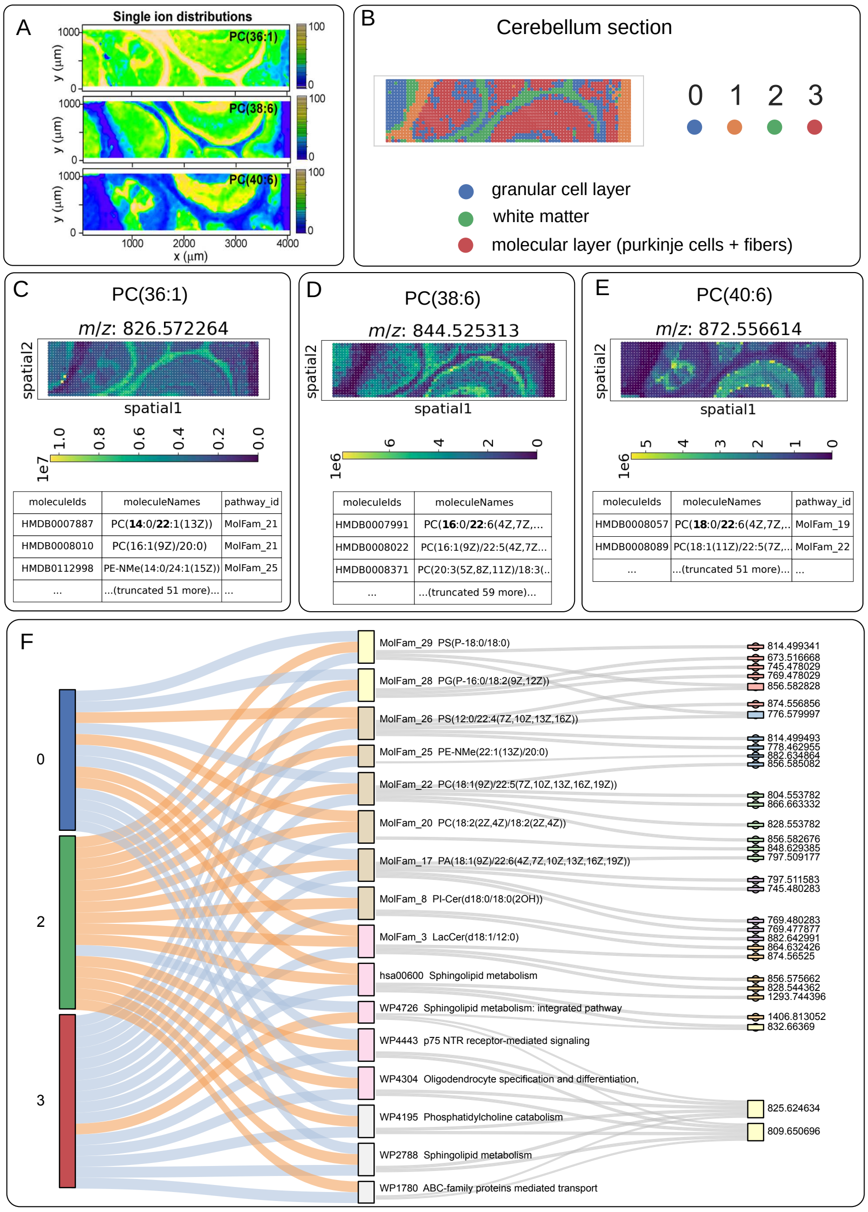

**Supplementary Fig. 6** Benchmarking on healthy mouse cerebellum. A) Original ion maps from Bond et al. B) Tissue segmentation by $k$-means ($k = 4$), including one non-tissue background cluster. C–E) Spatial distributions of phosphatidylcholine features recovered by SpacePath. F) Expanded alluvial representation linking spatial regions, significant canonical pathways and *de novo* molecular families, and contributing ions.

TG(58:7) was associated with peri-tumoral or non-tumor regions. These observations complement the representative fatty-acid and diglyceride signals reported in the main Results.

Additional canonical-set associations included wax/plasmalogen-related signals and the *Alpha-linolenic acid metabolism* and *Sebaleic acid metabolism* sets (Supplementary Fig. 8C). These associations were not reported in the source study and are therefore treated as dataset-specific candidate observations rather than as independently validated bladder-cancer programs.

### S4 Supplementary analysis of study-generated glioblastoma xenografts

The main Results report the principal SpacePath findings for the P3-intact and P3-post-resection xenografts. This section provides additional ion- and molecular-set associations for these tissues and presents the analysis of the two xenografts from the conditional *Mct1* cohort.

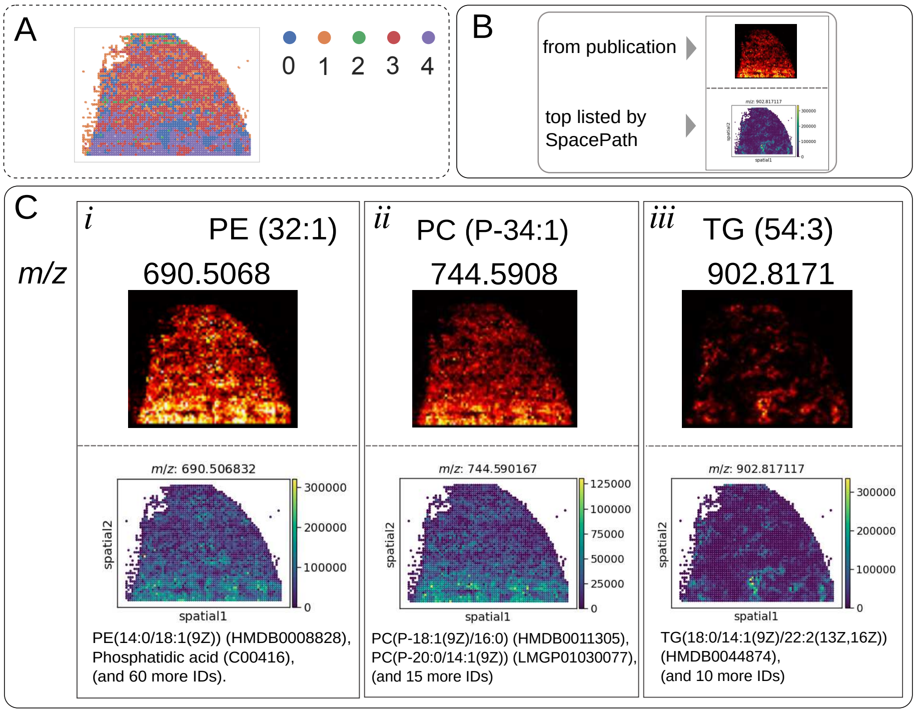

**Supplementary Fig. 7** Spatial distribution of selected ions in the bladder-cancer benchmark. A) Spatial regions identified in the analyzed tissue section. B) Legend for panel C. C) Ion distributions reported by Tu et al. (top) and corresponding features recovered by SpacePath (bottom).

**Supplementary Table 3 Spatial lipidomics datasets generated in this study.** Four glioblastoma xenograft tissues and three healthy mouse brain sections were acquired. Healthy-brain-1 was excluded from the cross-dataset analyses after application of the dataset-level inclusion criteria described in the main Methods. The four xenograft datasets are analyzed in this section.

| Dataset | Tissue | Experimental condition |
|---|---|---|
| P3-intact | brain | P3 xenograft, intact tumor |
| P3-post-resection | brain | P3 xenograft, post-resection |
| Mct1-Cre | brain | GL261 xenograft, $Mct1^{\mathrm{fl/fl;Cre/-}}$ |
| Mct1-nonCre | brain | GL261 xenograft, $Mct1^{\mathrm{fl/fl;nonCre}}$ |
| Healthy-brain-1 | brain section | healthy |
| Healthy-brain-2 | brain section | healthy |
| Healthy-brain-3 | brain section | healthy |

#### S4.1 P3 xenografts

##### S4.1.1 P3-intact xenograft

Supplementary Fig. 9 extends the P3-intact analysis shown in the main manuscript by displaying associations for all six spatial regions and additional representative ions.

Several additional ion-level patterns were not included in the main figure. The feature at $m/z$ 870.540963, assigned to PC(40:7) (HMDB0008090) and represented in MolFam_27, showed lower signal in the tumor and preferential localization to cortical and striatal regions (Supplementary Fig. 9B). LysoPE(20:4), represented in MolFam_36, was associated with the pia-mater region and also detected in the striatal compartment, whereas a glucosylceramide-associated signal localized preferentially to white matter.

The feature at $m/z$ 725.556796, assigned to SM(d34:1), showed higher signal in the tumor and mapped to the WikiPathways set *Sphingolipid metabolism integrated pathway* (WP4726). This association is not displayed in the alluvial summary because only a subset of statistically ranked overlapping sets is shown. The example illustrates that one candidate compound can contribute to several overlapping

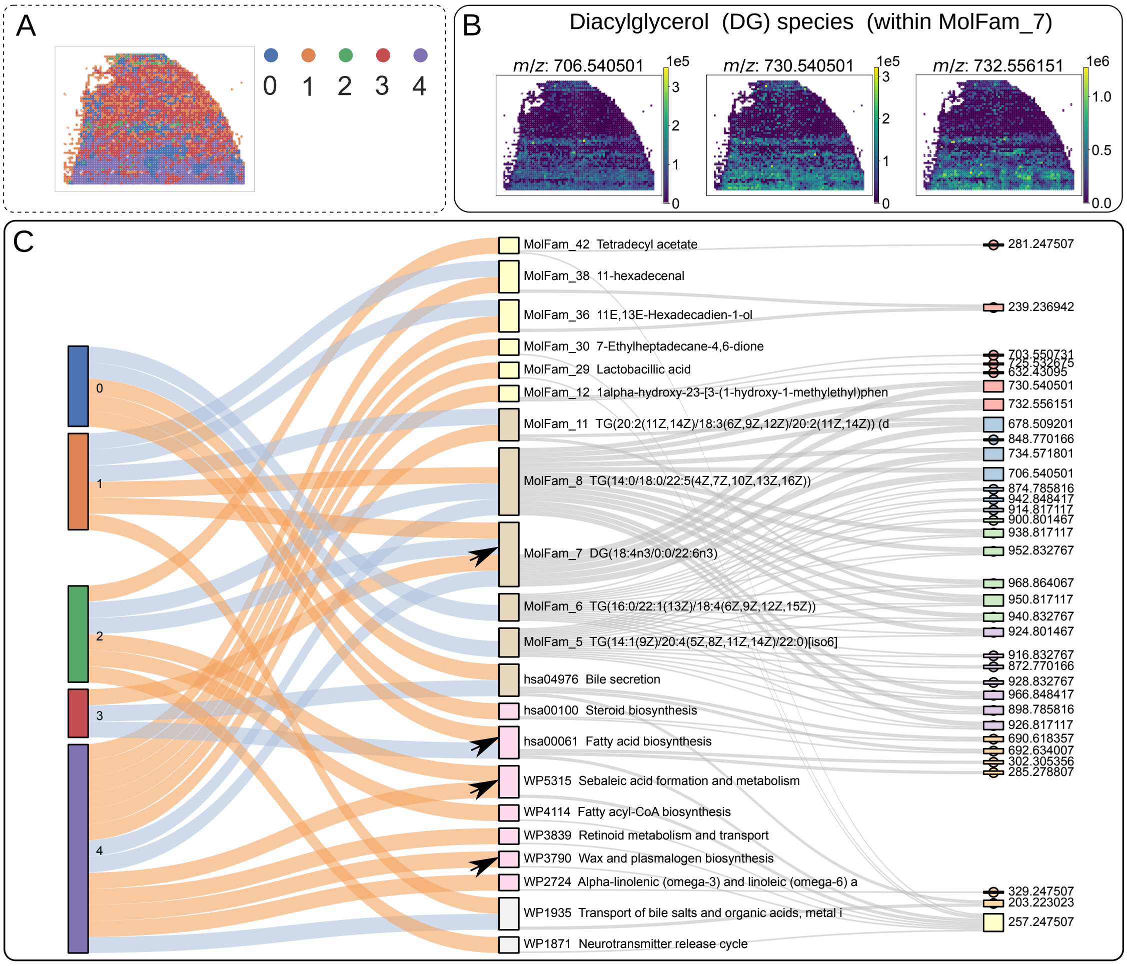

**Supplementary Fig. 8** Additional molecular-set results for the bladder-cancer benchmark. A) Spatial regions. B) Representative diacylglycerol distributions in tumor-associated tissue. C) Alluvial summary linking regions, canonical pathways and *de novo* molecular families, and contributing ions. Directionality labels in the original visualization represent relative within-section score differences and should not be interpreted as direct measurements of pathway activity. Arrows indicate associations discussed in the text.

canonical sets and that the alluvial representation is a visualization subset rather than an exhaustive pathway inventory.

SpacePath also identified tumor-associated ions whose corresponding molecular families did not reach the molecular-set significance threshold. In particular, $m/z$ 669.446577, assigned to PA(32:1) and represented in MolFam_33, was significant in the ion-level region-versus-rest analysis despite MolFam_33 itself not passing the set-level threshold (Supplementary Fig. 9C). This provides an example in which a spatially informative ion is retained even when its higher-level molecular set is not significantly differentiated.

A second tumor-localized feature, $m/z$ 854.590561, was assigned by METASPACE to PS(41:4) (LMGP03010478) and represented in MolFam_16 (Supplementary Fig. 9C). The odd total carbon number of this candidate assignment warrants cautious interpretation, and no alternative even-carbon candidate was available in the annotation used here. The spatial association of the ion is therefore distinguishable from certainty about its molecular identity, which would require orthogonal structural evidence.

##### S4.1.2 P3-post-resection xenograft

The main manuscript reports the principal spatial programs in the P3-post-resection section. Supplementary Fig. 10 provides the complete alluvial representation and additional ion-level associations across all seven spatial regions.

A phosphatidylcholine-rich molecular family (MolFam_23) was preferentially associated with gray-matter regions (clusters 4–6; Supplementary Fig. 10B). Additional sphingomyelin-associated signals

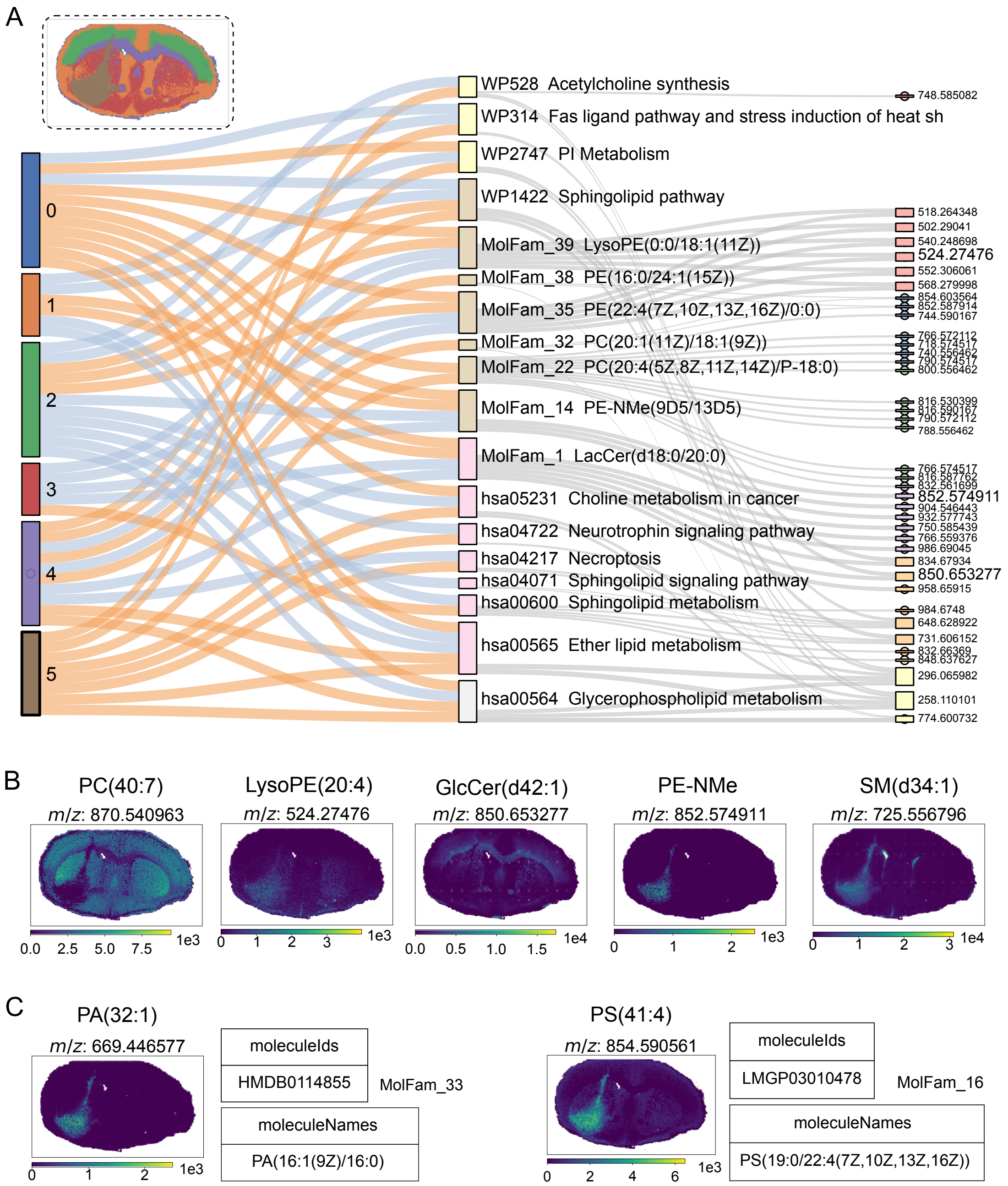

**Supplementary Fig. 9 Additional SpacePath results for the P3-intact glioblastoma xenograft.** A) Spatial regions and alluvial representation linking regions, canonical pathways and *de novo* molecular families, and contributing ions. All six spatial regions are included. B) Spatial distributions of additional representative ions. C) Examples of ion-level output combining candidate METASPACE assignments with canonical-pathway or molecular-family membership.

were observed in the recurrent-tumor region. For example, SM(d42:2) contributed to the canonical *Sphingolipid metabolism integrated pathway* (WP4726).

The tumor-associated SM(d34:1) assignment provides an additional example of overlapping canonical-set membership: the same candidate compound was represented in *Phosphatidylcholine catabolism* (WP4195), *Oligodendrocyte specification and differentiation* (WP4304), *ABC-family proteins mediated transport* (WP1780), and *Sphingolipid metabolism integrated pathway* (WP4726) (Supplementary Fig. 10B). These multiple memberships reflect overlap among pathway definitions and should not be interpreted as four independent biological signals.

The feature at $m/z$ 854.590561 was again associated with the tumor region and carried the PS(41:4) candidate assignment discussed above (Supplementary Fig. 10C). Its recurrence across the two P3

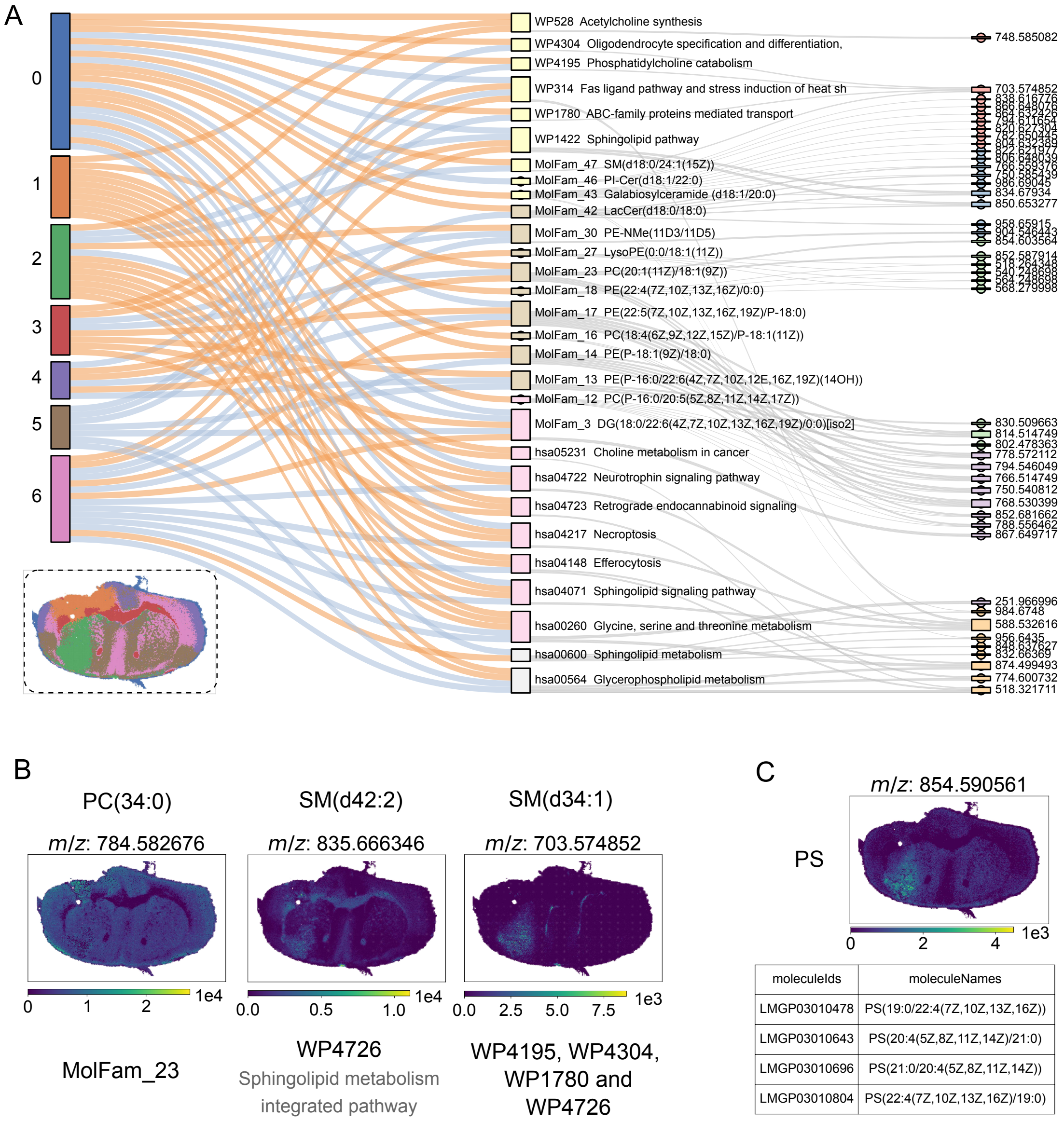

**Supplementary Fig. 10 Additional SpacePath results for the P3-post-resection glioblastoma xenograft.** A) Complete alluvial representation linking spatial regions, canonical pathways and *de novo* molecular families, and contributing ions. All seven spatial regions are included; the region-color key is shown below. B) Spatial distributions of additional representative ions. C) Example of a candidate molecular assignment requiring cautious interpretation.

sections supports the spatial reproducibility of the ion-level observation but does not resolve the molecular identity of the feature.

#### S4.2 Second cohort: GL261 xenografts in Mct1-Cre and Mct1-nonCre mice

The second cohort comprised two GL261 xenografts, one generated in an *Mct1*$^{\mathrm{fl/fl;Cre/-}}$ mouse and one in an *Mct1*$^{\mathrm{fl/fl;nonCre}}$ mouse (Supplementary Table 3). SpacePath was applied independently to both sections.

In both sections, the tumor-associated region was differentiated from surrounding brain by several molecular-set scores, including *Sphingolipid metabolism*, *Efferocytosis*, and the *Fas ligand pathway* (Supplementary Fig. 11). These observations are within-section spatial associations.

The MSI acquisition used a lateral resolution of 25 µm, which does not isolate macrophage-specific signals. The two sections are retained as additional examples of SpacePath analysis across distinct xenograft backgrounds, but no macrophage-specific mechanistic conclusion is drawn from them.

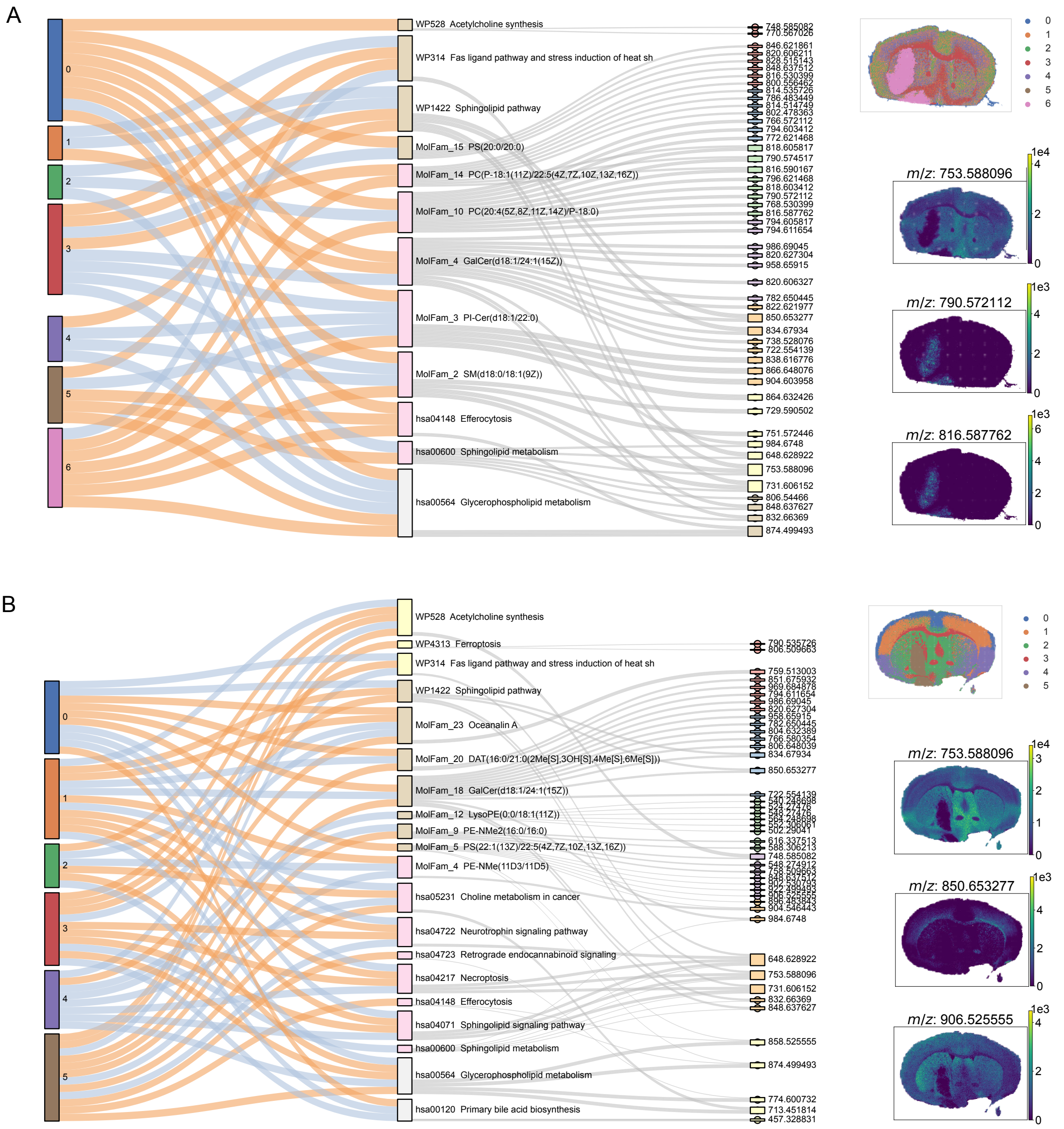

**Supplementary Fig. 11 SpacePath analysis of the second glioblastoma xenograft cohort.** Global SpacePath outputs are shown for the two GL261 xenograft sections, with all spatial regions included. A) Mct1-Cre. B) Mct1-nonCre.
